# A population-scale Red Blood Cell proteome atlas of 13,000 donors uncovers genetically encoded aging clocks predicting hemolysis, transfusion efficacy, and donor activity a decade later

**DOI:** 10.64898/2026.03.07.710284

**Authors:** Monika Dzieciatkowska, Aaron V. Issaian, Gregory R. Keele, Anthony Saviola, Daniel Stephenson, Shaun Bevers, Julie A Reisz, Zachary B. Haiman, Travis Nemkov, Fang Fang, Amy L Moore, Xutao Deng, Mars Stone, Steve Kleinman, Philip J. Norris, Xunde Wang, Swee-Lay Thein, Eldad A. Hod, Michael P. Busch, Nareg H. Roubinian, Grier P Page, Kirk C. Hansen, Angelo D’Alessandro

## Abstract

As the most abundant human cell and the foundation of transfusion medicine, red blood cells (RBCs) offer a unique readout of systemic health, yet they have never been characterized at population scale. We generated a proteome atlas of 13,091 blood donors with multi-omics longitudinal phenotyping, characterizing the influence of demographics and genetic variation on the reproducibility of RBC proteomes across donations. Elastic-net aging clocks captured biological aging with high accuracy and uncovered genetic regulators of ΔAge at FN1, C4/IKZF1, CRAT, PFAS, TRIM58. Across independent cohorts, ΔAge was accelerated in G6PD deficiency, sickle cell trait/disease, and iron deficiency, reversed by iron repletion, and slowed in high-frequency donors, linking molecular aging to brain iron/myelin and cognitive performance. Molecular aging signatures predicted storage, osmotic, and oxidative hemolysis, hemoglobin increments after transfusion, and long-term donor activity over 12-years. These results establish RBC proteomics as a scalable biomarker of aging, donor healthspan, and transfusion outcomes.

Dzieciatkowska *et al.* generate the first population-scale atlas of the RBC proteome across 13,000 donors and develop proteomic and metabolomic aging clocks that quantify biological age. Molecular ΔAge is reproducible across donations, genetically encoded and accelerated in G6PD deficiency, sickle cell trait/disease, and iron deficiency - yet reset by iron repletion, tracking with cognitive function and brain iron/myelin. RBC aging clocks predict hemolytic fragility, transfusion efficacy, and donor activity 12 years later.

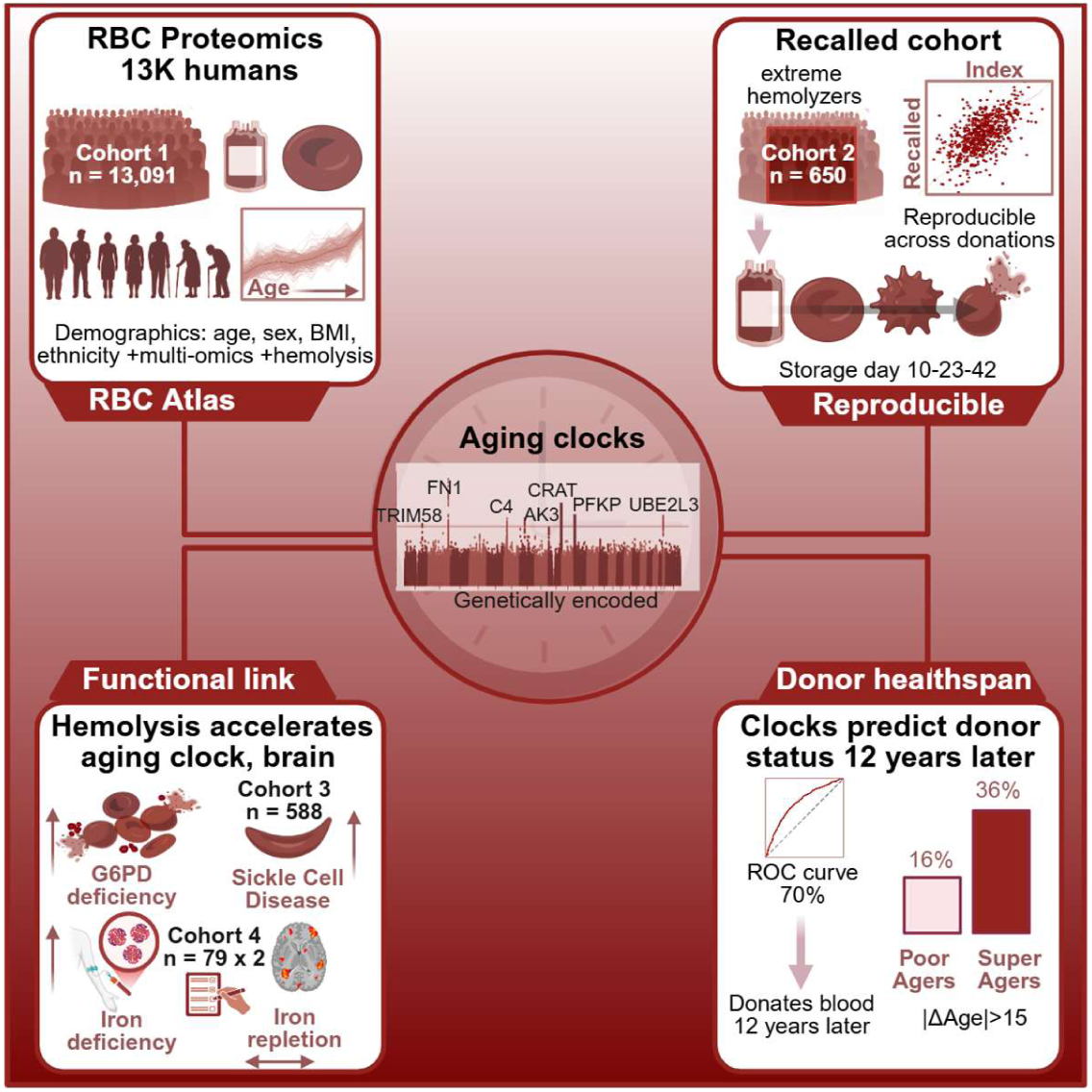

**Highlights:**

- RBC proteome atlas of 13,091 donors reveals demographic and genetic programs
- Genetically encoded RBC aging clocks identify regulators of molecular Δage
- Molecular aging features predict hemolysis and transfusion response across cohorts
- RBC molecular age forecasts long-term donor activity over a 12-year follow-up

## INTRODUCTION

Red blood cells (RBCs) are the most abundant human cell type - constituting ∼83% of all 30 trillion cells in an adult body^1,2^ - and the foundation of modern transfusion medicine, with ∼118.5 million units administered globally each year. For decades, RBCs were viewed as biochemically simple: enucleated cells lacking mitochondria and protein synthesis, with ∼90% of the proteome dominated by hemoglobin and limited metabolic plasticity. Classic biochemical studies reinforced this perception, and the RBC proteome was long considered essentially complete.^3^

Advances in mass spectrometry overturned this assumption. High-resolution proteomics now identifies more than one thousand RBC proteins^4–6^, revealing a complex and dynamic molecular landscape^7^. Although mature RBCs cannot synthesize new proteins, the proteome inherited from erythroid precursors is increasingly recognized as sensitive to aging^8^, sex^9^, genetic variation^10^, stress erythropoiesis^11^, labile iron pools^12^, inflammation^13,14^, high altitude^15^, exercise, and underlying pathology, including kidney disease, hemoglobinopathies and enzymopathies such as sickle cell disease (SCD)^16^ or glucose-6-phosphate dehydrogenase (G6PD) deficiency^9^ - as well as storage under routine blood bank conditions. ^17^

In this view, RBCs function as a circulating organ, ∼2.5 kg in mass, whose proteome and metabolome reflect systemic (patho)physiology as cells complete a full circuit every minute, up to ∼178,000 times over their 120-day lifespan. This principle already underlies clinical assays such as hemoglobin A1c, where long-lived RBC proteome modifications serve as integrative readouts of chronic glycemic exposure. Yet despite these advances, no study has characterized the RBC proteome at true population scale, nor systematically mapped how genetics, demographics, and physiology shape RBC molecular diversity across thousands of individuals.

Population-scale proteomics^18^ in humans has largely been driven by affinity-based platforms - including antibody-based proximity extension assays^19^ and aptamer-based technologies^20^, which have enabled deep proteogenomic studies in tens of thousands of participants^21–25^. However, these assays are optimized to quantify circulating plasma proteins or other proteomes not affected by the overwhelming abundance of a single class of protein like hemoglobins, and are fundamentally incompatible with RBC biology. Mature RBCs are dominated by extremely abundant intracellular proteins that overwhelm affinity-based assays, generate cross-reactivity artifacts, and obscure low-abundance targets. In addition, extensive oxidative and enzymatic post-translational modifications alter epitopes and disrupt both antibody and aptamer binding. As a result, affinity proteomics cannot reliably profile the intracellular RBC proteome, necessitating mass spectrometry–based approaches. In parallel, biological aging clocks have transformed aging research. The first epigenetic clocks - Horvath’s pan-tissue clock^26^ and the Hannum blood clock^27^ demonstrated that DNA methylation patterns quantify biological age more accurately than chronological age. Subsequent models such as epigenetic DNA methylation-based PhenoAge^28^ and GrimAge^29^ predict morbidity, mortality, and physiological decline, establishing aging clocks as powerful biomarkers. Multi-omic aging clocks derived from circulating cell-free mitochondrial DNA and imaging have since emerged^30,31^, yet no aging clock has been developed for RBCs despite their systemic circulation, metabolic specialization, and sensitivity to genetic, nutritional, and oxidative stress.

Mass-spectrometric profiling of RBCs has advanced rapidly.^7^. Still, most RBC proteomics studies have involved fewer than a hundred subjects, limiting generalizability. The national Recipient Epidemiology and Donor Evaluation Study (REDS) programs,^32,33^ with harmonized workflows across four U.S. blood centers, provide an unprecedented opportunity for population-scale RBC omics through highly standardized collection, additive solutions, and processing pipelines.

## RESULTS

### RBC proteomics of 13,091 donors defines the largest atlas of RBC protein variation to date

In the present study we leveraged a mass spectrometry-based DIA-PASEF workflow to generate a RBC proteome atlas of samples from blood units donated by 13,091 index volunteers enrolled in the REDS RBC Omics study (**Figure 1A**). Donors spanned broad and representative demographic distributions in age (18–90, median 47), sex (50.04% female), ethnicity (62% Caucasian, 14% Asian, 14% African American, 10% Hispanic), and BMI across four U.S. blood centers (**Figure 1B**), providing statistical power to dissect how demographic, genetic, and processing variables jointly shape red cell biology at population scale. Mass Spectrometry-based proteomics was performed on 142 96–well plates, using multiplexed S-Trap digestion and rapid DIA-PASEF acquisition on a timsTOF instrument, searched against a pooled spectral library encompassing all donors (**Supplementary Figure 1A**). Despite the overwhelming abundance of hemoglobin and the high-throughput workflow (5.6 min/sample), we quantified 956 proteins (**Figure 1C; Supplementary Table 1**). To our knowledge, this resource constitutes the largest RBC proteomics dataset assembled to date.

**Figure 1.**
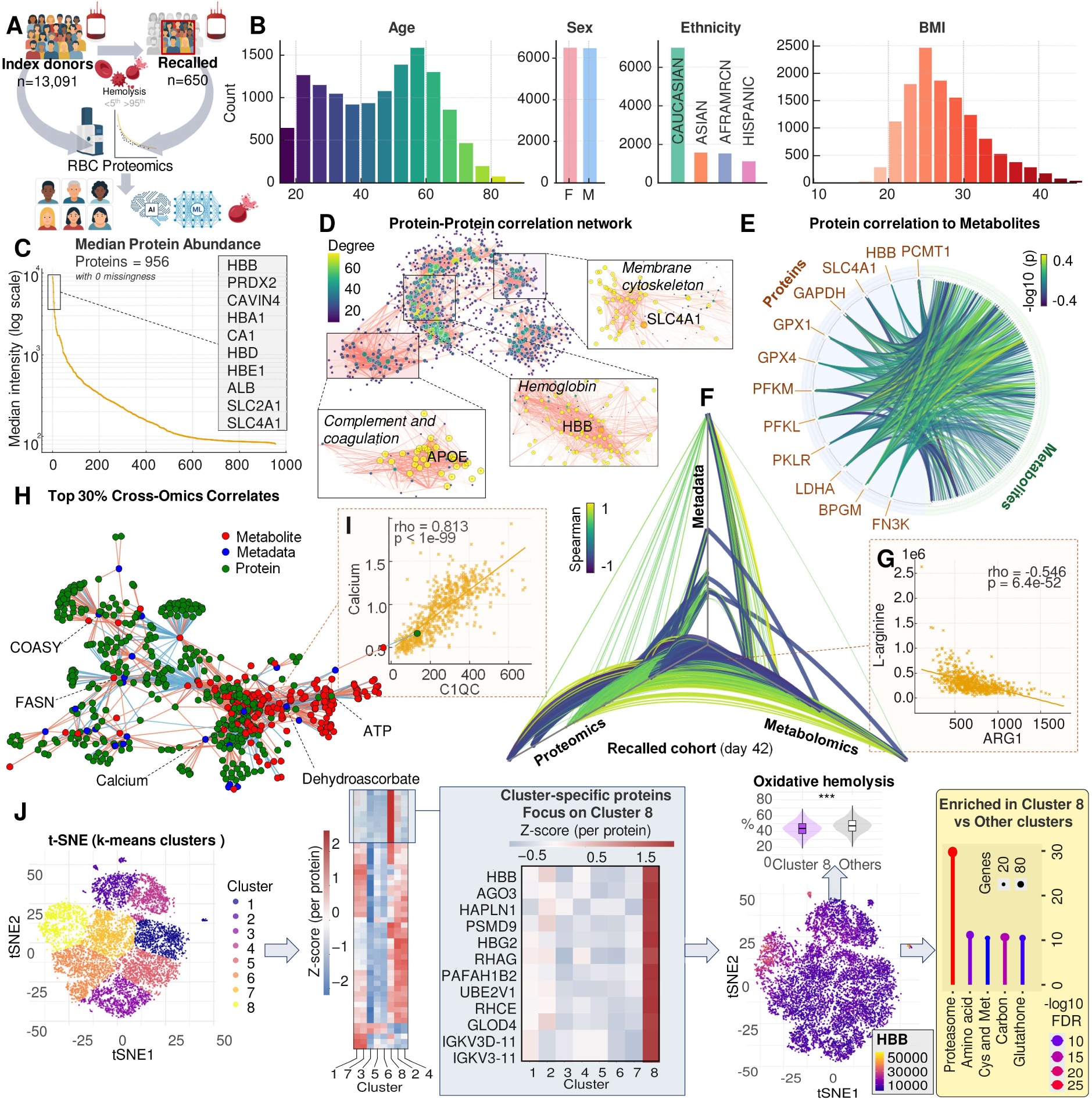
Multi-omics profiling of 13,091 blood donors reveals coordinated proteomic and metabolic structure across RBC biology, donor demographics and hematological parameters. Overview of the study design integrating RBC proteomics and metabolomics from 13,091 index donors, with deep phenotyping of a validation cohort (n=650 recalled donors from the index cohort), including hemolysis measures, metadata, and metabolomics **(A).** Distribution of donor demographics (Age, Sex, Ethnicity, and BMI) across index donors **(B).** Despite the overwhelming abundance of hemoglobin in mature RBCs, proteomics analyses quantified 956 proteins with 0 missing data across all samples, ordered by decreasing intensity in **(C).** Protein–protein correlation network (Spearman), visualized by degree centrality, identifying major functional neighborhoods: membrane cytoskeleton (e.g., SLC4A1), hemoglobin cluster (e.g., HBB), complement/coagulation (e.g., APOE), and redox/metabolic nodes **(D)**. Chord diagram showing the most significant (top 25% by Spearman rho, filtered by adjusted p-values) protein-metabolite correlations, highlighting coordinated variation linking glycolytic enzymes (GAPDH, PFKM, LDHA), hemoglobin subunits, and antioxidant enzymes (GPX1) to metabolic outputs **(E).** Hive plot summarizing correlation structures between metadata (complete blood count, ferritin levels, hemolysis parameters), proteomics, and metabolomics in recalled donors (day 42), demonstrating strong cross-layer connectivity, particularly between metabolites and clinical metadata **(F).** Representative examples of protein–metabolite associations: C1QC correlating positively with Calcium (ρ = 0.813), and ARG1 correlating negatively with L-arginine (ρ = –0.546) **(G).** Cross-omics and metadata network of the top 30% strongest correlates in recalled donors, showing hubs centered on metabolic pathways including ATP turnover, fatty acid synthesis (FASN), and redox metabolism (e.g., dehydroascorbate) **(H).** Scatter plots for selected high-confidence protein–metabolite relationships (expanded from panel H). t-SNE visualization of donors clustered by k-means (k = 8, selected using elbow, silhouette, and gap-statistic analyses to determine the optimal cluster structure), followed by identification of cluster-specific proteins and functional enrichment. Cluster 8 shows pronounced elevation of hemoglobin-associated proteins (HBB, HBA1, HBD) and is enriched for oxidative hemolysis signatures, proteasome components, and antioxidant systems relative to other clusters **(J).**

### Technical quality control and removal of plate-dependent structure

A comprehensive set of quality control analyses confirmed the technical robustness of the dataset. Principal component analysis (PCA) of raw intensities showed modest plate-dependent structure explaining approximately 3.2% of total variance (PC-weighted variance decomposition of the raw dataset in **Supplementary Figure 1B–C**), which was nearly eliminated after plate-centering normalization (**Supplementary Figure 1D**). Run-order dependence of the internal standard yeast ADH1 was abolished by this correction, and no protein retained injection-order correlations exceeding 0.04 (**Supplementary Figure 1E**). Spatial analyses across plate geometry demonstrated minimal row, column, or edge effects (R² = 0.02; Cohen’s d = 0.08; row p = 0.94; column p = 0.41; **Supplementary Figure 1F–H**). These results confirmed that technical artifacts had been effectively minimized and motivated explicit adjustment for plate in all downstream modeling.

### Population-scale architecture of the RBC proteome and cross-omics correlates

With technical influences minimized, the proteomic space resolved into its underlying biological dimensions. Protein–protein correlation networks based on Spearman associations delineated distinct functional neighborhoods, including a membrane-cytoskeleton module centered on band 3 (SLC4A1) and ankyrin; a dense hemoglobin cluster dominated by HBB and HBD; complement/coagulation and lipid-associated communities featuring APOE and fibrinogen chains; and redox/metabolic modules linking glycolytic enzymes with glutathione-dependent antioxidants (**Figure 1D**). These structured relationships illustrate that even in a terminally differentiated, nucleus-free cell, the RBC proteome is not a monotonous hemoglobin background but a highly coordinated ensemble of interacting pathways with functionally relevant implications for blood bank storage quality and transfusion efficacy.

Building on our prior genomics^34^ and metabolomics work in this same population^35,36^, the present dataset advances from a metabolism-centric view to a densely connected multi-omic atlas integrating proteomics, metabolomics, donor demographics, hemolysis phenotypes, and clinical outcomes from transfusion recipients linked via a vein-to-vein database. To establish cross-layer coherence, we integrated the proteomic atlas with matched metabolomics from the same donors. A chord diagram of the strongest correlations highlighted dense biochemical cross-talk between glycolysis (GAPDH, PFKM, LDHA), hemoglobin subunits, glutathione-dependent antioxidants such as GPX1, and metabolic intermediates involved in energy and redox balance (**Figure 1E**).

An additional strength of this study is the availability of a second, independent multi-omic dataset in a recalled sub-cohort of 650 donors, who returned 6–18 months after their index donation for collection of a new RBC unit. Recall criteria were derived from the index dataset and included extreme hemolytic propensity - defined as falling below the 5th or above the 95th percentile for spontaneous storage hemolysis or hemolysis after osmotic or oxidative stress - or designation as a high-frequency donor with more than three donations in the preceding 12 months (**Figure 1A**). For this recalled unit, proteomics, metabolomics, and hemolysis metrics were obtained at days 10, 23, and 42 of storage, providing a longitudinal, intra-donor view of RBC remodeling that complements the cross-sectional atlas.

In this recalled cohort, a hive plot integrating proteomics, metabolomics, CBC indices, ferritin, and hemolysis parameters revealed extensive cross-layer connectivity, with metabolite levels showing particularly tight alignment with hematologic and hemolytic traits (**Figure 1F**). Representative protein–metabolite associations further illustrated this specificity: C1QC tracked positively with calcium, consistent with complement–ion homeostasis coupling, whereas ARG1 inversely tracked with L-arginine, reflecting substrate depletion by arginase (**Figure 1G**). A correlation network comprising the top 30% cross-omic associations identified hubs centered on ATP turnover, fatty-acid synthesis, redox metabolism, and intracellular calcium flux, a hallmark of erythrocyte-specific non-apoptotic cell death or eryptosis^37^ (**Figure 1H–I**). Together, these data establish a strongly interlocked proteomic–metabolomic atlas that captures RBC biology as a coordinated systems-level network.

### Unsupervised proteomic clustering reveals a “super-donor–like” molecular state shaped by both biology and processing pipelines

To examine how RBC proteomes varied across the 13,091 donors once plate-related technical influences had been minimized, we applied t-SNE embeddings followed by k-means clustering, selecting k = 8 based on elbow, silhouette, and gap-statistic criteria (**Figure 1J**). This analysis resolved the donor population into discrete, biologically coherent proteomic states rather than a continuous spectrum. Heatmaps of cluster-defining proteins showed reproducible enrichment patterns involving oxidative stress regulators, cytoskeletal proteins, proteasome components, metabolic enzymes, and inflammatory markers (**Supplementary Figure 2A**). Spatial projection of individual protein intensities across the t-SNE map confirmed that these clusters represented true molecular neighborhoods rather than artifacts of dimensionality reduction (**Supplementary Figure 2B–D**).

Among these clusters, one stood out prominently. Cluster 8 exhibited a proteomic signature marked by elevated hemoglobin subunits (HBB, HBD, HBG2), high abundance of proteasome and ubiquitin pathway components, and increased glutathione- and thioredoxin-linked antioxidant enzymes, combined with lower levels of complement proteins, immunoglobulin fragments, and lipid-binding proteins (**Supplementary Figure 2C**, **Supplementary Figure 3B**). This constellation of features mirrors the proteomic hallmarks of RBCs with superior storage quality and post-transfusion performance described in our prior work.^38^ Consistently, RBCs from donors in this cluster showed significantly lower hemolysis (**Figure 1J**).

Importantly, the distribution of donors contributing to Cluster 8 was not uniform across the cohort. While demographic factors (age, sex, BMI) showed minimal differences, and ethnicity differences were modest (**Supplementary Figure 3A**), processing variables displayed striking structure. Cluster 8 was disproportionately enriched among donors from a single blood center (Blood Systems Research Institute – BSRI, originally Blood Center of Pacific and recently rebranded as Vitalant) and among units preserved in the CP2D;AS-3 additive, highlighting that center-specific recruitment patterns and processing pipelines can produce recognizable molecular strata within ostensibly biological clusters (**Supplementary Figure 3C–D,F**).

The enrichment of Cluster 8 within specific centers prompted a broader examination of center-level and additive-related influences across the entire cohort. Donor recruitment, blood collection, and pre-processing of both index and recalled cohorts occurred at four geographically distinct U.S. blood centers (American Red Cross – ARC; Blood Center of Wisconsin – BCW; BSRI; Institute for Transfusion Medicine – ItxM). Center-specific pre-processing explained approximately 7% of the total variance (PC-weighted decomposition; **Supplementary Figure 1B**), largely reflecting the use of different FDA-approved storage additives (CPD;AS-1 at BCW/ItxM versus CP2D;AS-3 at ARC/BSRI). Additive differences accounted for less than 2% of the total proteome variance. All blood units were platelet and leukocyte filtered (log2.5 and log4, respectively), yielding fewer than one million WBCs per bag against ∼2–2.5 trillion RBCs, while modern filters allow <10 mL residual plasma per 500 mL blood unit, offering an opportunity to detect small yet quantifiable plasma-derived protein variation. Broader comparisons across blood centers revealed systematic differences in donor composition, ferritin levels, hemolysis metrics, and additive use, further reinforcing the need to explicitly account for these variables (**Supplementary Figure 3E,G**). Proteome-wide contrasts between CPD;AS-1 and CP2D;AS-3 confirmed that additive solution itself was associated with broad molecular shifts (**Supplementary Figure 3H**).

**Figure 2A** summarizes the mean proportion of variance (mean R^2^) explained across all 956 quantified proteins, systematically ranking the contribution of each metadata variable. Altogether, these analyses confirm that donor-intrinsic factors, such as age, sex, BMI and ethnicity dominate proteome-wide variability, while processing variables - including plate, additive solution, and blood center - exert measurable, structured effects that will be accounted for in downstream analyses.

**Figure 2.**
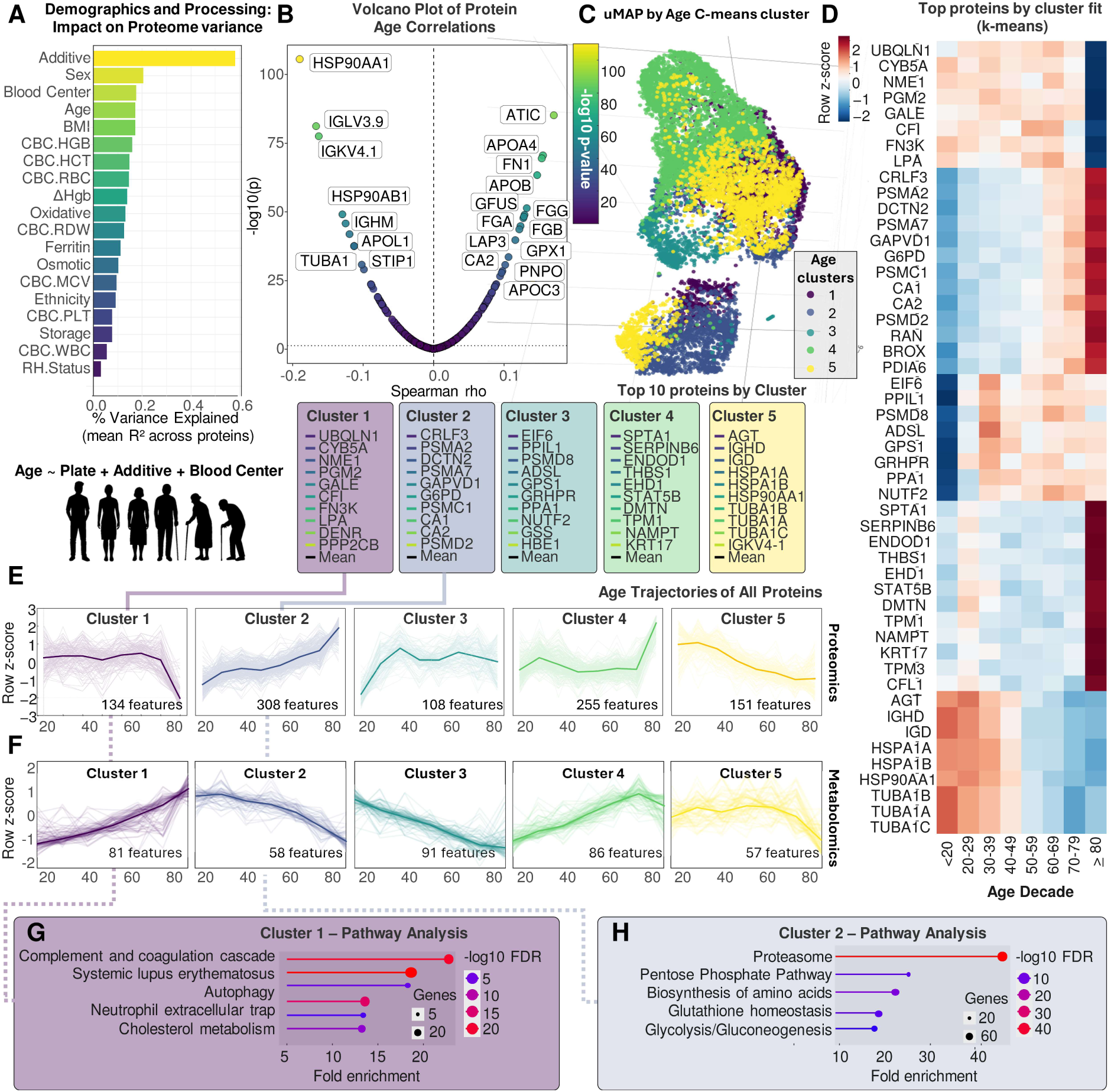
Age shapes the RBC proteome and metabolome through coordinated multi-omic remodeling across demographic strata. Variance partitioning analysis showing the relative contribution of donor demographics and processing variables to proteomic variability **(A)**. Processing (additive solution, and blood center) and demographics (Sex, Age, BMI) account for the largest proportion of explained variance across proteins. Volcano plot of protein correlations with Age, highlighting significantly age-associated proteins (Spearman, FDR-adjusted, adjusted by additive and blood center) **(B)**. uMAP of the proteome colored by Age-derived fuzzy c-means clusters (c = 6) **(C)**. Heatmap showing cluster-specific top proteins across age decades, illustrating progressive remodeling of cytoskeletal, metabolic, chaperone, and antioxidant pathways with advancing age **(D)**. Age trajectories of proteomic features (row-Z-scored), grouped by cluster. Distinct patterns emerge: Cluster 1 proteins peak in early adulthood, Cluster 2 increases steadily with age, while others decline or exhibit non-linear trajectories **(E)**. Cluster membership reflects coherent age-related proteomic signatures. The ten most representative proteins per cluster are listed, revealing distinct molecular identities (e.g., hemoglobin components, proteasomal subunits, cytoskeletal proteins, metabolic enzymes). Age trajectories of corresponding metabolomic features in the same REDS 13,091 index donors, mapped to the corresponding proteomic clusters **(F)**. Pathway enrichment for Cluster 1, revealing overrepresentation of innate immune and redox pathways, including complement/coagulation cascades, systemic lupus erythematosus–related pathways, neutrophil extracellular traps, autophagy regulation, and cholesterol metabolism **(G)**. Pathway enrichment for Cluster 2, dominated by proteostasis and metabolic redox pathways, including proteasome, pentose phosphate pathway, amino acid biosynthesis, glutathione homeostasis, and glycolysis/gluconeogenesis **(H)**.

### Impact of demographics on the RBC proteome

#### “Healthy donor” effect emerges from age-associated remodeling of the RBC proteome and metabolome

Age exerted a pervasive influence on RBC composition. A volcano plot of age–protein correlations identified extensive remodeling across cytoskeletal proteins (e.g., TUBA1B), chaperones (HSP90AA1), proteasomal subunits, redox-associated enzymes (PRDX1, GPX1), and multiple components of complement and lipid metabolism (**Figure 2B**). Of note, the presented analysis in this and subsequent sections is adjusted by plate, blood center and additive, though results substantially overlap with unadjusted analyses (offered for transparency in **Supplementary table 1**). Unsupervised fuzzy c-means analysis (c = 5) applied to the age-adjusted residual proteome resolved six age-associated proteomic clusters that captured coherent molecular trajectories across the lifespan. Visualization of the proteome using UMAP showed a smooth progression of these clusters across donor age, consistent with coordinated, rather than isolated, age-dependent changes (**Figure 2C**).

Heatmaps of the top 50 (top 10 by cluster) age-informative proteins revealed distinct cluster-specific temporal signatures (**Figure 2D**). Cluster 5 and 2 decreased or increased monotonically with advancing age, respectively. Other clusters displayed nonlinear or plateau-like trajectories, reflecting complex remodeling of metabolic, chaperone, and cytoskeletal pathways throughout adult life. Notably, proteins in Clusters 1 and 4 remained stable from early adulthood through age ∼65-70, but declined or increased rapidly after this age threshold, respectively – consistent with the concept of “marginal decade” posited by longevity experts. These cluster-level patterns were reproduced at the metabolomic level: age trajectories of metabolites corresponding to each proteomic cluster followed highly concordant patterns (**Figure 2F**; heat map breakdown in **Supplementary Figure 4A**), indicating tight proteome–metabolome coupling during RBC aging.

Pathway enrichment analyses demonstrated that age-dependent changes converge on immune and redox pathways. Cluster 1 (proteins declining with age) was enriched for complement/coagulation cascades, and cholesterol metabolism (**Figure 2G**). This observation is peculiar to the population of blood donors investigated here, in that a “healthy donor effect” introduces a selection bias, because an aging individual must meet certain health status criteria to still be able to donate blood.^39^ Consistent with this bias, Cluster 2 (proteins enriched with aging) was dominated by proteostasis and metabolic programs, including proteasome components, pentose phosphate pathway enzymes, amino-acid biosynthesis, glutathione homeostasis, and glycolytic/gluconeogenic pathways (**Figure 2H**) – all linked with improved RBC storage quality and post-transfusion circulatory performance^9,34,38,40,41^. Similarly, several immunoglobulin light chain components declined steadily with age, consistent with the negative selection of donors to minimize transfusion-related immunomodulation and alloimmunization events in recipients.^42^ In this view, the observation of ATIC (bifunctional enzyme involved in purine synthesis, also referred to as PAICS) was the top positive correlate to aging in healthy donors, while it has been reported that its inhibition via acetylation by ACSS2 is a hallmark of senescence-associated secretory phenotypes.^43^ Similar considerations apply to the pyrimidine synthesis-related enzyme PNPO, whose decline in the aging brain is linked to age-related cognitive decline.^44^

On the other hand, consistent with the broader aging literature, markers of extracellular matrix remodeling and fibrosis^45^, like fibronectin 1 (FN1) increased steadily in the aging donor. Fibrinogen chains (FGA, FGB, FGG) increased with aging, as extensively reported^46^, while albumin decline was observed – a predictor of mortality in the aging elderly^47,48^. HSP90AA1 – the top negative correlate to aging – is an established player in aging, as its decline is linked to age-related dysregulation of autophagy and neurodegeneration in the aging brain.^49^ Of note, apolipoproteins A1, A4, B, C3 increased with age, while others (e.g., L1) declined, consistent with the concept literature linking genetic polymorphisms of different apolipoproteins to the onset and severity of organ-specific age-related comorbidities and mortality.^50–53^

Together, these analyses show signatures of healthy aging in a selected blood donor population, along with established proteomics signatures of aging, which could be used to model trajectories of optimal aging for a general population.

### Sex-specific proteomic architecture and divergent aging trajectories pre-menopause

Along with age, sex emerged as a major axis of proteome-wide variation (**Figure 2A**). Differential abundance analysis revealed robust sex-associated signatures, with several proteins enriched in females (e.g., UBA1, SNCA, PKLR) and others in males (e.g., EIF1AX, FN1, VCP), spanning pathways related to redox regulation, proteostasis, lipid metabolism, and cytoskeletal organization (**Figure 3A**). These results also serve as additional internal validation of the quality of this dataset. For example, UBA1 is coded by a gene on chromosome X, it is expressed at higher levels in females, and males – carrying just one copy – are more susceptible to Vexas syndrome, a rare, late-onset autoinflammatory disease that causes systemic inflammation and blood disorders in adult males. It is caused by a somatic (non-inherited) mutation in the *UBA1* gene.^54^

**Figure 3.**
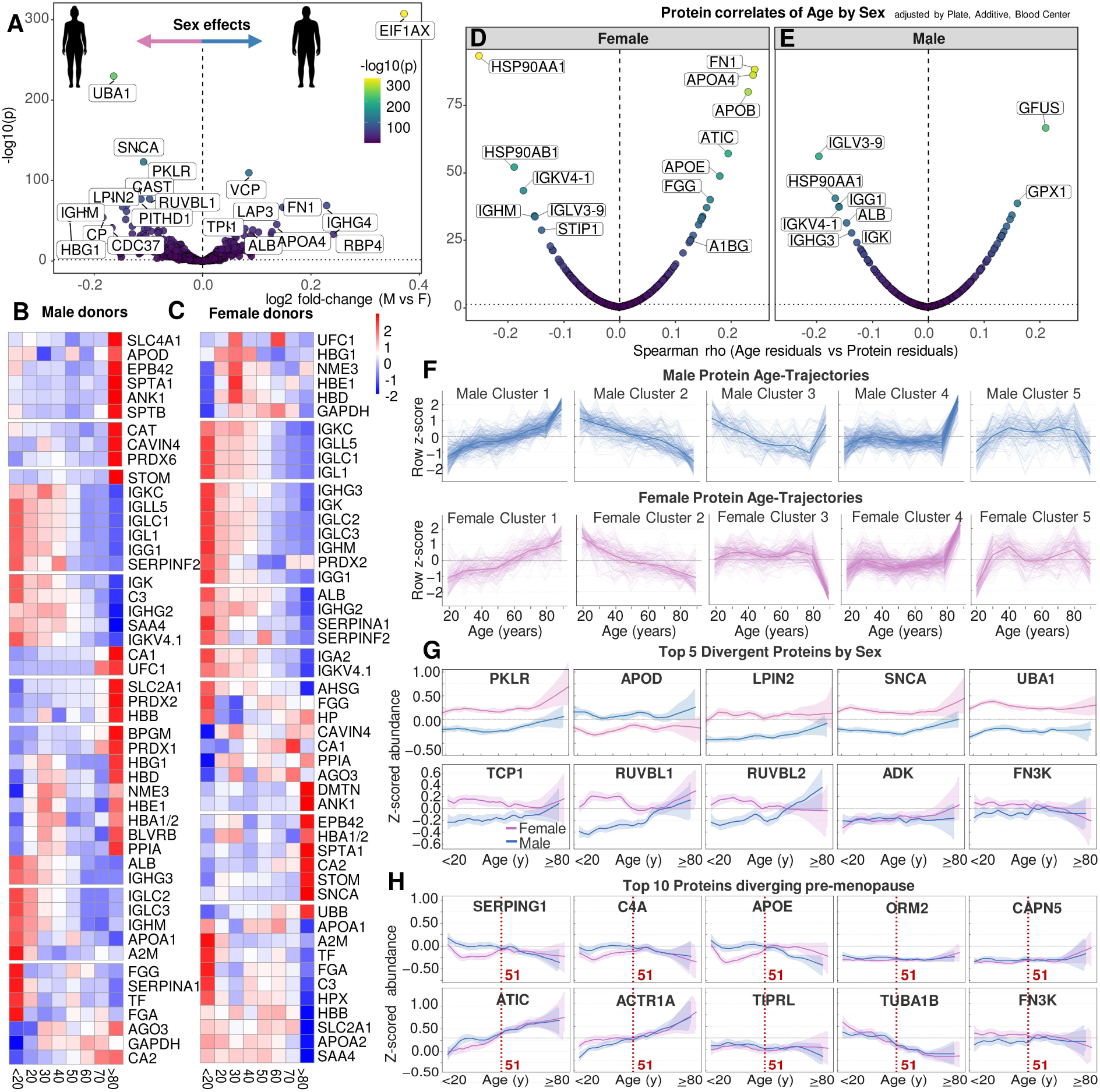
Sex-specific proteomic architecture and divergent age trajectories reveal coordinated remodeling of RBC biology in males and females. Volcano plot of sex differences in RBC protein abundance (Female/Male), identifying significantly enriched proteins in females (e.g., UBA1, SNCA, PKLR) and males (e.g., EIF1AX, VCP, FN1, LAP3). Effect sizes (log2 fold change) and –log10(p) reflect robust sex-dependent proteomic signatures **(A)**. Heatmaps of the most sex-divergent proteins across age decades, shown separately for males **(B)** and females **(C)**. Distinct age-by-sex architectures emerge, including enhanced immunoglobulin, cytoskeletal, and redox signatures in females and increased lipid- and hemoglobin-associated proteins in males. Volcano plots of age–protein correlations computed independently in female donors **(D)** and male donors **(E)**. While both sexes show strong age-associated remodeling, several proteins display highly sex-specific correlation patterns (e.g., ATIC, GFUS, PNPO, HSP90AA1 in females; GFUS, ATIC1, RUVBL1, P4HB in males). Sex-stratified age trajectories for proteomic clusters (fuzzy c-means = 5) derived separately in males (top) and females (bottom) **(F)**. Each cluster shows a unique temporal pattern - early-life peaks, midlife inflections, or late-life declines. Cluster 3 shows sex-specific coordination of metabolic, structural, and proteostasis pathways across the lifespan. The top five proteins most divergent by sex, either showing similar trends but different overall abundance by sex (PKLR, APOD, LPIN2, SNCA, UBA1) or different trajetories (TCP1, RUVBL1, RUVBL2, ADK, FN3K), plotted as smoothed age trajectories (mean ± 95% CI) **(G)**. Age trajectories for the top 10 proteins showing pre-menopausal divergence (marked by dashed red line at age 51, average age for menopause in the US). Several proteins - including SERPING1, C4A, APOE, ORM2, CAPN5, ATIC, ACTR1A, TRIP1, TUBA1B, FN3K - exhibit marked shifts around the menopausal transition, indicating hormone-linked or immune-metabolic remodeling of the female RBC proteome **(H)**.

Heatmaps stratified by age decade demonstrated that males and females do not simply differ in baseline protein abundance, but follow distinct temporal patterns across the lifespan (**Figure 3B–C**). In females, immune- and antioxidant-related proteins were consistently elevated, whereas males showed higher levels of hemoglobin-associated and lipid-linked proteins - consistent with established differences in basic hematological and complete blood count (CBC) parameters^55^. Sex-stratified correlations between age and protein abundance confirmed divergent age trajectories for proteins (**Figure 3D–E**) and metabolites (**Supplementary Figure 4B-C**). Several proteins displayed stronger age associations in one sex (e.g., ATIC, GFUS, PNPO, HSP90AA1 in females; GFUS, ATIC1, RUVBL1, P4HB in males), underscoring that aging of the RBC proteome is not uniform across sexes. Fuzzy c-means clustering performed separately in males and females (c = 5 per sex) revealed sex-specific cluster structures (**Figure 3F**). While almost all Clusters showed overlapping trajectories, Cluster 3 exhibited coordinated remodeling of proteostasis and metabolic pathways but with different timing and slope between males and females (top proteins in this cluster: TCP1, RUVBL1 and 2, ADK, FN3K). However, even for Clusters showing coherent trajectories between sexes, different basal levels were observed by sex throughout the lifespan - PKLR, APOD, LPIN2, SNCA, and UBA1 (**Figure 3G**). Finally, the menopausal transition imposed an additional inflection point in females. Several proteins - SERPING1, C4A, APOE, ORM2, CAPN5, ATIC, ACTR1A, TRIP1, TUBA1B, FN3K - showed clear breakpoints or shifts in slope around age 51, the average age of menopause in the U.S.^56^ (**Figure 3H**). These patterns highlight hormone-linked remodeling of immune, metabolic, and cytoskeletal networks in female RBCs.

### BMI and ethnicity influence inflammatory, metabolic, and ancestry-linked variation

BMI exerted a strong independent influence on RBC composition. The top 50 BMI-associated proteins included multiple complement and coagulation components (C3, C4A, CFB, FGA/FGB/FGG), acute-phase reactants (HP, HPX), lipid-associated proteins (APOB, APOA1), and glycolysis-related enzymes (**Figure 4A**). The top ten proteins jointly associated with BMI showed clear dose-dependent trends across WHO BMI categories - underweight to class III obesity - indicating a graded pro-inflammatory and lipid-linked signature (**Figure 4B**). A volcano plot of BMI-protein associations confirmed strong positive β-coefficients for complement and coagulation factors, reinforcing BMI as a major axis of systemic inflammation detectable in circulating RBCs (**Figure 4C**), as confirmed by pathway analysis of BMI-linked proteins (**Figure 4D**). This observation offers a molecular rationale for the observed hemolytic fragility of RBCs from donors with elevated BMI,^57,58^ consistent with a role for complement activation in post-transfusion hemolysis and hyperhemolysis reactions.^59^ Age trajectories of BMI-associated proteins revealed that inflammatory signatures were attenuated in aging donors who remained eligible to donate blood, consistent with a healthy-donor selection effect (**Figure 4E**).

**Figure 4.**
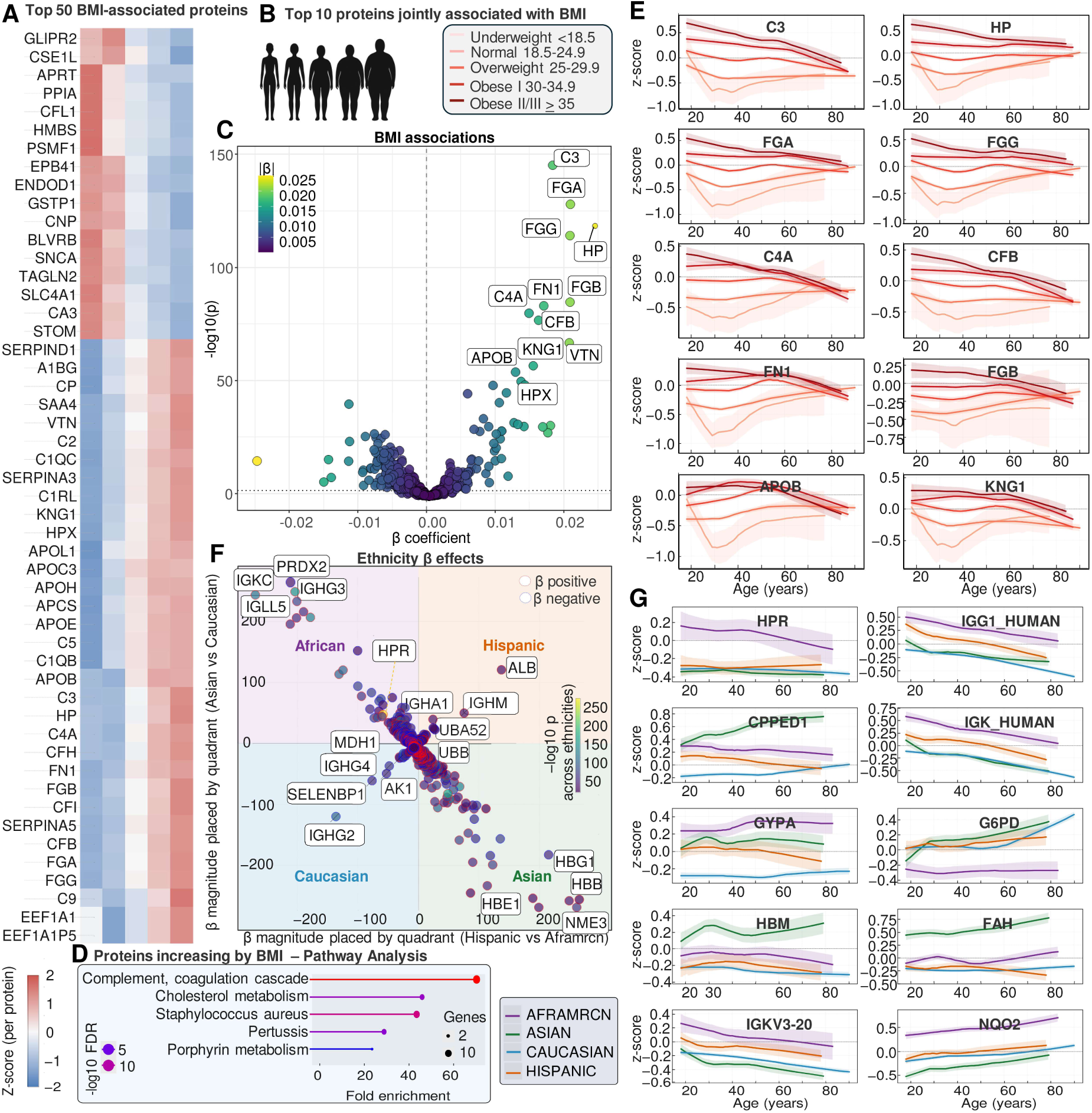
BMI and ethnicity shape distinct axes of RBC proteome variation with coordinated age-dependent remodeling. Heatmap of the top 50 BMI-associated proteins, ordered by effect size (adjusted by processing factors, age or sex) **(A)**. Higher BMI is associated with coordinated increases in complement components (e.g., C3, C4A, CFB, FGA/FGB/FGG), acute-phase reactants (HP, HPX), and lipid-associated proteins (APOB, APOA1), indicating inflammation- and lipid-linked remodeling of the RBC proteome. Top 10 proteins jointly associated with BMI shown across BMI categories (WHO classification), illustrating dose-dependent relationships across underweight to class III obesity **(B)**. Volcano plot of BMI–protein associations (β coefficients vs –log10(p)). Multiple complement and coagulation components (e.g., C3, HP, C4A, FGA, FGB, KNG1) and proteins involved in cholesterol metabolism show strong positive BMI associations, highlighting BMI as a major axis of systemic inflammation reflected in RBC composition **(C)**, as confirmed by pathway analysis **(D)**. Age trajectories of the most BMI-associated proteins (mean ± CI), demonstrating that BMI effects interact with aging, revealing a healthy donor effect with lower inflammation in aging subjects who still donate blood **(E)**. Multidimensional ethnicity effects on the RBC proteome visualized using a quadrant-based β-coefficient map **(F)**. Each protein is positioned in a 2-dimensional space according to the magnitude and direction of its ethnicity-associated regression coefficients: the x-axis encodes effects for Hispanic vs African American and the y-axis encodes effects for Asian vs Caucasian. This geometric placement allows simultaneous visualization of multi-ethnic differences, with background quadrants representing ancestry-associated proteomic signatures (either Caucasian, African, Hispanic or Asian vs the other three). Data color reflects the sign of the β-effect (positive or negative), and point size corresponds to the overall statistical significance across all ethnicity contrasts. Age trajectories of proteins most strongly diverging by ethnicity (e.g., HPR, IGG1, GYP.A, G6PD, FAH, NQO2) **(G)**.

Ethnicity introduced an additional structured axis of variability. A quadrant-based β-coefficient map positioned proteins according to effect sizes for Asian vs Caucasian and Hispanic vs African American comparisons (**Figure 4F; Supplementary Figure 5**). This two-dimensional representation revealed coherent ancestry-linked signatures, with proteins such as HPR, IGHG3, G6PD, FAH, NQO2, and multiple hemoglobin variants showing strong ethnicity-dependent abundance differences. This analysis recapitulates expected ancestry-associated effects, including reduced G6PD abundance in donors of African descent, consistent with the high global burden of X-linked G6PD deficiency and its ∼13% prevalence in this group within the REDS cohort. Age trajectories of the most ancestry-divergent proteins showed that ethnicity effects evolve with age, with some proteins (e.g., HPR, IGG1, GYP.A, G6PD) showing widening or narrowing differences across the lifespan (**Figure 4G**). These patterns reflect combined genetic and environmental influences on RBC composition across diverse U.S. donor populations.

Taken together, these analyses define the main demographic and processing axes along which the RBC proteome and metabolome vary, providing a framework to ask whether this information can be condensed into quantitative molecular “aging clocks” and to identify factors that accelerate or decelerate them.

### Aging clocks and their genetic, physiological, and clinical modifiers

We next trained penalized regression models (elastic net) to predict chronological age from proteomic or metabolomic features, thereby deriving multi-omics RBC aging clocks (**Figure 5A–E**, **Supplementary Figure 6**). To maximize generalizability, we trained the clocks in a meta-data–agnostic and batch-robust framework. Age, sex, BMI and other demographic fields or hematological parameters were excluded from the predictor space, and any features that were simply age recodings or showed near-perfect correlation with age (e.g., decade-based age bins) were removed to prevent information leakage. Proteomic and metabolomic features were then median-imputed and normalized (either as within-sample ranks or as feature-wise z-scores), so that each feature was placed on a comparable scale and residual plate/center effects were minimized. The same normalization parameters learned in the 13,091-donor training set were then reused unchanged when mapping external or future cohorts onto the trained clock. This design makes the models explicitly independent of cohort-specific metadata and batch structure, allowing ΔAge estimates in new datasets to be interpreted as deviations from a fixed, population-level molecular aging axis rather than as artifacts of local preprocessing or demographic composition.

**Figure 5.**
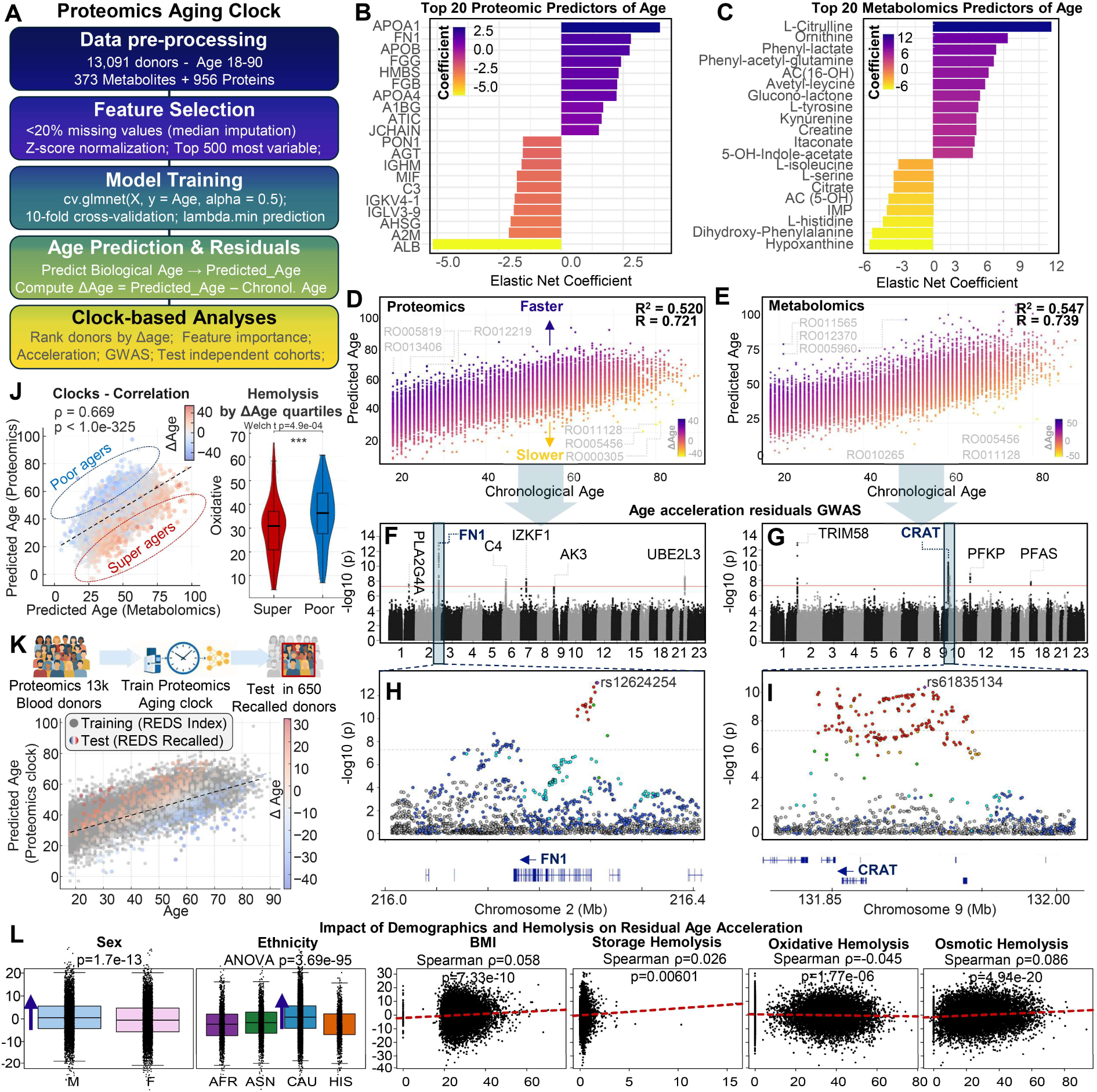
Multi-omic aging clocks identify proteomic and metabolic predictors of biological age, reveal genomic architecture of age acceleration, and link RBC aging to hemolysis phenotypes. Workflow for constructing proteomics- and metabolomics-based aging clocks across 13,091 REDS donors **(A)**. After preprocessing (missing value filtering, imputation, Z-scoring), the top 500 variable features were used for elastic net regression with cross-validation to predict biological age and compute age acceleration (ΔAge = Predicted – Chronological Age). Top 20 proteomic **(B)** and metabolomic **(C)** predictors of age ranked by elastic net coefficients. Performance of the proteomics **(D)** and metabolomics **(E)** aging clocks, showing predicted vs chronological age with high correlation (Proteomics rho = 0.721; Metabolomics rho = 0.739, cross-validated). Colors indicate magnitude and direction of residual age acceleration (faster vs slower biological aging). Manhattan plots from GWAS of age acceleration residuals, revealing significant loci for proteomic ΔAge at FN1 (chr2) and C4/IKZF1 (chr7), and for metabolomic ΔAge at CRAT (chr9) and TRIM58 (chr1) **(F–G)**. Locus zoom for selected signals: FN1 locus (chr2, top SNP rs12624254); CRAT locus (chr9, top SNP rs61835134) **(H–I)**. Cross-clock correlation between proteomic and metabolomic ΔAge shows consistency across donors (ρ = 0.669, p < 10^-325^) **(J)**. Quartet analysis of extreme donors identifies “super agers” (ΔAge<15) and “poor agers” (ΔAge>15), the latter showing increased susceptibility to hemolysis after oxidant insult (p = 4.9×10⁻4). Training-testing scheme for external validation: the proteomics clock trained in **13k donors** accurately predicts age in the 650-donor recalled cohort (R = 0.66), demonstrating model generalizability **(K)**. Impact of demographics and hemolysis phenotypes on age acceleration residuals **(L)**. Sex and ethnicity show significant differences in ΔAge distributions (Sex p = 1.7×10^-13^; Ethnicity ANOVA p = 3.6×10^-95^). BMI, storage hemolysis and osmotic hemolysis – but not oxidative hemolysis – accelerate aging residuals, linking physiological and RBC-intrinsic stress pathways to accelerated biological aging **(L)**.

In the 13,091 index donors, both aging clocks performed strongly. The proteomics clock achieved a tight relationship between predicted and chronological age (R = 0.721), and the metabolomics clock showed similarly robust performance (R = 0.739), with mean absolute errors of approximately 10.4 and 8.8 years, respectively (**Figure 5D–E**). Both models exhibited the characteristic pattern of molecular clocks in large cohorts, with systematic underestimation of predicted age in the oldest donors, likely reflecting a “healthy donor” selection effect rather than limited sample size, as the dataset includes over 1,600 donors aged ≥65. Feature importance analyses highlighted declines in albumin, increases in complement and coagulation factors, apolipoproteins, redox enzymes, FN1 and key amino-acid–derived metabolites (citrulline, ornithine, kynurenine – consistent with prior work,^60,61^ hydroxy-acyl-carnitines – markers of impaired fatty acid oxidation and exercise intolerance^62^; and phenyl-lactate – a natural organic acid derived from phenylalanine metabolism primary produced by *Lactobacilli*) as major contributors to the aging signal (**Figure 5B–C**). A merged proteomic–metabolomic model further strengthened this relationship, yielding improved overall prediction accuracy (r = 0.801 – **Supplementary Figure 6G-H**).

We next investigated the genetic architecture of ΔAge and residual age acceleration (ΔAge/Age). Residual age acceleration (ΔAge/Age) provides a normalized measure of biological aging rate, independent of chronological age. Raw ΔAge values (Predicted – Observed age) tend to correlate with age because all aging clocks over- or underestimate certain age ranges. By regressing predicted age on chronological age and taking the residuals, the analysis removes this built-in statistical coupling. The resulting residuals capture only the age-independent deviations - that is, how much “faster” or “slower” an individual’s RBC molecular profile appears to be aging relative to peers of the same chronological age. This enables unbiased comparisons across age groups, detection of modifiers of biological aging (e.g., BMI, hemolytic stress, genotype), and identification of “super agers” and “poor agers” whose biological trajectories diverge most strongly from expected age norms. GWAS of proteomic ΔAge and residual age acceleration identified loci at FN1, C4/IKZF1, AK3, UBE2L3, and PLA2G4A, implicating extracellular matrix remodeling, immune regulation, mitochondrial nucleotide metabolism, ubiquitination, and phospholipid signaling in RBC molecular aging (**Figure 5F, Supplementary Figure 7A,C–D**). GWAS of metabolomic ΔAge highlighted loci at CRAT, TRIM58, PFKP, and PFAS, pointing to roles for fatty-acid oxidation, erythropoiesis-linked transcriptional control, glycolysis, and one-carbon metabolism in shaping metabolic aging (**Figure 5G, Supplementary Figure 7B,E–F**).

### Protein levels and aging clocks are reproducible across multiple donations from the same donors, as a function of genetic traits

To evaluate whether the demographic and biological signatures identified in the 13,091-donor index cohort generalized to independent samples and to quantify how storage modifies the RBC proteome, we next leveraged the recalled-donor proteomics dataset comprising 1,929 samples from 650 donors profiled at days 10, 23, and 42 of storage (**Supplementary Figure 8A**). Storage-day trajectories revealed coordinated remodeling across cytoskeletal, proteasomal, redox, and translational pathways (**Supplementary Figure 8B**). Among all quantified proteins, HBG2 (the fetal γ-globin) emerged as the top storage-responsive feature, showing a consistent decline across storage while also exhibiting strong inter-individual variability. Demographic analyses further showed that HBG2 abundance was markedly higher in younger donors, in females, and in individuals with lower BMI (**Supplementary Figure 8C–D**), closely mirroring demographic effects observed in the index cohort and providing an orthogonal internal validation of the proteomics. GWAS of HBG2 levels identified a single, genome-wide–significant **cis-QTL** at the β-globin gene cluster on chromosome 11 (lead SNP rs3759074), consistent with classical haplotypes regulating fetal hemoglobin persistence, and a secondary trans association at BCL11A (**Supplementary Figure 8E**), the canonical regulator of the fetal-to-adult hemoglobin switch^63^. Together, these findings illustrate how the recalled cohort captures both genetically encoded and demographically patterned variation in globin expression, while also resolving superimposed storage-dependent dynamics, while suggesting that donors carrying specific BCL11A SNPs or from certain demographics resulting in higher fetal hemoglobin expression may represent eligible candidates for selected donation to pediatrics recipients.

The recalled cohort additionally enabled a stringent test of temporal reproducibility, as 650 donors had mass-spectrometry proteomes generated twice - once from their index unit at Day 42 and again from their recalled unit 2–12 months later (**Supplementary Figure 9A**). Across all proteins, the distribution of cross-donation correlations was strongly right-skewed (median ρ ≈ 0.55), with a subset of proteins exhibiting exceptionally high reproducibility (ρ ≈ 0.60–0.75, p < 10⁻⁵⁰), including NQO2, IGHG4, C3, GSTT1, FAH, GPX1, HPX, KNG1, and fibrinogen chains (**Supplementary Figure 9B–C**). For the top reproducible protein, NQO2, proteomic QTL analysis identified a single, sharp cis-QTL on chromosome 6 directly overlying the NQO2 locus (**Supplementary Figure 9D–E**), indicating that intra-donor stability in protein abundance is overwhelmingly genetically driven and not an artifact of pre-analytical variability. Pathway enrichment of the 50 most reproducible proteins highlighted complement/coagulation, glutathione metabolism, cholesterol and vitamin digestion pathways, glycolysis/gluconeogenesis, and xenobiotic metabolism - core RBC processes known to be under strong genetic and developmental constraint (**Supplementary Figure 9F**). Remarkably, ΔAge estimates were also highly reproducible over time, with ΔAge estimates from clock training on the smaller recalled donor cohort tightly tracking ΔAge from the index unit (ρ = 0.66, p = 6.5×10^-54^; **Supplementary Figure 9G**), demonstrating that RBC molecular aging rate is a stable, donor-specific trait preserved across independent donations separated up to a year. Collectively, these analyses show that demographic, genetic, and biological signatures identified in the index cohort are reaffirmed in the recalled cohort, while the longitudinal sampling additionally reveals storage-dependent proteome remodeling and robust intra-donor stability across >6 months (on average) as regulated by genetic traits. The reproducibility of the RBC proteome and external validation with cis-pQTL hits from GWAS approaches both also constitute hallmarks of a high-quality, deeply reproducible proteomics dataset.

Beyond training an independent clock on the recalled donor sample dataset, we also tested the performance of the clock trained on the index cohort in this new dataset. Proteomic and metabolomic ΔAge were strongly correlated across donors (ρ ≈ 0.67, p<1×10^-325^), indicating that the clocks capture convergent aspects of RBC molecular aging (**Figure 5J**), consistent with the influence of donor-related genetic traits in the reproducibility of clock performances. Stratification by extreme ΔAge values identified “super agers” and “poor agers” (ΔAge = predicted – chronological age by <-15 or >15 years, respectively); donors in the latter group showed significantly higher susceptibility to oxidant-induced hemolysis, directly linking accelerated RBC molecular aging to increased hemolytic vulnerability (**Figure 5K**). Indeed, modest albeit significant residual age-acceleration was observed as a function of sex (faster in males), ethnicity (faster in donors of Caucasian descent), BMI (higher in obese donors), storage an osmotic hemolysis (accelerators – **Figure 5L**).

These multi-omic ΔAge residuals captured inter-individual variability in RBC molecular aging, enabling downstream analyses of biological modifiers, hemolytic phenotypes, and genetic determinants of accelerated or decelerated aging trajectories. Given the observed association between hemolytic traits and aging clock acceleration, we asked how inherited hemolytic conditions modulate ΔAge. In REDS RBC Omics donors, increasing severity of G6PD deficiency – the most common enzymopathy in humans that is associated to increased hemolytic propensity following oxidant insults (e.g., during storage in the blood bank) but it is not ground for deferral – or sickle cell trait (one allele of *HBB* E6V - HbAS) were both linked to accelerated aging clocks (ΔAge >10-**Supplementary Figure 10A-B**).

To test these results prospectively, aging clock testing against proteomics and metabolomics results from RBC omics analyses in an independent cohort of 248 healthy subjects (HbAA), subjects carrying sickle cell trait (HbAS; n=174) or sickle cell disease (HbSS) were associated with stepwise increases in age acceleration (**Supplementary Figure 10C-H**). These observations support the concept that chronic redox and hemolytic stress accelerate RBC molecular aging, linking classical hemolytic genotypes to broader systemic aging signatures captured by the clocks.

### High-frequency donation slows molecular aging unless iron deficiency develops, while iron repletion resets the clock

We next leveraged longitudinal iron studies to examine how donation intensity and iron status impact molecular aging. In the REDS cohort, high-frequency donors (>3 donations over the past 12 months) displayed significantly lower ΔAge than the training cohort. However, this held true only for repeat donors without iron deficiency, as defined by ferritin levels above 15 ng/mL (**Supplementary Figure 11A–C**). These results suggest that, in the context of preserved iron stores, frequent blood donation may be associated with slower biological aging of RBCs. Of note, it has been argued that the act of donating blood may favor slower aging by lowering iron stores ∼200 mg per donated unit^64^. However, while whole blood can be donated every 56 days (up to six times per year in the United States), frequent donation is a well-recognized risk factor for iron deficiency—even in donors who remain eligible on the basis of normal hemoglobin levels (≥13.0 g/dL in men and ≥12.5 g/dL in women).

Therefore, to test this hypothesis, we generated proteomics and metabolomics data from RBCs obtained from volunteers enrolled in the Donor Iron-Deficiency Study (DIDS)^65^. In this double-blind randomized study, adult whole-blood donors with iron deficient erythropoiesis (ferritin < 15 µg/L and zinc protoporphyrin > 60 µmol/mol heme), who were still eligible to donate, were randomized to receive either 1 g low-molecular-weight intravenous iron dextran or placebo, and after a subsequent donation their RBC 24-h post-transfusion recovery (as well as donor cognition and quality-of-life) was assessed — with the outcome showing no significant improvement in post-transfusion RBC circulatory capacity. Systemic iron recovery was accompanied by trends toward increased whole-brain iron and myelination (P=0.04 and P=0.02 respectively), and enhanced functional-MRI network activation (P<0.001), although overall cognitive scores did not differ.^66^ Donors participating in the DIDS trial had accelerated ΔAge at baseline relative to the general REDS iron-replete controls. Randomization to iron dextran supplementation versus saline revealed that only iron repletion, not placebo, normalized ΔAge, returning aging clock residuals to levels indistinguishable from the control training set (**Supplementary Figure 11D–G**). Consistent with previously reported findings, iron supplementation improved hematologic and iron indices, including hemoglobin, ferritin, iron, and zinc protoporphyrin, an observation here expanded with the characterization of broad remodeling of the RBC proteome and metabolome, including normalization of TF levels, elevation of iron-dependent aminoacyl-tRNA synthases (CARS1, YARS, etc), metabolic enzymes (GAPDH), structural membrane proteins, and proteostasis pathways (**Supplementary Figure 11F-G**). Importantly, these systemic changes to RBC were significantly associated to changes in brain myelin and iron levels and cognitive function. MRI-based measurements of brain iron and myelin in iron-utilizing regions such as globus pallidus, putamen, and caudate increased after iron repletion, and omics–brain correlation analyses showed that many of the strongest proteomic and metabolomic predictors of ΔAge - such as APOA1, citrulline, and neurotransmitter-related metabolites including dopamine derivatives - also tracked with brain iron content and cognitive performance (**Supplementary Figure 11H–J**). These observations suggest that the RBC aging clock does not only capture intrinsic RBC biology, but reflects a broader systemic axis linking iron metabolism, RBC composition, brain iron status, and neurocognitive function.

### Shiny app resource for testing of any new proteomics or metabolomics dataset vs REDS

To facilitate reuse of these models, we provide an interactive Shiny application that allows the interested readers to train and apply RBC aging clocks on their own omics datasets using the REDS cohort as a reference resource (https://github.com/Angelo-DAlessandro/AgingClocks). The app ingests large proteomics or metabolomics CSV files, harmonizes sample IDs and metadata, and offers multiple preprocessing options (raw values, within-sample ranks, or autoscaling) while explicitly excluding Age, Sex, and BMI - and any age-like correlates (e.g., age-decade bins) - from the predictor space to prevent information leakage. Users can either retrain an elastic-net clock on the REDS training file or map new cohorts onto an existing clock, visualizing predicted versus chronological age, ΔAge distributions, and residual age acceleration relative to the donor reference. The interface reports feature importance, generates volcano plots contrasting “younger” versus “older” residuals, and tests modifiers such as sex, BMI, and hemolysis metrics. All plots and underlying statistics can be exported as SVG/CSV, enabling transparent benchmarking of external cohorts against a healthy, population-scale RBC aging reference.

### Proteomic architecture of hemolysis and transfusion response

To understand how proteomic variation influences functional storage phenotypes, we mapped associations between proteins and storage, osmotic, and oxidative hemolysis, as well as hemoglobin increments at 24 hours after single-unit transfusion (ΔHgb) (**Figure 6**). A hive plot summarizing these associations highlighted densely connected modules involving inflammation, complement and coagulation, membrane scaffolding, and nucleotide metabolism (**Figure 6A**). Chromosomal Manhattan-style plots (proteome wide association study – PWAS) showed that oxidative hemolysis, in particular, was associated with strong proteomic signals from antioxidant enzymes and cytoskeletal proteins, including PCMT1, GPX1, PRDX1, SOD1 or STOM, ANK3, and SLC4A1 (**Figure 6B–D**), the latter class showing notable overlap with genome-wide determined signatures of osmotic fragility^34^. Elastic net models trained to predict hemolysis modalities or transfusion increments converged on overlapping sets of predictors, including apolipoproteins, complement and coagulation factors, membrane proteins such as ankyrin and vitronectin, and metabolic enzymes such as G6PD, PKLR, MDH1, PAICS, and regulatory subunits of phosphatases (**Figure 6E–F**, **Supplementary Figure 12**). A neural network model using the full proteome to predict ΔHgb further emphasized hierarchical integration of complement (C1QB, C3, C4B, CFH), purine metabolism (PNPO, PAICS), cytoskeletal (ANK1, STOM), apolipoproteins (APOA1), coagulation (FGA) and redox features (PCMT1, G6PD, SOD1, GPX1 - **Figure 6G**). When comparing the top 30 determinants across all phenotypes, recurring patterns were observed for proteins predicting 2 or more phenotypes, above all APOA1, ANK1, C3, STOM and G6PD (**Figure 6E-G**). Age trajectories of key predictors, stratified by quartiles of hemolytic propensity or transfusion response, showed that donors with higher hemolysis or lower ΔHgb exhibit steeper age-linked increases in proteasome and redox markers (**Figure 6H**, **Supplementary Figure 13**), again supporting a coupling between donor aging and RBC aging.

**Figure 6.**
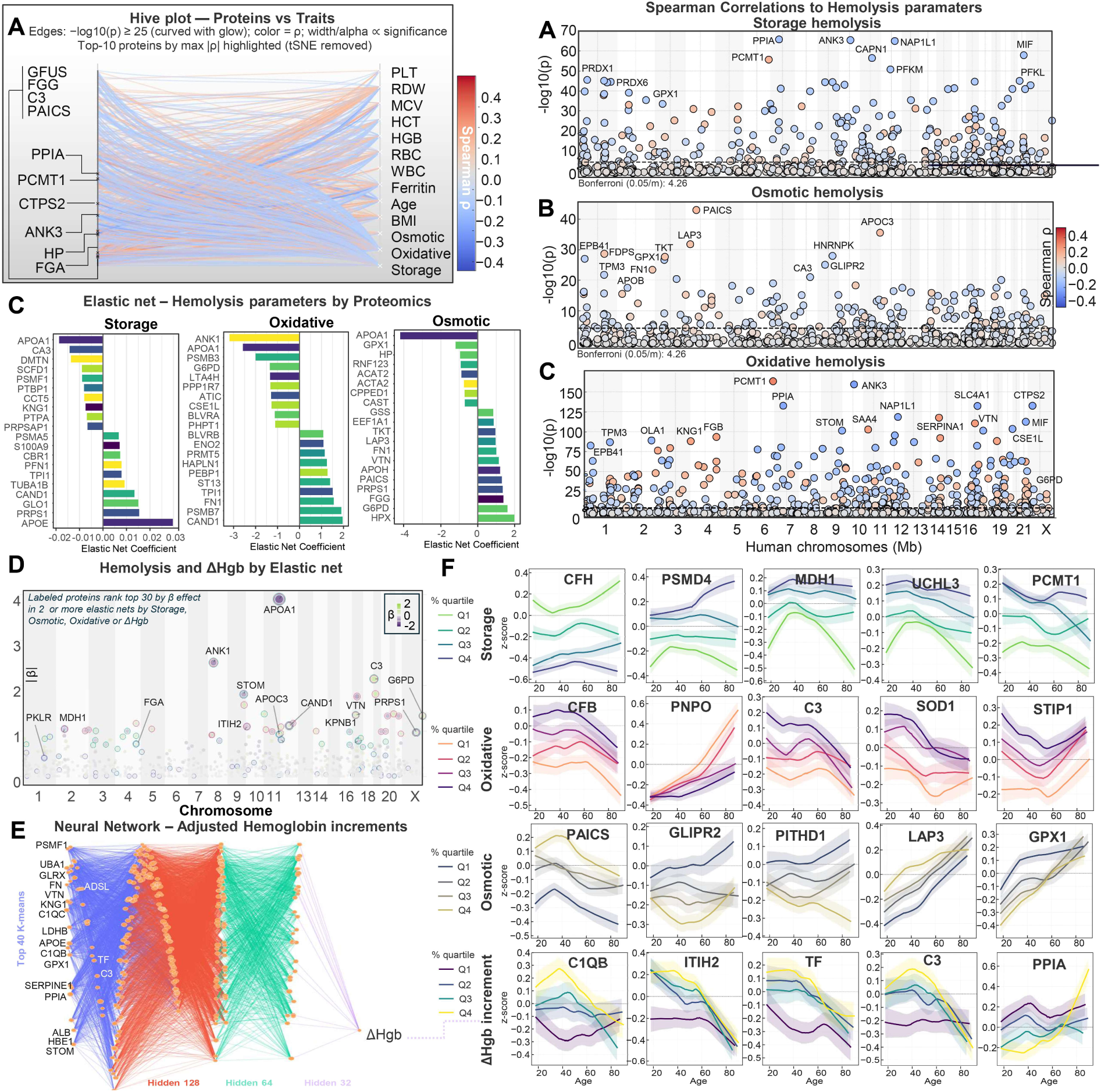
Proteomics-informed Machine Learning models of RBC storage, osmotic, and oxidative hemolysis and hemoglobin increments after single unit transfusion. Hive plot summarizing proteome-trait associations for all hemolysis parameters (storage, osmotic, oxidative) and related donor variables (complete blood count – CBC indices, BMI, Age) **(A)**. Edges represent significant associations at –log10(p) ≥ 2.5, colored by direction (blue = negative; red = positive). The top-10 proteins most strongly associated with hemolysis are labeled, highlighting inflammation-, complement-, and membrane-linked proteins including HP, FGA, FGG, PAICS, GFUS, and CTPS2. Manhattan-style scatterplots of Spearman correlations for storage hemolysis **(B)**, osmotic hemolysis **(C)**, and oxidative hemolysis **(D)**. Each protein is positioned by chromosomal location mapping for the respective coding gene (x-axis) and –log10(p) of the correlation (y-axis), with point color reflecting Spearman rho correlation modules and direction. Oxidative hemolysis shows the strongest signal magnitude, identifying antioxidant, redox, and cytoskeletal proteins (GPX1, PRDX1, STOM, ANK3, SLC4A1) as top contributors. Beyond Spearman correlations, we trained elastic net models identifying proteins predicting storage, osmotic, and oxidative hemolysis **(E)** or adjusted hemoglobin increments in transfusion recipients at 24h after single unit blood transfusion events **(F)**. Shared drivers across multiple functional outcomes include apolipoproteins (APOA1, APOA3) complement/coagulation proteins (FGA, C3), membrane proteins (ANK1, STOM, VTN), and metabolic regulators (G6PD, PKLR, MDH1, PAICS, PPP1R), suggesting overlapping mechanisms of RBC resilience and hemolytic propensity in the bag or after transfusion. Neural network architecture modeling adjusted hemoglobin increment (ΔHgb) using the full RBC proteome **(G)**. The network illustrates hierarchical feature integration across hidden layers (128 → 64 → 32 nodes). Strong positive and negative contributors cluster within complement, coagulation, cytoskeletal, and redox modules, highlighting how RBC intrinsic protein composition predicts post-transfusion hemoglobin response. Age trajectories (mean ± CI across ΔHgb or hemolysis quartiles) of proteins most strongly associated with each hemolysis phenotype **(H)**.

### Peptide-level resolution reveals PTM-driven heterogeneity, genetic modifiers, and a peptide-based aging axis

To dissect biochemical sources of variability that remain invisible at the protein level, we next analyzed the RBC proteome at **peptide-level** resolution, quantifying >30,000 peptides across all donors. PHATE embedding of peptide versus parent-protein abundances revealed extensive intra-protein heterogeneity (**Supplementary Figure 14A**), with structurally diverse proteins—including NME3, MIF, PRDX2, UFC1, CAVIN4, and hemoglobin β (HBB; **Supplementary Figure 14B–D**)—forming broad peptide “clouds” whose spatial dispersion reflected PTMs, cleavage events, and sequence-specific detectability.

HBB provided an archetypal case: peptide-level correlations formed discrete PTM-defined clusters corresponding to oxidation, methylation, glutathionylation, acetylation, and ubiquitin-remnant (K–GlyGly) modifications (**Supplementary Figure 14D–E**), revealing a structural and regulatory landscape far richer than suggested by aggregate protein abundance alone. Ranking peptides by their deviation from parent-protein abundance identified the top 5% most discordant peptides (**Supplementary Figure 14C**), which fell into three major biochemical categories:

i. peptides enriched for PTM hotspots—including glutathionylation, oxidation, acetylation, and cysteine-directed alkylation (**Supplementary Figure 14E–F**);
ii. peptides prone to enzyme- or ROS-mediated proteolysis (e.g., band 3 N-terminus; **Supplementary Figure 14H**)^67,68^;
iii. peptides harboring amino-acid substitutions arising from common missense polymorphisms. This analysis also revealed genetic modifiers of peptide abundance. Carriers of the G6PD N126D missense variant - prevalent in donors of African descent - showed selective enrichment of oxidative HBB peptides, consistent with elevated redox burden in G6PD-deficient RBCs (**Supplementary Figure 14I**). Conversely, donors carrying the PCMT1 (also known as PIMT) V120I allele (rs4816; >60% allele frequency in Asian ancestry groups) exhibited markedly reduced correlations between total PIMT abundance and the V120-containing peptide, suggesting genotype-dependent destabilization or altered PTM state (**Supplementary Figure 14J–K**).

Collectively, these analyses demonstrate that peptide-level mapping resolves previously unmeasurable axes of RBC heterogeneity, exposing PTM-, fragmentation-, and genotype-dependent features that are completely occluded at the protein level. Importantly, this degree of resolution - including direct quantification of PTM isoforms and missense-variant peptides - is inherently inaccessible to affinity-based high-throughput platforms such as Olink or SomaScan, whose antibody- and aptamer-based measurements interrogate predefined epitopes rather than individual peptide species.

We next systematically quantified how demographic variables shape peptide-level biology. Volcano plots demonstrated widespread peptide-specific associations with age, BMI, sex, and ethnicity (**Supplementary Figure 15A–D**), with PTM-bearing peptides disproportionately represented among the strongest effects. These patterns highlight that demographic remodeling of the RBC proteome often operates at the level of specific PTM isoforms or cleavage products, rather than shifting bulk protein abundance.

We then trained a peptide-based molecular aging clock leveraging the full peptide matrix. Despite excluding demographic predictors, the peptide clock accurately recapitulated chronological age (ρ = 0.75; MAE = 8.90 years), with strong out-of-fold calibration (**Supplementary Figure 16B–D**). Feature attribution revealed key PTM- and cleavage-driven predictors, including oxidized or acetylated HBB peptides, immunoglobulin fragments, and albumin/APOA1-derived peptides (**Supplementary Figure 16E**). Age-trajectory plots of top contributors highlighted diverse peptide classes with monotonic, nonlinear, or saturating age behavior, demonstrating that PTM-level remodeling encodes a rich biological aging signal independent of protein-level trends (**Supplementary Figure 16F**). Together, these analyses reveal that RBC peptide biology captures a multilayered structure of PTM heterogeneity, proteolysis, genetic variation, and demographic influences, and provides an interpretable, high-resolution axis for modeling biochemical aging.

### Biological age predicts long-term donor retention

Finally, we asked whether baseline omics signatures from the index donation collected between December 2013 through October 2015 predict long-term donor behavior a decade later in 2025. Focusing on two of the four REDS centers for which follow-up data were available (BSRI and ITxM, both now under the Vitalant umbrella), we classified donors as active if they had at least one donation in 2025 and inactive otherwise (**Figure 7A**). Age-adjusted logistic models incorporating demographics, CBC indices, and hemolysis phenotypes identified expected influences, including higher likelihood of long-term activity among donors with greater pre-index donation frequency, older age, certain ethnicities, and specific blood groups, especially O- or + (**Figure 7B**), consistent with targeted donor retention campaign for universal donors. When proteomic and metabolomic data were added, elastic net, logistic regression, and random forest classifiers all achieved robust discrimination between active and inactive donors, with AUC values around 0.70 (**Figure 7C–F**). The molecular features driving these models overlapped extensively with aging-clock predictors and included FN1, APOA1, ITIH2, HRG, KNG1, C1QB, citrulline, glutathione disulfide or cysteine adducts, phenylacetylglutamine, kynurenine, nicotinamide metabolites, and glycolytic intermediates, suggesting that donors who remain engaged in blood donation are biologically “younger” and metabolically healthier at the level of their RBC proteome and metabolome.

**Figure 7.**
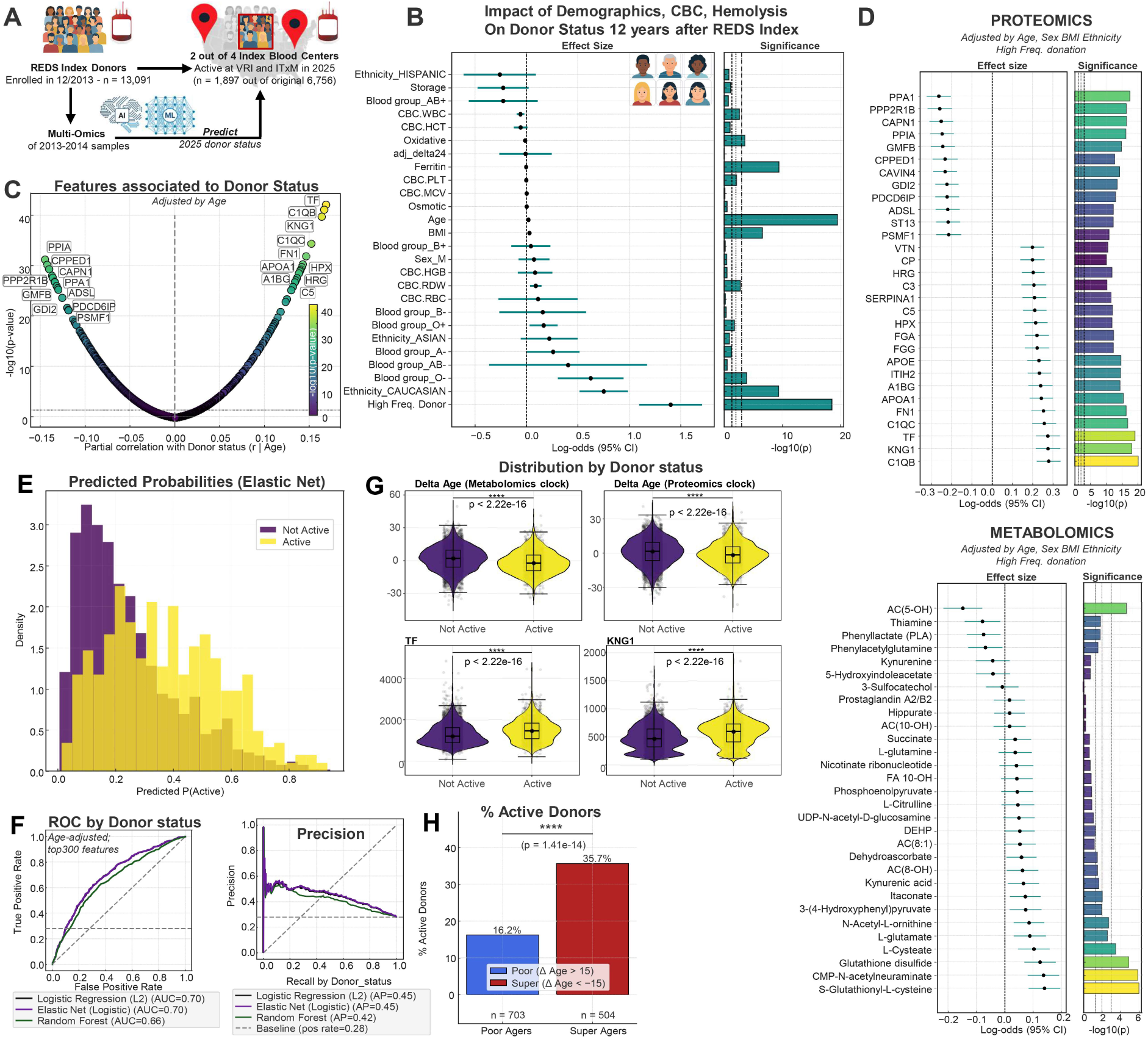
Multi-omic signatures predict long-term donor activity and link biological aging, hemolysis, and molecular features to sustained donation behavior 12 years after REDS Index enrollment. Study schema **(A)**. Among 13,091 index donors, multi-omics data were used to train models that classify donors as Active (≥1 donation in 2025) or Not Active, across two of the four U.S. blood centers participating in the original REDS RBC Omics study. Influence of demographics, CBC parameters, and hemolysis phenotypes on donor status (age-adjusted logistic regression) **(B)**. Higher likelihood of long-term donor activity is associated with original donation frequency (high for donors with >3 donations 12 months prior to REDS Index), donor age (older donors are more likely to return), ethnicity (higher likelihood to be active for donors of Caucasian or Asian descent) and specific blood groups (e.g., O negative or positive, mirroring establishment efforts to secure targeted recruitment/retention of universal blood donors). Volcano plot **(C)** or elastic net-based model **(D)** of omics features associated with donor status, after adjusting for donor age. Effect sizes for proteomic (top) and metabolomic (bottom) predictors of long-term donor activity (adjusted for age, sex, BMI, ethnicity, and donation frequency). Despite adjusting for donor age, key proteomic and metabolomics positive predictors include several high ranked variables in both aging clocks: FN1, APOA1, ITIH2, HRG, KNG1, and C1QB for proteomics; citrulline, glutathione (disulfide or cysteine adduct), phenylacetylglutamine, kynurenine, nicotinamide metabolites, and glycolytic intermediates, reflecting metabolic health and RBC energy/redox status. Distribution of predicted probabilities from elastic net classification, showing clear separation between Active vs Not Active donors **(E)**. Prediction performance evaluated by ROC curves and precision–recall analysis. Logistic regression, elastic net, and random forest models all achieve robust discrimination (AUC ≈ 0.70), demonstrating strong multi-omic predictability of long-term donor activity **(F)**. Biological age acceleration is strongly associated with donor status. Active donors exhibit significantly lower ΔAge (younger biological age) in both metabolomics and proteomics clocks (p < 2.2×10^-16^). Active donors also show higher levels of proteins and metabolites associated with healthier RBC phenotypes, including TF, KNG1, and others **(G)**. The proportion of Active donors is markedly different among Super Agers (ΔAge ≤ –15) and Poor Agers (ΔAge ≥ +15). Super Agers show 35.7% long-term activity versus 16.2% among Poor Agers (p = 1.4×10⁻¹⁴), linking biological aging trajectories to sustained donor engagement **(H)**.

Consistent with this interpretation, donors who were still active in 2025 had significantly lower ΔAge than those who were inactive, in both proteomics and metabolomics clocks (**Figure 7G**). Stratification by extreme ΔAge delineated “super agers” (ΔAge ≤ –15) and “poor agers” (ΔAge ≥ +15), revealing that super agers were more than twice as likely to still be donating 12 years after index donation (35.7% vs 16.2%, p = 1.4×10⁻¹⁴) (**Figure 7H**). These data establish a direct link between molecular aging trajectories captured in stored RBC units, hemolytic resilience, systemic metabolic health, and long-term donor retention, positioning the RBC proteome and metabolome as practical, scalable readouts of both transfusion quality and donor “healthspan.”

## DISCUSSION

This study provides the first population-scale map of the human red blood cell proteome, revealing that demographic and genetic factors reproducibly shape RBC molecular composition across independent donations. By integrating proteomics, metabolomics, genomics, storage phenotypes, transfusion outcomes, and long-term donor follow-up, we show that inter-individual variation in RBC biology is structured, heritable, and clinically meaningful. For example, a key translational insight arises from the discovery that HBG2 abundance in stored RBC units varies by donor genetics, demographics, and storage duration. Because fetal hemoglobin increases oxygen affinity and can blunt the Bohr effect, units with higher HBG2 levels may be better suited for pediatric, neonatal, or fetal transfusion, where elevated HbF is physiologically appropriate. Conversely, units with low HBG2 may be preferable for adult recipients requiring maximal oxygen delivery. This establishes the foundation for rational precision transfusion medicine strategies aimed at donor-recipient matching based on proteo-genomic phenotypes rather than indirect clinical surrogates.

Based on proteomics and metabolomics signatures, we trained machine learning models (e.g., elastic-net regression) to inform molecular aging clocks that further stratify donors in ways directly relevant to transfusion outcomes. Accelerated ΔAge identifies donors whose units are intrinsically more susceptible to spontaneous, osmotic, and oxidative hemolysis; who generate lower hemoglobin increments after transfusion; and who carry signatures of chronic oxidative burden, low iron availability, or genetic hemolytic predisposition (e.g., G6PD deficiency, sickle cell trait or SCD). Conversely, donors with “younger” molecular phenotypes provide units with superior storability and post-transfusion performance, as determined by clinical records of hemoglobin increments in transfusion recipients. These observations support pre-donation phenotyping - analogous to blood typing or infectious disease screening - to prioritize lower negative ΔAge donors for vulnerable patient groups such as trauma victims, neonates, or chronically transfused individuals. Given the intrinsic stability of the genome, and the reproducibility of the proteome across donations we report in this study, this phenotyping could occur at first donation, while repeated metabolic characterization over time could account for changes in dietary, environmental, professional or recreational exposures (i.e., the exposome).^69^

The responsiveness of ΔAge to interventions highlights its clinical tractability. Iron-deficient donors exhibited marked age acceleration that was reversed by iron repletion in an independent intervention cohort. ΔAge could therefore serve as a biomarker for iron supplementation programs aimed at improving donor health and product quality. Similarly, the slowed molecular aging in high-frequency iron-replete donors suggests that that donor aging clocks can be reset through simple interventions like iron supplementation. Linkage of ΔAge and iron repletion to brain iron/myelin and cognitive function – a hallmark age-related comorbidity – suggests that the aging clocks developed here may open a peripheral (blood) window on the aging of the central nervous system.

Importantly, because repeat blood donors represent a highly screened, relatively healthy subset of the population, the clocks we describe reflect ideal, “healthy aging” trajectories. As such, they provide a robust reference framework for evaluating less healthy or clinically enriched cohorts, enabling prospective comparisons of molecular aging across chronic disease, metabolic syndromes, hemoglobinopathies, or inflammatory states. This positions the RBC aging clock as a broadly deployable metric for studying systemic aging biology in diverse clinical contexts. This task will be facilitated by the Shiny app we offer to the interested reader for testing on any new dataset against our resource omics database from the REDS study.

Our proteogenomic analyses reveal additional pathways with immediate translational relevance. Polymorphisms at regions coding for FN1 link aging clocks to extracellular matrix remodeling and fibrosis secondary to senescence-associated secretory phenotypes.^70^ Variation at CRAT and PFAS - enzymes linked to acyl-carnitine flux in mitochondria and purine synthesis (with PAICS and PNPO ranking amongst the top sex-dependent age correlates), whose dysregulation represent a hallmark of aging^71^ - tracks with hemolysis susceptibility and transfusion efficacy, suggesting metabolic nodes whose modulation may improve RBC resilience ex vivo. Likewise, membrane, complement, and redox signatures associated with hemolysis risk (PCMT1, GPX1) provide candidate markers for real-time quality assessment of stored units.

Finally, the observation that ΔAge predicts donor activity 12 years later links RBC molecular phenotypes to long-term donor health behavior. Individuals with biologically younger RBCs remain active donors, suggesting that RBC molecular aging is not only a marker of cell health but also of broader donor healthspan. Identification of donors with long-term retention potential may inform current recruitment and retention campaigns to counteract the global trends towards shortages in blood inventories owing to aging of the donor population.^72^

Together, these results shift the role of RBC omics from descriptive biology to actionable clinical stratification, laying the foundation for precision transfusion strategies, targeted donor interventions, and the use of RBC molecular aging as a scalable biomarker of human health.

## STAR METHODS

### Recipient Epidemiology and Donor evaluation Study (REDS) RBC Omics

A detailed description of the study design, enrollment criteria and main outcomes has been provided in previous publications,^32,33^ and – while redundant with the literature - will be included below with significant overlap with published methods.

### Donor recruitment in the REDS RBC Omics study

#### Index donors

A total of 13,758 donors were enrolled in the Recipient Epidemiology and Donor evaluation Study (REDS) RBC Omics at four different blood centers across the United States (https://biolincc.nhlbi.nih.gov/studies/reds_iii/). Of these, 97% (13,403) provided informed consent and 13,091 were available for metabolomics analyses in this study – referred to as “index donors”. A subset of these donors were evaluable for hemolysis parameters, including spontaneous (n=12,753) and stress (oxidative and osmotic) hemolysis analysis (n=10,476 and 12,799, respectively) in ∼42-day stored leukocyte-filtered packed RBCs derived from whole blood donations from this cohort ^73^. Methods for the determination of FDA-standard spontaneous (storage) hemolysis test, osmotic hemolysis (pink test) and oxidative hemolysis upon challenge with AAPH have been extensively described elsewhere ^58^. The Vitalant Blood Establishment Computer System (BECS) was queried to determine which index donors from two of the four REDS RBC Omics centers (the former BSRI and ITxM centers) made a donation in 2025.

#### Recalled donors

A total of 643 donors scoring in the 5^th^ and 95^th^ percentile for hemolysis parameters or who donated nine or more donations over the prior 24 months at the index phase of the study were invited to donate a second unit of pRBCs, a cohort henceforth referred to as “recalled donors”.^74^ These units were assayed at storage days 10, 23 and 42 for hemolytic parameters and mass spectrometry-based high-throughput metabolomics ^75^, proteomics ^76^, lipidomics ^77^ and ICP-MS analyses ^78^. Under the aegis of the REDS-IV-P project ^32^, a total of 1,929 samples (n=643, storage day 10, 23 and 42) were processed with this multi-omics workflow. Of the 643 parent units, 638 were genotyped for 362 G6PD SNPs.

#### Sample Preparation

Frozen 96-well plates containing RBCs samples were thawed on ice. Each 10 µL RBCs sample was lysed with 90 µL of distilled water. Then, 5 µL of the lysed sample was transferred to a deep-well 96-well plate containing 45 µL of 4% SDS and 20 mM N-ethylmaleimide (NEM) in 50 mM triethylammonium bicarbonate (TEAB), along with 0.25 µg of internal standard (yeast Alcohol dehydrogenase, Worthington Biochemical Corporation). Samples were incubated at room temperature for 30 minutes with shaking to ensure complete alkylation.

Following incubation, phosphoric acid was added to a final concentration of 1.2% (v/v), and six volumes of binding buffer (90% methanol, 100 mM TEAB, pH 7.0) were added. The mixture was gently mixed before transfer to an S-Trap™ 96-well Mini Plate (Protifi). Approximately 200 µL of each sample was loaded per well and centrifuged at 1500 × g for 2 minutes. The plate was washed four times with 300 µL of binding buffer under the same centrifugation conditions.

For on-column digestion, the S-Trap™ plate was moved to a new 1 mL 96-well collection plate, and 1 µg of sequencing-grade trypsin (Promega) in 125 µL of 50 mM TEAB was added to each well. Samples were incubated at 37 °C for 6 hours in a humidified incubator.

Peptides were eluted sequentially as follows:

a. 100 µL of 50 mM TEAB, centrifuged at 1500 × g for 2 minutes;
b. 100 µL of 0.2% formic acid in water, centrifuged at 1500 × g for 2 minutes;
c. 100 µL of 80% acetonitrile in 0.1% formic acid, centrifuged at 1500 × g for 2 minutes.

All eluates were combined, dried using a SpeedVac concentrator, and resuspended in 200 µL of 0.1% formic acid. Samples were filtered using an ISOLUTE® FILTER+ Plate (25 µm/0.2 µm) to remove particulates. A 10 µL aliquot of each filtered sample was diluted with 190 µL of 0.1% formic acid and stored in a 96-well plate.

#### Nano-UHPLC-MS/MS Analysis

For LC-MS analysis, 20 µL of each diluted peptide sample was loaded onto conditioned and equilibrated Evotips (Evosep, Denmark) according to the manufacturer’s instructions. Tips were washed twice with 200 µL of solvent A (0.1% formic acid in water) and stored in solvent A until analysis.

Peptide separation was achieved using the Evosep One LC system (EV-1000, Evosep, Denmark) with the pre-programmed 200 samples per day (SPD) method. Samples were separated over 5.6 minutes on a 4 cm × 150 µm reversed-phase column packed with 1.9 µm C18 beads. The mobile phases consisted of solvent A (0.1% formic acid in water) and solvent B (0.1% formic acid in acetonitrile).

Mass spectrometric analysis was performed on a timsTOF Pro system (Bruker Daltonics) operated in diaPASEF mode. Ion mobility was set from 0.70 to 1.30 1/k₀, with an accumulation time of 100 ms and a 100% duty cycle. Collision energy was applied as a linear function of 1/k₀ (0.6 1/k₀ = 20 eV; 1.60 1/k₀ = 59 eV). The diaPASEF windows were optimized using pydiAID based on precursor density and chromatographic separation for the 200 SPD method. The total cycle time was 1.35 s.

#### Spectral Library Generation

A project-specific spectral library was generated by pooling small aliquots from each sample. The pooled digest was fractionated by high-pH reversed-phase chromatography on a Gemini-NH C18 analytical column (4.6 × 250 mm, 3 µm particles) at a flow rate of 0.6 mL/min. The mobile phases were 10 mM ammonium bicarbonate (pH 10) as phase A and 10 mM ammonium bicarbonate with 75% acetonitrile (pH 10) as phase B. Peptides were separated using a linear gradient from 0–5% B in 10 minutes, 5–50% B in 80 minutes, 50–100% B in 10 minutes, followed by a 10 minute hold at 100% B.

A total of 96 fractions were collected and concatenated into 24 pooled fractions by combining every 24th fraction (e.g., 1, 25, 49, 73, etc.). The pooled fractions were dried and reconstituted in 100 ul of 0.1 % FA.

#### Quality Control and Maintenance

Each 96-well plate included wash, blank and technical replicate samples to monitor reproducibility and carryover. Instrument calibration, buffer refilling, and ion transfer interface (ITI) cleaning were performed weekly. The column and emitter were replaced approximately every 24–26 plates to maintain chromatographic performance.

Overall, the system performance remained stable throughout the REDS project, with consistent peptide identification and retention time reproducibility.

#### Statistical modeling of age effects on protein abundance, adjusted by Plate, Additive, and Blood Center

To quantify age-associated changes in protein abundance while adjusting for technical covariates, we used a partial Spearman correlation approach. For each protein *j*, we first regressed chronological age on Blood Center, Additive, and Plate, and extracted the residuals (age residuals):

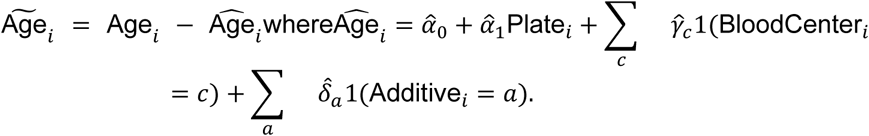

In parallel, we regressed the normalized abundance of protein *j* on the same covariates and extracted protein residuals:

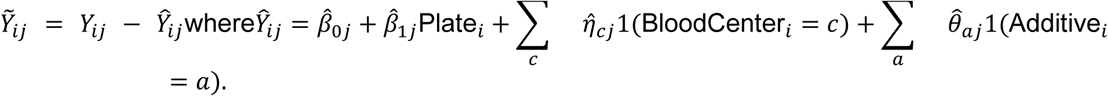

We then computed Spearman’s rank correlation coefficient between the two sets of residuals, yielding a **partial Spearman correlation** for protein *j*:

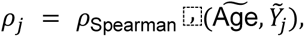

which captures the association between age and protein *j* after adjusting for Blood Center, Additive solution, and Plate effects.

P values were obtained using cor.test() in R and corrected for multiple hypothesis testing using the Benjamini–Hochberg procedure. The resulting *ρ_j_* values and adjusted P values were used to construct volcano plots summarizing the age-associated RBC proteome signature.

### Age-binned clustering of protein abundance trajectories

To visualize age-related protein abundance patterns, adjusted by plate, additive or blood center, we first binned donors by chronological age in 5-year intervals and summarized protein intensities within each bin. Adjusted protein-level intensity data were extracted from the full proteomics matrix, after separating metadata from quantitative columns. Age was discretized into 5-year bins using cut() in R, with inclusive lower bounds and non-overlapping intervals. For each age bin, we computed the median intensity of every protein across donors in that bin, yielding a matrix of median protein abundance (proteins × age bins). We then selected the 50 most variable proteins based on the variance of their median profiles across age bins and standardized each protein’s trajectory by z-score normalization across bins. To identify groups of proteins with similar age-related trajectories, we applied fuzzy c-means clustering (cmeans function, e1071 package) to the scaled matrix with *k* = 8clusters and a maximum of 100 iterations. Proteins were ordered by their primary c-means cluster assignment, and the resulting clustered matrix was visualized as a heatmap using pheatmap with a blue–white–red color scale, fixed row/column ordering, and row gaps marking cluster boundaries. The final heatmap was exported as an SVG file using svglite for downstream figure editing.

### uMAP embedding and k-means clustering of donor proteomes

uMAP was used to visualize inter-donor structure in the REDS population-scale proteomics dataset. Features with >20% missing values were removed, and remaining missing intensities were imputed using within-feature medians. Protein features were then z-score standardized as

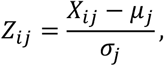

where *μ_j_* and *σ_j_* denote the sample mean and standard deviation for protein *j*. To determine the intrinsic cluster structure of donor proteomic profiles, we applied k-means clustering over a grid of candidate cluster numbers and evaluated cluster quality using the silhouette index (factoextra::fviz_nbclust), selecting *k* = 5as the optimal solution. The final clustering was obtained by minimizing within-cluster sum of squares in the k-means objective

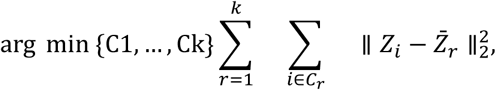

where *Z_i_* is the standardized proteome of donor *i* and *Z̄_r_* is the centroid of cluster *r*.

uMAP was then run on the standardized protein matrix (umap::umap) with three embedding dimensions (*n*_components_ = 3) to generate a nonlinear manifold representation of the donor proteome landscape. The resulting embedding (UMAP1, UMAP2, UMAP3)was merged with k-means cluster labels and visualized using Plotly as a fully interactive 3D scatterplot. Cluster-discriminating proteins were identified by performing a Kruskal–Wallis test for each protein *j*:

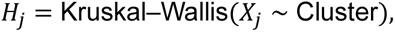

and corresponding p values were false-discovery–rate–adjusted using the Benjamini–Hochberg procedure. The top discriminating proteins (lowest adjusted *P*) were exported as a .csv file for downstream interpretation.

#### Shiny application for RBC Aging Clock Modeling and external cohort mapping

We developed a Shiny application that enables users to train red blood cell (RBC) molecular aging clocks, apply them to external cohorts, and visualize ΔAge-associated proteomic and metabolomic signatures using a standardized analytical workflow. Users upload CSV files containing quantitative features and metadata; variables encoding chronological Age, Sex, BMI, age bins, or age-derived fields are automatically excluded from the predictor space to prevent information leakage. The app supports several preprocessing strategies, including raw values 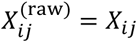, within-sample rank normalization 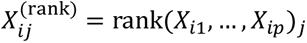, and feature-wise autoscaling 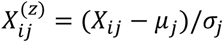, where *μ_j_* and *σ_j_* are estimated from the REDS training set and reapplied unchanged when mapping external datasets; missing values are imputed using sample-wise medians. RBC aging clocks are trained using elastic-net regression implemented in glmnet, solving 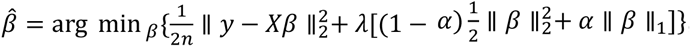, with hyperparameters *α* and *λ* selected by cross-validation. Biological age predictions for sample *i* are computed as 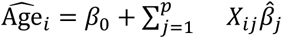 and biological age acceleration is defined as 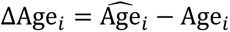_i_. The app visualizes predicted versus chronological age, ΔAge distributions, and residual variation, and enables differential analyses of molecular features associated with ΔAge: for each feature *j*, the app computes Spearman correlation *ρ_j_* = *ρ*_Spearman_(*X_⋅j_*, ΔAge), P values via cor.test(), and Benjamini–Hochberg–adjusted FDRs, generating volcano plots (*ρ_j_* vs. −log_10_ *P_j_*) and exportable result tables. External datasets are automatically harmonized with the REDS reference clock, normalized using stored (*μ_j_*, *σ_j_*)parameters, and mapped to predicted age and ΔAge values, enabling benchmarking of new cohorts against a population-scale “healthy ager” reference. All visualizations, including predicted versus observed age, ΔAge histograms, feature-importance coefficients, and volcano plots, are generated using ggplot2 and exported as fully editable SVG files, with all intermediate tables available as CSV for downstream analysis.

#### Peptide-level PHATE embedding and peptide–protein coupling

To visualize structure in the peptide-level landscape and its relationship to parent proteins, we performed PHATE embedding on peptide intensities from stored RBCs. After excluding meta-data, we identified protein and peptide quantitative columns and aligned samples by a shared ID field. Intensities were log2-transformed as log _2_(*x* + 1)for both peptides and proteins. For each peptide, we inferred its parent protein by splitting the peptide column name at the first underscore and computed a Spearman correlation between that peptide and its parent protein across donors, requiring at least 200 samples with finite values; peptides with insufficient data were assigned *ρ* = NA. PHATE was then applied to the peptide matrix using the phateR package, treating each peptide as an observation (row) and donors as features (columns): we ran phate(t(pep_mat), knn = 10, ndim = 2, seed = 42) to obtain a 2D embedding of peptides. The resulting coordinates (PHATE1, PHATE2) were merged with peptide–protein correlation coefficients, and peptides were visualized as points colored by their Spearman ρ with the corresponding parent protein, using a viridis color scale. The PHATE map was exported as an SVG file for high-resolution figure preparation. In an additional exploratory analysis, we approximated co-embedding of proteins by performing PCA on the same peptide space (on *t*(pep_mat)) and projecting protein intensities (from *t*(prot_mat)) into the first two principal components, allowing parent proteins to be overlaid as labeled points on the peptide manifold background.

#### GWAS for aging clock and pQTL studies

The workflow for the GWAS analysis of aging clocks and pQTL analyses of selected protein levels in REDS Index (HBG2, NQO2) is consistent with previously described methods from our pilot mQTL study on 250 recalled donors^79^ and follow up studies on specific metabolic pathways (e.g., glycolysis^80^) other proteins (e.g., GPX4^81^) on the full index cohort. Details of the genotyping and imputation of the RBC Omics study participants have been previously described by Page, et al.^34^ Briefly, genotyping was performed using a Transfusion Medicine microarray^82^ consisting of 879,000 single nucleotide polymorphisms (SNPs); the data are available in dbGAP accession number phs001955.v1.p1. Imputation was performed using 811,782 SNPs that passed quality control. After phasing using Shape-IT^83^, imputation was performed using Impute2 ^84^ with the 1000 Genomes Project phase 3^84^ all-ancestry reference haplotypes. We used the R package SNPRelate^85^ to calculate principal components (PCs) of ancestry. We performed association analyses using an additive SNP model in the R package ProbABEL^86^ and 13064 REDS Index donor study participants who had HBG2, NQO2 protein levels or ΔAge and ΔAge residuals calculated based on proteomics or metabolomics aging clock data, and imputation data on serial samples from stored RBC components that passed respective quality control procedures. We adjusted for sex, age (continuous), frequency of blood donation in the last two years (continuous), blood donor center, and ten ancestry PCs. Statistical significance was determined using a p-value threshold of 5×10^-8^. We only considered variants with a minimum minor allele frequency of 1% and a minimum imputation quality score of 0.80. The OASIS: Omics Analysis, Search & Information a TOPMED funded resources ^87^, was used to annotate the top SNPs. OASIS annotation includes information on position, chromosome, allele frequencies, closest gene, type of variant, position relative to closest gene model, if predicted to functionally consequential, tissues specific gene expression, and other information.

#### Determination of hemoglobin and bilirubin increment via the vein-to-vein database

Association of ΔAge or protein levels with hemoglobin increments was performed by interrogating the vein-to-vein database, as described in Roubinian et al.^40^. Analyses were performed on the whole REDS population and stratified for measurements ΔHgb increment at 24h (adjusted by relevant donor and recipient variables).

#### Vein-to-vein database: General Study Design

We conducted a retrospective cohort study using electronic health records from the National Heart Lung and Blood Institute (NHLBI) Recipient Epidemiology and Donor Evaluation Study-III (REDS-III) program available as public use data through BioLINCC ^88,89^. The database includes blood donor, component manufacturing, and patient data collected at 12 academic and community hospitals from four geographically diverse regions in the United States (Connecticut, Pennsylvania, Wisconsin, and California) for the 4-year period from January 1, 2013 to December 31, 2016. Genotype and metabolomic data from the subset of blood donors who participated in the REDS-III RBC-Omics study ^90^ was linked to the dataset using unique donor identifiers.

#### Study Population and Definitions

Available donor proteomics data were linked to issued RBC units using random donor identification numbers. Among transfusion recipients, we included all adult patients who received a single RBC unit during one or more transfusion episodes between January 1, 2013 and December 30, 2016. Recipient details included pRBC storage age, and blood product issue date and time. We collected hemoglobin levels measured by the corresponding clinical laboratory prior to and following each RBC transfusion event (0h and 24h after transfusion).

#### Transfusion Exposures and Outcome Measures

All single RBC unit transfusion episodes linked to proteomics data were included in this analysis. A RBC unit transfusion episode was defined as any single RBC transfusion from a single donor with both informative pre- and post-transfusion laboratory measures and without any other RBC units transfused in the following 24-hour time period. The outcome measures of interest were change in hemoglobin (ΔHb; g/dL) following a single RBC unit transfusion episode. These outcomes were defined as the difference between the post-transfusion and pre-transfusion levels. Pre-transfusion thresholds for these measures were chosen to limit patient confounding (e.g., underlying hepatic disease). For pre-transfusion hemoglobin, the value used was the most proximal hemoglobin measurements prior to RBC transfusion, but at most 24 hours prior to transfusion. Furthermore, we excluded transfusion episodes where the pre-transfusion hemoglobin was greater than 9.5 g/dL, and the hemoglobin increment may be confounded by hemorrhage events. For post-transfusion hemoglobin, the laboratory measure nearest to 24-hours post-transfusion, but between 12- and 36-hours following transfusion was used.

#### Metabolomics and lipidomics analyses in REDS RBC Omics (Index and Recalled)

Omics methods described below follow exactly those described in previous publications, as referenced in each section that follows.

#### High-throughput metabolomics

Metabolomics extraction and analyses in 96 well-plate format were performed as described, with identical protocols for human or murine RBCs^91,92^. RBC samples were transferred on ice on 96 well plate and frozen at −80 °C at Vitalant San Francisco (human RBCs) or University of Virginia (murine RBCs) prior to shipment in dry ice to the University of Colorado Anschutz Medical Campus. Plates were thawed on ice then a 10 uL aliquot was transferred with a multi-channel pipettor to 96-well extraction plates. A volume of 90 uL of ice cold 5:3:2 MeOH:MeCN:water (*v/v/v*) was added to each well, with an electronically-assisted cycle of sample mixing repeated three times. Extracts were transferred to 0.2 µm filter plates (Biotage) and insoluble material was removed under positive pressure using nitrogen applied via a 96-well plate manifold. Filtered extracts were transferred to an ultra-high-pressure liquid chromatography (UHPLC-MS — Vanquish) equipped with a plate charger. A blank containing a mix of standards detailed before ^93^ and a quality control sample (the same across all plates) were injected 2 or 5 times each per plate, respectively, and used to monitor instrument performance throughout the analysis. Metabolites were resolved on a Phenomenex Kinetex C18 column (2.1 x 30 mm, 1.7 um) at 45 °C using a 1-minute ballistic gradient method in positive and negative ion modes (separate runs) over the scan range 65-975 m/z exactly as previously described.^91^ The UHPLC was coupled online to a Q Exactive mass spectrometer (Thermo Fisher). The Q Exactive MS was operated in negative ion mode, scanning in Full MS mode (2 μscans) from 90 to 900 m/z at 70,000 resolution, with 4 kV spray voltage, 45 sheath gas, 15 auxiliary gas. Following data acquisition, .raw files were converted to .mzXML using RawConverter then metabolites assigned and peaks integrated using ElMaven (Elucidata) in conjunction with an in-house standard library^94^.

#### High-throughput Oxylipin Analysis

Analyses were performed as previously published via a modified gradient optimized for the high-throughput analysis of oxylipins.^10,14,16^ Briefly, the analytical platform employs a Vanquish UHPLC system (Thermo Fisher Scientific, San Jose, CA, USA) coupled online to a Q Exactive mass spectrometer (Thermo Fisher Scientific, San Jose, CA, USA). Lipid extracts were resolved over an ACQUITY UPLC BEH C18 column (2.1 x 100 mm, 1.7 µm particle size (Waters, MA, USA) using mobile phase (A) of 20:80:0.02 ACN:H_2_O:FA and a mobile phase (B) 20:80:0.02 ACN:IPA:FA. For negative mode analysis the chromatographic the gradient was as follows: 0.35 mL/min flowrate for the entire run, 0% B at 0 min, 0% B at 0.5 min, 25%B at 1 min, 40%B at 2.5min, 55% B at 2.6min, 70% B at 4.5 min, 100% B at 4.6 min, 100% B at 6 min, 0% B at 6.1 min and 0% B at 7 min. The Q Exactive mass spectrometer (Thermo Fisher) was operated in negative ion mode, scanning in Full MS mode (2 μscans) from 150 to 1500 m/z at 70,000 resolution, with 4 kV spray voltage, 45 sheath gas, 15 auxiliary gas. Calibration was performed prior to analysis using the Pierce^TM^ Positive and Negative Ion Calibration Solutions (Thermo Fisher Scientific).

#### High-throughput Lipidomic Analysis

Lipid extracts were analyzed (10 µL per injection) on a Thermo Vanquish UHPLC/Q Exactive MS system using a previously described^77^ 5 min lipidomics gradient and a Kinetex C18 column (30 x 2.1 mm, 1.7 µm, Phenomenex) held at 50 °C. Mobile phase (A): 25:75 MeCN:H_2_O with 5 mM ammonium acetate; Mobile phase (B): 90:10 IPA:MeCN with 5 mM ammonium acetate. The gradient and flow rate were as follows: 0.3 mL/min of 10% B at 0 min, 0.3 mL/min of 95% B at 3 min, 0.3 mL/min of 95% B at 4.2 min, 0.45 mL/min 10% B at 4.3 min, 0.4 mL/min of 10% B at 4.9 min, and 0.3 mL/min of 10% B at 5 min. Samples were run in positive and negative ion modes (both ESI, separate runs) at 125 to 1500 m/z and 70,000 resolution, 4 kV spray voltage, 45 sheath gas, 25 auxiliary gas. The MS was run in data-dependent acquisition mode (ddMS^2^) with top10 fragmentation. Raw MS data files were searched using LipidSearch v 5.0 (Thermo).

#### Statistical analysis

Acquired data was converted from raw to mzXML file format using Mass Matrix (Cleveland, OH, USA). Analysis was done using MAVEN, an open-source software program for oxylipin analysis. Lipid assignments and peak integration were performed using LipidSearch v 5.0 (Thermo Fisher Scientific). Samples were analyzed in randomized order with a technical mixture injected interspersed throughout the run to qualify instrument performance. Data analysis and Statistical analyses – including hierarchical clustering analysis (HCA), linear discriminant analysis (LDA), uniform Manifold Approximation and Projection (uMAP), correlation analyses and Lasso regression were performed using both MetaboAnalyst 5.0 and RStudio (2024.12.1 Build 563).

## Supporting information

Supplementary Table - Raw data and Elaborations

## Author Contribution

Method development: MD. Proteomics: MD, AS, KCH, AD. Proteomics data searches and normalization: AVI, SB, AS, KCH, AD. Metabolomics and lipidomics analyses: JAR, DS, TN, AD. Biostatistics and Bioinformatics: AVI, SB, GRK, FF, ALM, XD, GPP, NR, AD. REDS RBC Omics: MS, XD, SK, SLS, PJN, MPB, NHR. Systems biology models: ZBH; Vein-to-vein database: NHR; QTL analyses: GRK, ALM, GPP. Figure preparation: AD. Conceptualization: AD. Writing and finalization: first draft by AD and all co-authors reviewed and approved the final version.

## Funding

This study was supported by funds by the National Heart, Lung, and Blood Institute (NHLBI) (R01HL148151, R01HL146442, R01HL149714 to AD; R01HL126130 to NHR). The REDS RBC Omics and REDS-IV-P CTLS programs are sponsored by the NHLBI contract 75N2019D00033, and from the NHLBI Recipient Epidemiology and Donor Evaluation Study-III (REDS-III) RBC Omics project, which was supported by NHLBI contracts HHSN2682011-00001I, −00002I, −00003I, −00004I, −00005I, −00006I, −00007I, −00008I, and −00009I. G.R.K was supported by grants from the National Institute of General Medical Sciences (NIGMS), F32GM124599. The content is solely the responsibility of the authors and does not necessarily represent the official views of the National Institutes of Health. The authors would like to thank all the donor volunteers who participated in this study and all the global blood donor communities for their life-saving altruistic gifts.

## Clinical trial

The NHLBI Recipient Epidemiology Donor Evaluation Study (REDS)-III Red Blood Cell Omics (RBC-Omics) and Vein to Vein databases are accessible at https://biolincc.nhlbi.nih.gov/studies/reds_iii/ and Genomics data are deposited at dbGaP Study Accession: phs001955.v1.p1

## Competing Interest

The authors declare that AD, KCH, TN are founders of Omix Technologies Inc. AD, TN are Scientific Advisory Board (SAB) members for Hemanext Inc. AD is SAB member for Macopharma Inc and SynthMed Biotechnologies. All the other authors have no conflicts to disclose in relation to this study. AD, TN, MD, DS, KCH filed a provisional patent application on the proteomics and multi-omics aging clock.

## Data and Materials availability

All **METHODS** are extensively described in **Supplementary Materials.pdf**. All raw data and elaborations are included in Supplementary Table 1.xlsx. The Aging Clocks Shiny App is available for download at https://github.com/Angelo-DAlessandro/AgingClocks. Further information and requests for resources and reagents should be directed to and will be fulfilled by the Lead Contact, Angelo D’Alessandro.

## Notes

https://github.com/Angelo-DAlessandro/AgingClocks

## REFERENCES

1. Sender, R., Fuchs, S., and Milo, R. (2016). Revised Estimates for the Number of Human and Bacteria Cells in the Body. PLoS Biol 14, e1002533. 10.1371/journal.pbio.1002533.

2. Bianconi, E., Piovesan, A., Facchin, F., Beraudi, A., Casadei, R., Frabetti, F., Vitale, L., Pelleri, M.C., Tassani, S., Piva, F., et al. (2013). An estimation of the number of cells in the human body. Ann Hum Biol 40, 463–471. 10.3109/03014460.2013.807878.

3. Goodman, S.R., Daescu, O., Kakhniashvili, D.G., and Zivanic, M. (2013). The proteomics and interactomics of human erythrocytes. Exp Biol Med (Maywood) 238, 509–518. 10.1177/1535370213488474.

4. Bryk, A.H., and Wiśniewski, J.R. (2017). Quantitative Analysis of Human Red Blood Cell Proteome. Journal of Proteome Research 16, 2752–2761. 10.1021/acs.jproteome.7b00025.

5. D’Alessandro, A., Dzieciatkowska, M., Nemkov, T., and Hansen, K.C. (2017). Red blood cell proteomics update: is there more to discover? Blood Transfus 15, 182–187. 10.2450/2017.0293-16.

6. Gautier, E.-F., Leduc, M., Cochet, S., Bailly, K., Lacombe, C., Mohandas, N., Guillonneau, F., El Nemer, W., and Mayeux, P. (2018). Absolute proteome quantification of highly purified populations of circulating reticulocytes and mature erythrocytes. Blood Adv 2, 2646–2657. 10.1182/bloodadvances.2018023515.

7. Haiman, Z.B., Key, A., D’Alessandro, A., and Palsson, B.O. (2025). RBC-GEM: A genome-scale metabolic model for systems biology of the human red blood cell. PLoS Comput Biol 21, e1012109.

8. Dong, S., Wang, Q., Kao, Y.R., Diaz, A., Tasset, I., Kaushik, S., Thiruthuvanathan, V., Zintiridou, A., Nieves, E., Dzieciatkowska, M., et al. (2021). Chaperone-mediated autophagy sustains haematopoietic stem-cell function. Nature 591, 117–123. 10.1038/s41586-020-03129-z.

9. D’Alessandro, A., Fu, X., Kanias, T., Reisz, J.A., Culp-Hill, R., Guo, Y., Gladwin, M.T., Page, G., Kleinman, S., Lanteri, M., et al. (2021). Donor sex, age and ethnicity impact stored red blood cell antioxidant metabolism through mechanisms in part explained by glucose 6-phosphate dehydrogenase levels and activity. Haematologica 106, 1290–1302. 10.3324/haematol.2020.246603.

10. Keele, G.R., Dzieciatkowska, M., Hay, A.M., Vincent, M., O’Connor, C., Stephenson, D., Reisz, J.A., Nemkov, T., Hansen, K.C., Page, G.P., et al. (2025). Genetic architecture of the red blood cell proteome in genetically diverse mice reveals central role of hemoglobin beta cysteine redox status in maintaining circulating glutathione pools. bioRxiv. 10.1101/2025.02.27.640676.

11. Singh, A.K., D’Alessandro, A., Wellendorf, A.M., Gonzalez-Nieto, D., Kofron, M., Dzieciatkowska, M., Mejia, L., Barrio, L.C., Filippi, M.-D., and Cancelas, J.A. (2025). Metabolic adaptation of regenerative hematopoiesis depends on docking-independent mitochondrial connexin 43. Blood 146, 2306–2321. 10.1182/blood.2024028079.

12. Kao, Y.R., Chen, J., Kumari, R., Ng, A., Zintiridou, A., Tatiparthy, M., Ma, Y., Aivalioti, M.M., Moulik, D., Sundaravel, S., et al. (2024). An iron rheostat controls hematopoietic stem cell fate. Cell Stem Cell 31, 378–397.e312. 10.1016/j.stem.2024.01.011.

13. Mills, T.S., Kain, B., Burchill, M.A., Danis, E., Lucas, E.D., Culp-Hill, R., Cowan, C.M., Schleicher, W.E., Patel, S.B., Tran, B.T., et al. (2024). A distinct metabolic and epigenetic state drives trained immunity in HSC-derived macrophages from autoimmune mice. Cell Stem Cell 31, 1630–1649.e1638. 10.1016/j.stem.2024.09.010.

14. Cavalli, G., Justice, J.N., Boyle, K.E., D’Alessandro, A., Eisenmesser, E.Z., Herrera, J.J., Hansen, K.C., Nemkov, T., Stienstra, R., Garlanda, C., et al. (2017). Interleukin 37 reverses the metabolic cost of inflammation, increases oxidative respiration, and improves exercise tolerance. Proc Natl Acad Sci U S A 114, 2313–2318. 10.1073/pnas.1619011114.

15. Xu, P., Chen, C., Zhang, Y., Dzieciatkowska, M., Brown, B.C., Zhang, W., Xie, T., Abdulmalik, O., Song, A., Tong, C., et al. (2022). Erythrocyte transglutaminase-2 combats hypoxia and chronic kidney disease by promoting oxygen delivery and carnitine homeostasis. Cell Metab 34, 299–316.e296. 10.1016/j.cmet.2021.12.019.

16. D’Alessandro, A., Le, K., Lundt, M., Li, Q., Dunkelberger, E.B., Cellmer, T., Worth, A.J., Patil, S., Huston, C., Grier, A., et al. (2024). Functional and multi-omics signatures of mitapivat efficacy upon activation of pyruvate kinase in red blood cells from patients with sickle cell disease. Haematologica 109, 2639–2652. 10.3324/haematol.2023.284831.

17. Peltier, S., Marin, M., Dzieciatkowska, M., Dussiot, M., Roy, M.K., Bruce, J., Leblanc, L., Hadjou, Y., Georgeault, S., Fricot, A., et al. (2025). Proteostasis and metabolic dysfunction characterize a subset of storage-induced senescent erythrocytes targeted for posttransfusion clearance. J Clin Invest 135. 10.1172/jci183099.

18. Sun, B.B., Suhre, K., and Gibson, B.W. (2024). Promises and Challenges of populational Proteomics in Health and Disease. Mol Cell Proteomics 23, 100786. 10.1016/j.mcpro.2024.100786.

19. Assarsson, E., Lundberg, M., Holmquist, G., Björkesten, J., Thorsen, S.B., Ekman, D., Eriksson, A., Rennel Dickens, E., Ohlsson, S., Edfeldt, G., et al. (2014). Homogenous 96-plex PEA immunoassay exhibiting high sensitivity, specificity, and excellent scalability. PLoS One 9, e95192. 10.1371/journal.pone.0095192.

20. Gold, L., Ayers, D., Bertino, J., Bock, C., Bock, A., Brody, E.N., Carter, J., Dalby, A.B., Eaton, B.E., Fitzwater, T., et al. (2010). Aptamer-Based Multiplexed Proteomic Technology for Biomarker Discovery. PLOS ONE 5, e15004. 10.1371/journal.pone.0015004.

21. Sun, B.B., Chiou, J., Traylor, M., Benner, C., Hsu, Y.H., Richardson, T.G., Surendran, P., Mahajan, A., Robins, C., Vasquez-Grinnell, S.G., et al. (2023). Plasma proteomic associations with genetics and health in the UK Biobank. Nature 622, 329–338. 10.1038/s41586-023-06592-6.

22. Dhindsa, R.S., Burren, O.S., Sun, B.B., Prins, B.P., Matelska, D., Wheeler, E., Mitchell, J., Oerton, E., Hristova, V.A., Smith, K.R., et al. (2023). Rare variant associations with plasma protein levels in the UK Biobank. Nature 622, 339–347. 10.1038/s41586-023-06547-x.

23. Eldjarn, G.H., Ferkingstad, E., Lund, S.H., Helgason, H., Magnusson, O.T., Gunnarsdottir, K., Olafsdottir, T.A., Halldorsson, B.V., Olason, P.I., Zink, F., et al. (2023). Large-scale plasma proteomics comparisons through genetics and disease associations. Nature 622, 348–358. 10.1038/s41586-023-06563-x.

24. Suhre, K., Arnold, M., Bhagwat, A.M., Cotton, R.J., Engelke, R., Raffler, J., Sarwath, H., Thareja, G., Wahl, A., DeLisle, R.K., et al. (2017). Connecting genetic risk to disease end points through the human blood plasma proteome. Nature Communications 8, 14357. 10.1038/ncomms14357.

25. Pietzner, M., Wheeler, E., Carrasco-Zanini, J., Cortes, A., Koprulu, M., Wörheide, M.A., Oerton, E., Cook, J., Stewart, I.D., Kerrison, N.D., et al. (2021). Mapping the proteo-genomic convergence of human diseases. Science 374, eabj1541. 10.1126/science.abj1541.

26. Horvath, S. (2013). DNA methylation age of human tissues and cell types. Genome Biol 14, R115. 10.1186/gb-2013-14-10-r115.

27. Hannum, G., Guinney, J., Zhao, L., Zhang, L., Hughes, G., Sadda, S., Klotzle, B., Bibikova, M., Fan, J.-B., Gao, Y., et al. (2013). Genome-wide Methylation Profiles Reveal Quantitative Views of Human Aging Rates. Molecular Cell 49, 359–367. 10.1016/j.molcel.2012.10.016.

28. Levine, M.E., Lu, A.T., Quach, A., Chen, B.H., Assimes, T.L., Bandinelli, S., Hou, L., Baccarelli, A.A., Stewart, J.D., Li, Y., et al. (2018). An epigenetic biomarker of aging for lifespan and healthspan. Aging (Albany NY) 10, 573–591. 10.18632/aging.101414.

29. Lu, A.T., Quach, A., Wilson, J.G., Reiner, A.P., Aviv, A., Raj, K., Hou, L., Baccarelli, A.A., Li, Y., Stewart, J.D., et al. (2019). DNA methylation GrimAge strongly predicts lifespan and healthspan. Aging 11, 303–327. 10.18632/aging.101684.

30. Liu, Z., Leung, D., Thrush, K., Zhao, W., Ratliff, S., Tanaka, T., Schmitz, L.L., Smith, J.A., Ferrucci, L., and Levine, M.E. (2020). Underlying features of epigenetic aging clocks in vivo and in vitro. Aging Cell 19, e13229. 10.1111/acel.13229.

31. Trumpff, C., Marsland, A.L., Basualto-Alarcón, C., Martin, J.L., Carroll, J.E., Sturm, G., Vincent, A.E., Mosharov, E.V., Gu, Z., Kaufman, B.A., and Picard, M. (2019). Acute psychological stress increases serum circulating cell-free mitochondrial DNA. Psychoneuroendocrinology 106, 268–276. 10.1016/j.psyneuen.2019.03.026.

32. Josephson, C.D., Glynn, S., Mathew, S., Birch, R., Bakkour, S., Baumann Kreuziger, L., Busch, M.P., Chapman, K., Dinardo, C., Hendrickson, J., et al. (2022). The Recipient Epidemiology and Donor Evaluation Study-IV-Pediatric (REDS-IV-P): A research program striving to improve blood donor safety and optimize transfusion outcomes across the lifespan. Transfusion 62, 982–999. 10.1111/trf.16869.

33. Endres-Dighe, S.M., Guo, Y., Kanias, T., Lanteri, M., Stone, M., Spencer, B., Cable, R.G., Kiss, J.E., Kleinman, S., Gladwin, M.T., et al. (2019). Blood, sweat, and tears: Red Blood Cell-Omics study objectives, design, and recruitment activities. Transfusion 59, 46–56. 10.1111/trf.14971.

34. Page, G.P., Kanias, T., Guo, Y.J., Lanteri, M.C., Zhang, X., Mast, A.E., Cable, R.G., Spencer, B.R., Kiss, J.E., Fang, F., et al. (2021). Multiple-ancestry genome-wide association study identifies 27 loci associated with measures of hemolysis following blood storage. J Clin Invest 131. 10.1172/jci146077.

35. D’Alessandro, A., Keele, G.R., Hay, A., Nemkov, T., Earley, E.J., Stephenson, D., Vincent, M., Deng, X., Stone, M., Dzieciatkowska, M., et al. (2025). Ferroptosis regulates hemolysis in stored murine and human red blood cells. Blood 145, 765–783. 10.1182/blood.2024026109.

36. Nemkov, T., Stephenson, D., Earley, E.J., Keele, G.R., Hay, A., Key, A., Haiman, Z.B., Erickson, C., Dzieciatkowska, M., Reisz, J.A., et al. (2024). Biological and genetic determinants of glycolysis: Phosphofructokinase isoforms boost energy status of stored red blood cells and transfusion outcomes. Cell Metabolism. 10.1016/j.cmet.2024.06.007.

37. Dreischer, P., Duszenko, M., Stein, J., and Wieder, T. (2022). Eryptosis: Programmed Death of Nucleus-Free, Iron-Filled Blood Cells. Cells 11. 10.3390/cells11030503.

38. Peltier, S., Marin, M., Dzieciatkowska, M., Dussiot, M., Roy, M.K., Bruce, J., Leblanc, L., Hadjou, Y., Georgeault, S., Fricot, A., et al. (2025). Proteostasis and metabolic dysfunction characterize a subset of storage-induced senescent erythrocytes targeted for post-transfusion clearance. J Clin Invest. 10.1172/jci183099.

39. Brodersen, T., Rostgaard, K., Lau, C.J., Juel, K., Erikstrup, C., Nielsen, K.R., Ostrowski, S.R., Titlestad, K., Saekmose, S.G., Pedersen, O.B.V., and Hjalgrim, H. (2023). The healthy donor effect and survey participation, becoming a donor and donor career. Transfusion 63, 143–155. 10.1111/trf.17190.

40. Roubinian, N.H., Reese, S.E., Qiao, H., Plimier, C., Fang, F., Page, G.P., Cable, R.G., Custer, B., Gladwin, M.T., Goel, R., et al. (2022). Donor genetic and nongenetic factors affecting red blood cell transfusion effectiveness. JCI Insight 7. 10.1172/jci.insight.152598.

41. Francis, R.O., D’Alessandro, A., Eisenberger, A., Soffing, M., Yeh, R., Coronel, E., Sheikh, A., Rapido, F., La Carpia, F., Reisz, J.A., et al. (2020). Donor glucose-6-phosphate dehydrogenase deficiency decreases blood quality for transfusion. The Journal of Clinical Investigation 130, 2270–2285. 10.1172/JCI133530.

42. Remy, K.E., Hall, M.W., Cholette, J., Juffermans, N.P., Nicol, K., Doctor, A., Blumberg, N., Spinella, P.C., Norris, P.J., Dahmer, M.K., and Muszynski, J.A. (2018). Mechanisms of red blood cell transfusion-related immunomodulation. Transfusion 58, 804–815. 10.1111/trf.14488.

43. Yang, L., You, J., Yang, X., Jiao, R., Xu, J., Zhang, Y., Mi, W., Zhu, L., Ye, Y., Ren, R., et al. (2025). ACSS2 drives senescence-associated secretory phenotype by limiting purine biosynthesis through PAICS acetylation. Nat Commun 16, 2071. 10.1038/s41467-025-57334-3.

44. Hwang, I.K., Kim, D.W., Jung, J.Y., Yoo, K.Y., Cho, J.H., Kwon, O.S., Kang, T.C., Choi, S.Y., Kim, Y.S., and Won, M.H. (2005). Age-dependent changes of pyridoxal phosphate synthesizing enzymes immunoreactivities and activities in the gerbil hippocampal CA1 region. Mech Ageing Dev 126, 1322–1330. 10.1016/j.mad.2005.08.007.

45. Antia, M., Baneyx, G., Kubow, K.E., and Vogel, V. (2008). Fibronectin in aging extracellular matrix fibrils is progressively unfolded by cells and elicits an enhanced rigidity response. Faraday Discuss 139, 229–249; discussion 309-225, 419-220. 10.1039/b718714a.

46. Hager, K., Felicetti, M., Seefried, G., and Platt, D. (1994). Fibrinogen and aging. Aging (Milano) 6, 133–138. 10.1007/bf03324226.

47. Klonoff-Cohen, H., Barrett-Connor, E.L., and Edelstein, S.L. (1992). Albumin levels as a predictor of mortality in the healthy elderly. J Clin Epidemiol 45, 207–212. 10.1016/0895-4356(92)90080-7.

48. Di Castelnuovo, A., Iacoviello, L., and Violi, F. (2025). Albumin and mortality: addressing critiques and reaffirming findings. eClinicalMedicine 80. 10.1016/j.eclinm.2024.102948.

49. Wang, K., Shang, Y., and Dou, F. (2018). Brain Aging: Hsp90 and Neurodegenerative Diseases. Adv Exp Med Biol 1086, 93–103. 10.1007/978-981-13-1117-8_6.

50. Hoy, W.E., Hughson, M.D., Kopp, J.B., Mott, S.A., Bertram, J.F., and Winkler, C.A. (2015). APOL1 Risk Alleles Are Associated with Exaggerated Age-Related Changes in Glomerular Number and Volume in African-American Adults: An Autopsy Study. J Am Soc Nephrol 26, 3179–3189. 10.1681/asn.2014080768.

51. Mukamal, K.J., Tremaglio, J., Friedman, D.J., Ix, J.H., Kuller, L.H., Tracy, R.P., and Pollak, M.R. (2016). *APOL1* Genotype, Kidney and Cardiovascular Disease, and Death in Older Adults. Arteriosclerosis, Thrombosis, and Vascular Biology 36, 398–403. doi:10.1161/ATVBAHA.115.305970.

52. Sniderman, A.D., Islam, S., McQueen, M., Pencina, M., Furberg, C.D., Thanassoulis, G., and Yusuf, S. (2016). Age and Cardiovascular Risk Attributable to Apolipoprotein B, Low-Density Lipoprotein Cholesterol or Non-High-Density Lipoprotein Cholesterol. J Am Heart Assoc 5. 10.1161/jaha.116.003665.

53. Gross, L. (2006). The key to longevity? Having long-lived parents is a good start. PLoS Biol 4, e119. 10.1371/journal.pbio.0040119.

54. Molteni, R., Fiumara, M., Campochiaro, C., Alfieri, R., Pacini, G., Licari, E., Tomelleri, A., Diral, E., Varesi, A., Weber, A., et al. (2025). Mechanisms of hematopoietic clonal dominance in VEXAS syndrome. Nature Medicine 31, 1911–1924. 10.1038/s41591-025-03623-9.

55. Murphy, W.G. (2014). The sex difference in haemoglobin levels in adults - mechanisms, causes, and consequences. Blood Rev 28, 41–47. 10.1016/j.blre.2013.12.003.

56. Gold, E.B. (2011). The timing of the age at which natural menopause occurs. Obstet Gynecol Clin North Am 38, 425–440. 10.1016/j.ogc.2011.05.002.

57. Hazegh, K., Fang, F., Bravo, M.D., Tran, J.Q., Muench, M.O., Jackman, R.P., Roubinian, N., Bertolone, L., D’Alessandro, A., Dumont, L., et al. (2021). Blood donor obesity is associated with changes in red blood cell metabolism and susceptibility to hemolysis in cold storage and in response to osmotic and oxidative stress. Transfusion 61, 435–448. 10.1111/trf.16168.

58. Kanias, T., Lanteri, M.C., Page, G.P., Guo, Y., Endres, S.M., Stone, M., Keating, S., Mast, A.E., Cable, R.G., Triulzi, D.J., et al. (2017). Ethnicity, sex, and age are determinants of red blood cell storage and stress hemolysis: results of the REDS-III RBC-Omics study. Blood Adv 1, 1132–1141. 10.1182/bloodadvances.2017004820.

59. Roumenina, L.T., Bartolucci, P., and Pirenne, F. (2019). The role of Complement in Post-Transfusion Hemolysis and Hyperhemolysis Reaction. Transfusion Medicine Reviews 33, 225–230. 10.1016/j.tmrv.2019.09.007.

60. Reisz, J.A., Earley, E.J., Nemkov, T., Key, A., Stephenson, D., Keele, G.R., Dzieciatkowska, M., Spitalnik, S.L., Hod, E.A., Kleinman, S., et al. (2025). Arginine metabolism is a biomarker of red blood cell and human aging. Aging Cell 24, e14388. 10.1111/acel.14388.

61. Nemkov, T., Stephenson, D., Erickson, C., Dzieciatkowska, M., Key, A., Moore, A., Earley, E.J., Page, G.P., Lacroix, I.S., Stone, M., et al. (2024). Regulation of kynurenine metabolism by blood donor genetics and biology impacts red cell hemolysis in vitro and in vivo. Blood 143, 456–472. 10.1182/blood.2023022052.

62. Guntur, V.P., Nemkov, T., de Boer, E., Mohning, M.P., Baraghoshi, D., Cendali, F.I., San-Millán, I., Petrache, I., and D’Alessandro, A. (2022). Signatures of Mitochondrial Dysfunction and Impaired Fatty Acid Metabolism in Plasma of Patients with Post-Acute Sequelae of COVID-19 (PASC). Metabolites 12, 1026.

63. Basak, A., and Sankaran, V.G. (2016). Regulation of the fetal hemoglobin silencing factor BCL11A. Ann N Y Acad Sci 1368, 25–30. 10.1111/nyas.13024.

64. Liu, J., Chen, T., Zhao, Y., Ding, Z., Ge, W., and Zhang, J. (2022). Blood donation improves skin aging through the reduction of iron deposits and the increase of TGF-β1 in elderly skin. Mechanisms of Ageing and Development 205, 111687. 10.1016/j.mad.2022.111687.

65. Hod, E.A., Brittenham, G.M., Bitan, Z.C., Feit, Y., Gaelen, J.I., La Carpia, F., Sandoval, L.A., Zhou, A.T., Soffing, M., Mintz, A., et al. (2022). A randomized trial of blood donor iron repletion on red cell quality for transfusion and donor cognition and well-being. Blood 140, 2730–2739. 10.1182/blood.2022017288.

66. Hod, E.A., Habeck, C., Zhuang, H., Dimov, A., Spincemaille, P., Kessler, D., Bitan, Z.C., Feit, Y., Fliginger, D., Stone, E.F., et al. (2025). Effects of iron repletion on brain iron content, myelination, neural network activation, and cognition. JCI Insight. 10.1172/jci.insight.194442.

67. Rinalducci, S., Ferru, E., Blasi, B., Turrini, F., and Zolla, L. (2012). Oxidative stress and caspase-mediated fragmentation of cytoplasmic domain of erythrocyte band 3 during blood storage. Blood Transfus 10 *Suppl 2*, s55–62. 10.2450/2012.009s.

68. Issaian, A., Hay, A., Dzieciatkowska, M., Roberti, D., Perrotta, S., Darula, Z., Redzic, J., Busch, M.P., Page, G.P., Rogers, S.C., et al. (2021). The interactome of the N-terminus of band 3 regulates red blood cell metabolism and storage quality. Haematologica. 10.3324/haematol.2020.278252.

69. Nemkov, T., Stefanoni, D., Bordbar, A., Issaian, A., Palsson, B.O., Dumont, L.J., Hay, A.M., Song, A., Xia, Y., Redzic, J.S., et al. (2020). Blood donor exposome and impact of common drugs on red blood cell metabolism. JCI Insight. 10.1172/jci.insight.146175.

70. Mebratu, Y.A., Soni, S., Rosas, L., Rojas, M., Horowitz, J.C., and Nho, R. (2023). The aged extracellular matrix and the profibrotic role of senescence-associated secretory phenotype. Am J Physiol Cell Physiol 325, C565–c579. 10.1152/ajpcell.00124.2023.

71. López-Otín, C., Blasco, M.A., Partridge, L., Serrano, M., and Kroemer, G. (2023). Hallmarks of aging: An expanding universe. Cell 186, 243–278. 10.1016/j.cell.2022.11.001.

72. Hyde, M.K., Masser, B.M., Thorpe, R., Philip, A.A., Salmon, A., Scott, T.L., and Davison, T.E. (2025). Rethinking the role of older donors in a sustainable blood supply. Transfusion 65, 758–766. 10.1111/trf.18190.

73. D’Alessandro, A., Culp-Hill, R., Reisz, J.A., Anderson, M., Fu, X., Nemkov, T., Gehrke, S., Zheng, C., Kanias, T., Guo, Y., et al. (2019). Heterogeneity of blood processing and storage additives in different centers impacts stored red blood cell metabolism as much as storage time: lessons from REDS-III-Omics. Transfusion 59, 89–100. 10.1111/trf.14979.

74. Lanteri, M.C., Kanias, T., Keating, S., Stone, M., Guo, Y., Page, G.P., Brambilla, D.J., Endres-Dighe, S.M., Mast, A.E., Bialkowski, W., et al. (2019). Intradonor reproducibility and changes in hemolytic variables during red blood cell storage: results of recall phase of the REDS-III RBC-Omics study. Transfusion 59, 79–88. 10.1111/trf.14987.

75. Nemkov, T., Yoshida, T., Nikulina, M., and D’Alessandro, A. (2022). High-Throughput Metabolomics Platform for the Rapid Data-Driven Development of Novel Additive Solutions for Blood Storage. Frontiers in Physiology 13. 10.3389/fphys.2022.833242.

76. Thomas, T., Stefanoni, D., Dzieciatkowska, M., Issaian, A., Nemkov, T., Hill, R.C., Francis, R.O., Hudson, K.E., Buehler, P.W., Zimring, J.C., et al. (2020). Evidence for structural protein damage and membrane lipid remodeling in red blood cells from COVID-19 patients. medRxiv. 10.1101/2020.06.29.20142703.

77. Reisz, J.A., Zheng, C., D’Alessandro, A., and Nemkov, T. (2019). Untargeted and Semi-targeted Lipid Analysis of Biological Samples Using Mass Spectrometry-Based Metabolomics. Methods Mol Biol 1978, 121–135. 10.1007/978-1-4939-9236-2_8.

78. Stephenson, D., Nemkov, T., Qadri, S.M., Sheffield, W.P., and D’Alessandro, A. (2022). Inductively-Coupled Plasma Mass Spectrometry-Novel Insights From an Old Technology Into Stressed Red Blood Cell Physiology. Front Physiol 13, 828087. 10.3389/fphys.2022.828087.

79. Moore, A., Busch, M.P., Dziewulska, K., Francis, R.O., Hod, E.A., Zimring, J.C., D’Alessandro, A., and Page, G.P. (2022). Genome-wide metabolite quantitative trait loci analysis (mQTL) in red blood cells from volunteer blood donors. J Biol Chem 298, 102706. 10.1016/j.jbc.2022.102706.

80. Nemkov, T., Stephenson, D., Earley, E.J., Keele, G.R., Hay, A., Key, A., Haiman, Z.B., Erickson, C., Dzieciatkowska, M., Reisz, J.A., et al. (2024). Biological and genetic determinants of glycolysis: Phosphofructokinase isoforms boost energy status of stored red blood cells and transfusion outcomes. Cell Metab. 10.1016/j.cmet.2024.06.007.

81. Stephenson, D., Keele, G.R., Hay, A., Dzieciatkowska, M., Reisz, J.A., Haiman, Z.B., Moore, A.L., Nemkov, T., Deng, X., Stone, M., et al. (2025). GPX4 regulates lipid peroxidation and ferroptosis of stored red blood cells. Blood Red Cells & Iron, 100020. 10.1016/j.brci.2025.100020.

82. Guo, Y., Busch, M.P., Seielstad, M., Endres-Dighe, S., Westhoff, C.M., Keating, B., Hoppe, C., Bordbar, A., Custer, B., Butterworth, A.S., et al. (2019). Development and evaluation of a transfusion medicine genome wide genotyping array. Transfusion 59, 101–111. 10.1111/trf.15012.

83. Delaneau, O., Coulonges, C., and Zagury, J.-F. (2008). Shape-IT: new rapid and accurate algorithm for haplotype inference. BMC Bioinformatics 9, 540. 10.1186/1471-2105-9-540.

84. Howie, B., Marchini, J., and Stephens, M. (2011). Genotype Imputation with Thousands of Genomes. G3 Genes|Genomes|Genetics 1, 457–470. 10.1534/g3.111.001198.

85. Zheng, X., Levine, D., Shen, J., Gogarten, S.M., Laurie, C., and Weir, B.S. (2012). A high-performance computing toolset for relatedness and principal component analysis of SNP data. Bioinformatics 28, 3326–3328. 10.1093/bioinformatics/bts606.

86. Aulchenko, Y.S., Struchalin, M.V., and van Duijn, C.M. (2010). ProbABEL package for genome-wide association analysis of imputed data. BMC Bioinformatics 11, 134. 10.1186/1471-2105-11-134.

87. Perry, J.A., Gaynor, B.J., Mitchell, B.D., and O’Connell, J.R. (2021). An Omics Analysis Search and Information System (OASIS) for Enabling Biological Discovery in the Old Order Amish. bioRxiv, 2021.2005.2002.442370. 10.1101/2021.05.02.442370.

88. https://biolincc.nhlbi.nih.gov/studies/reds_iii/.

89. Karafin, M.S., Bruhn, R., Westlake, M., Sullivan, M.T., Bialkowski, W., Edgren, G., Roubinian, N.H., Hauser, R.G., Kor, D.J., Fleischmann, D., et al. (2017). Demographic and epidemiologic characterization of transfusion recipients from four US regions: evidence from the REDS-III recipient database. Transfusion 57, 2903–2913. 10.1111/trf.14370.

90. Endres-Dighe, S.M., Guo, Y., Kanias, T., Lanteri, M., Stone, M., Spencer, B., Cable, R.G., Kiss, J.E., Kleinman, S., Gladwin, M.T., et al. (2019). Blood, sweat, and tears: Red Blood Cell-Omics study objectives, design, and recruitment activities. Transfusion 59, 46–56. 10.1111/trf.14971.

91. Nemkov, T., Yoshida, T., Nikulina, M., and D’Alessandro, A. (2022). High-Throughput Metabolomics Platform for the Rapid Data-Driven Development of Novel Additive Solutions for Blood Storage. Front Physiol 13, 833242. 10.3389/fphys.2022.833242.

92. Nemkov, T., Reisz, J.A., Gehrke, S., Hansen, K.C., and D’Alessandro, A. (2019). High-Throughput Metabolomics: Isocratic and Gradient Mass Spectrometry-Based Methods. Methods Mol Biol 1978, 13–26. 10.1007/978-1-4939-9236-2_2.

93. Stefanoni, D., Shin, H.K.H., Baek, J.H., Champagne, D.P., Nemkov, T., Thomas, T., Francis, R.O., Zimring, J.C., Yoshida, T., Reisz, J.A., et al. (2020). Red blood cell metabolism in Rhesus macaques and humans: comparative biology of blood storage. Haematologica 105, 2174–2186. 10.3324/haematol.2019.229930.

94. Nemkov, T., Hansen, K.C., and D’Alessandro, A. (2017). A three-minute method for high-throughput quantitative metabolomics and quantitative tracing experiments of central carbon and nitrogen pathways. Rapid Commun Mass Spectrom 31, 663–673. 10.1002/rcm.7834.

