## Supplementary Figures for "A population-scale Red Blood Cell proteome atlas of 13,000 donors uncovers genetically encoded aging clocks predicting hemolysis, transfusion efficacy, and donor activity a decade later"

**SUPPLEMENTARY TABLE 1** .......... XLSX

### SUPPLEMENTARY FIGURES

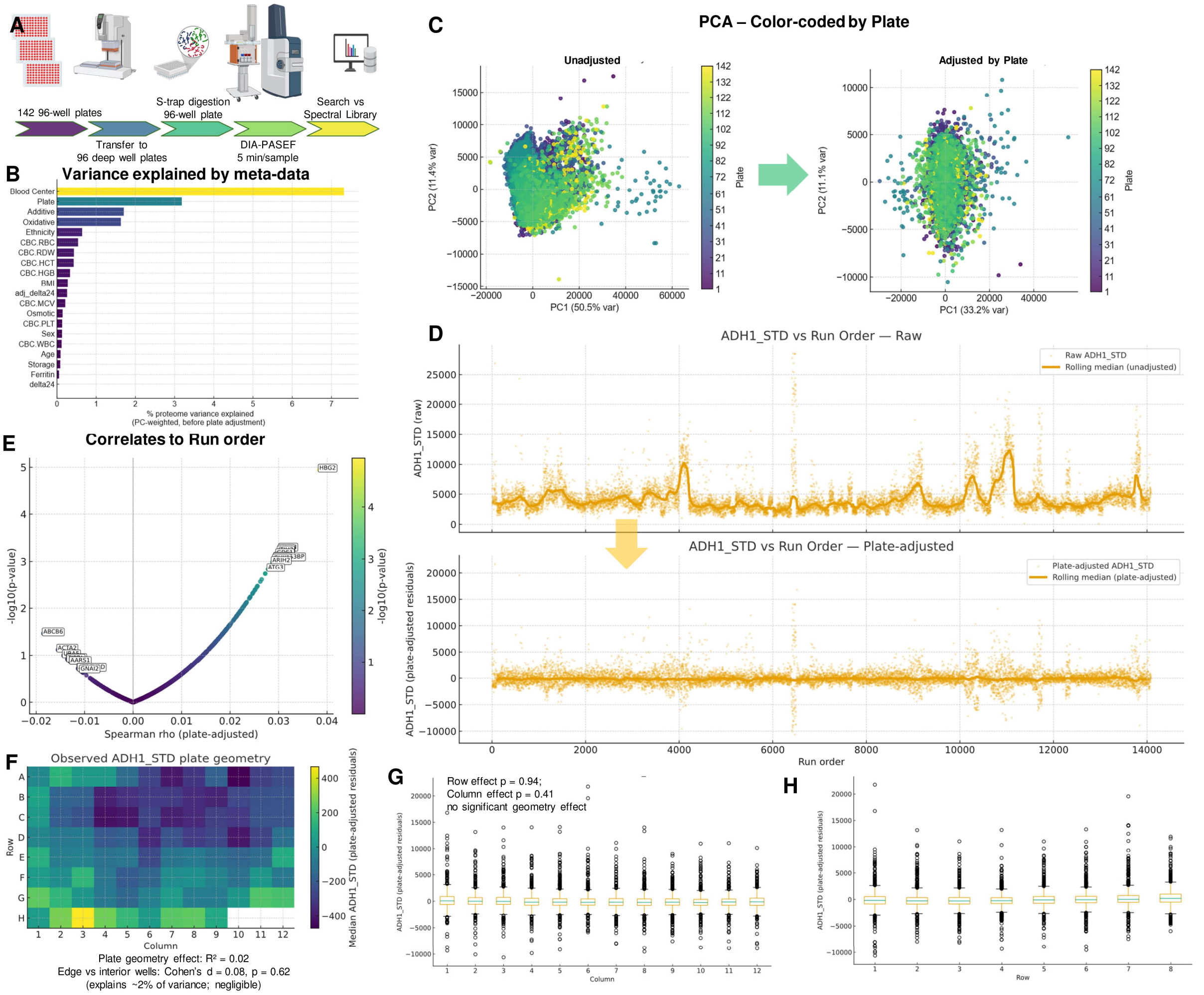

**Supplementary Figure 1. Quality control assessment of the proteomics dataset.** Overview of the proteomics workflow **(A)**. A total of 142 ninety-six–well plates were processed, including sample transfer to 96-deep-well plates, multiplexed S-Trap digestion on plate, and DIA-PASEF analysis (5-min gradients) on a timsTOF instrument, followed by search against a pre-built spectral-library obtained from pooling of an aliquot from all ~13 thousand samples. Pre-adjustment determination of percentage of proteome variance explained by sample-level metadata, computed using a PC-weighted variance decomposition approach **(B)**. Blood Center was the predominant contributor to total variance, followed by Plate and Additive, while biological and storage/hemolysis variables accounted for smaller proportions of variance. Based on this observation, further analyses focusing on biological variables in the main manuscript included an adjustment step for plate, blood center and additive (processing and pre-processing variables). Principal component analysis (PCA) of the full dataset before (left) and after (right) plate adjustment **(C)**. Samples are colored by plate number on a continuous viridis scale. Prior to adjustment, substantial plate-level structure is evident, whereas plate-centering markedly reduces technical clustering while preserving biological variability. Minimal association was observed between protein abundances and run order after plate-based adjustment **(D)**. Each point represents a protein, with the x-axis showing Spearman correlation between abundance and run order and the y-axis showing –log10(p-value). Proteins exhibiting significant run-order–dependent trends are highlighted. Raw (top) and plate-adjusted (bottom) intensities of the internal standard *yeast* ADH1 plotted across run order **(E)**. The rolling median (solid line) highlights substantial run-order drift in the raw data, which is effectively removed after plate adjustment. Heatmap of ADH1_STD intensities across plate geometry (rows A–H, columns 1–12) **(F)**. Plate-position effects were minimal after adjustment ($R^2 = 0.02$ for plate geometry; Cohen's d = 0.08 for edge vs interior wells), indicating negligible bias attributable to well location. Distribution of ADH1_STD (plate-adjusted) across plate columns **(G)** and rows **(H)**. No significant row or column effects were observed (row effect $p = 0.94$; column effect $p = 0.41$), confirming the absence of systematic spatial bias.

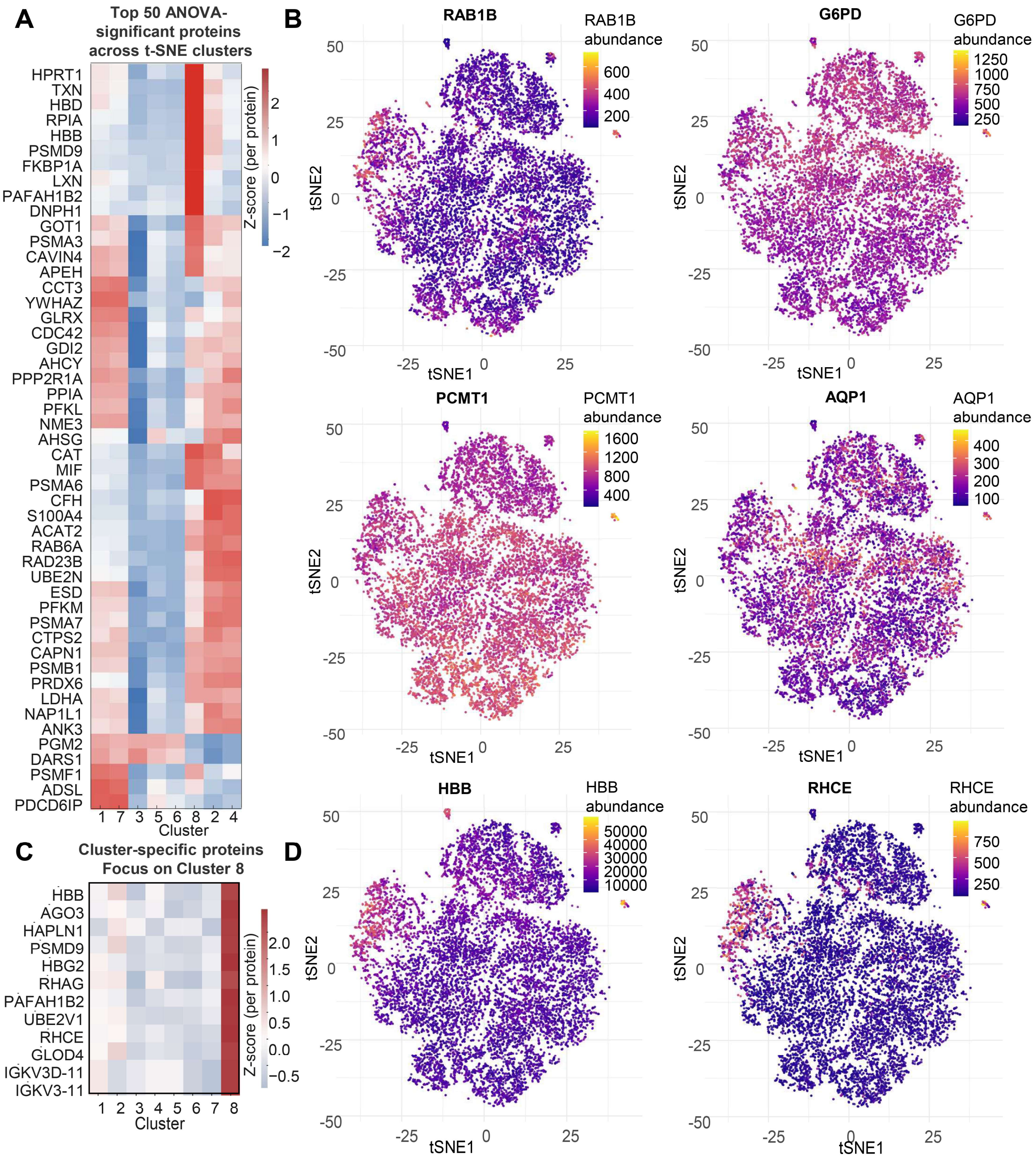

**Supplementary Figure 2. Proteomic features underlying t-SNE clusters and spatial expression patterns of key RBC proteins.** Heatmap of the top 50 significant proteins (FDR-adjusted ANOVA) across t-SNE–defined donor clusters. Z-scored abundances highlight proteins with strong cluster-specific enrichment, including oxidative stress regulators (HPRT1, TXN, PRDX6), cytoskeletal components (HBD, HBB, RHAG), chaperones (CCT3, PSMD9), and metabolic enzymes (LDHA, PGK1, PFKM), reflecting heterogeneous proteomic substructures within the donor population **(A)**. Spatial mapping of individual protein abundances over the t-SNE embedding. Each dot represents a donor, colored by protein intensity **(B–D)**. Heatmap of cluster-specific proteins with a focus on Cluster 8, which shows pronounced enrichment for hemoglobin subunits (HBB, HBG2, HBA1/HBA2), proteasomal components (PSMD9, UBE2V1), cytoskeletal regulators (RHCE, GLUD4), and immunoglobulin fragments (IGKV3D-11, IGKV3-11). These features define Cluster 8 as a hemoglobin-rich, oxidative-stress-linked proteomic state **(C)**.

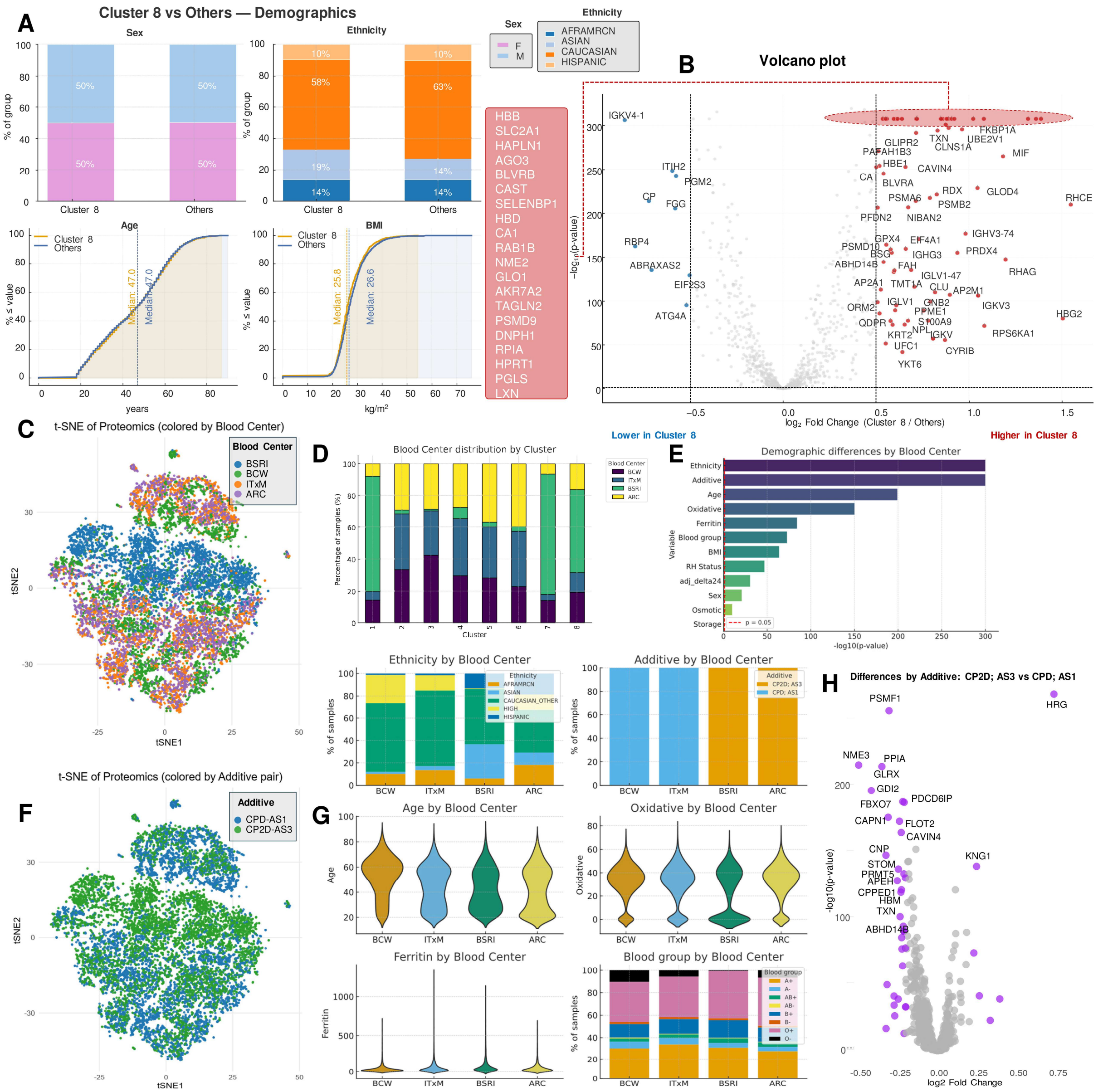

**Supplementary Figure 3. Demographic and processing characteristics distinguishing Cluster 8 from all other proteomic clusters.** Demographic profiles of Cluster 8 versus all other donors **(A)**. Sex, age and BMI distributions are balanced, while ethnicity analysis shows modest enrichment for donors of Asian descent in Cluster 8 compared to the other clusters combined. Volcano plot of most strongly enriched in Cluster 8 (top), including hemoglobin subunits (HBB, HBD, HBG2), membrane proteins (SLC2A1, RHAG), chaperones (HAPLN1, PSMD9), and redox enzymes (PRDX1, TXN, BLVRB), consistent with the hemoglobin-rich, oxidative-metabolic signature of this cluster **(B)**. Results show significantly elevated hemoglobin subunits, proteasome components, antioxidant enzymes, and membrane proteins in Cluster 8 (right side), and lower abundance of immunoglobulins, complement components, and lipid-binding proteins on the left side.
t-SNE visualization of the proteomic landscape colored by Blood Center **(C)**, revealing overlapping but distinct regional enrichment patterns across centers (ITxM, BSRI, BCW, ARC). Distributions of Blood Center, Ethnicity, and Additive Solution across clusters. Certain centers (e.g., ARC, ITxM) contribute proportionally more donors to Cluster 8 **(D)**. Additive usage (CPD-AS1 vs CP2D-AS3) varies substantially by center, with BCW and ITxM adopting CPD;AS-1, while BSRI and ARC using CP2D;AS-3, motivating the need for center-adjusted analyses. Demographic differences across Blood Centers, with ethnicity, additive solution, age, BMI, ferritin, and hemolysis metrics showing center-specific biases **(E)**. These differences justify the inclusion of Blood Center and additives as a covariate in all downstream models through the present study. t-SNE embeddings colored by additive pair (CPD-AS1 vs CP2D-AS3), showing partial segregation consistent with additive-dependent

proteomic shifts **(F)**. Violin plots showing Blood Center–specific distributions for age, oxidative hemolysis, ferritin, and blood group. These variables exhibit marked inter-center variability, underscoring structural differences in donor populations and processing pipelines across centers **(G)**. Volcano plot showing proteomic differences by additive solution (CPD-AS1 vs CP2D-AS3) **(H)**.

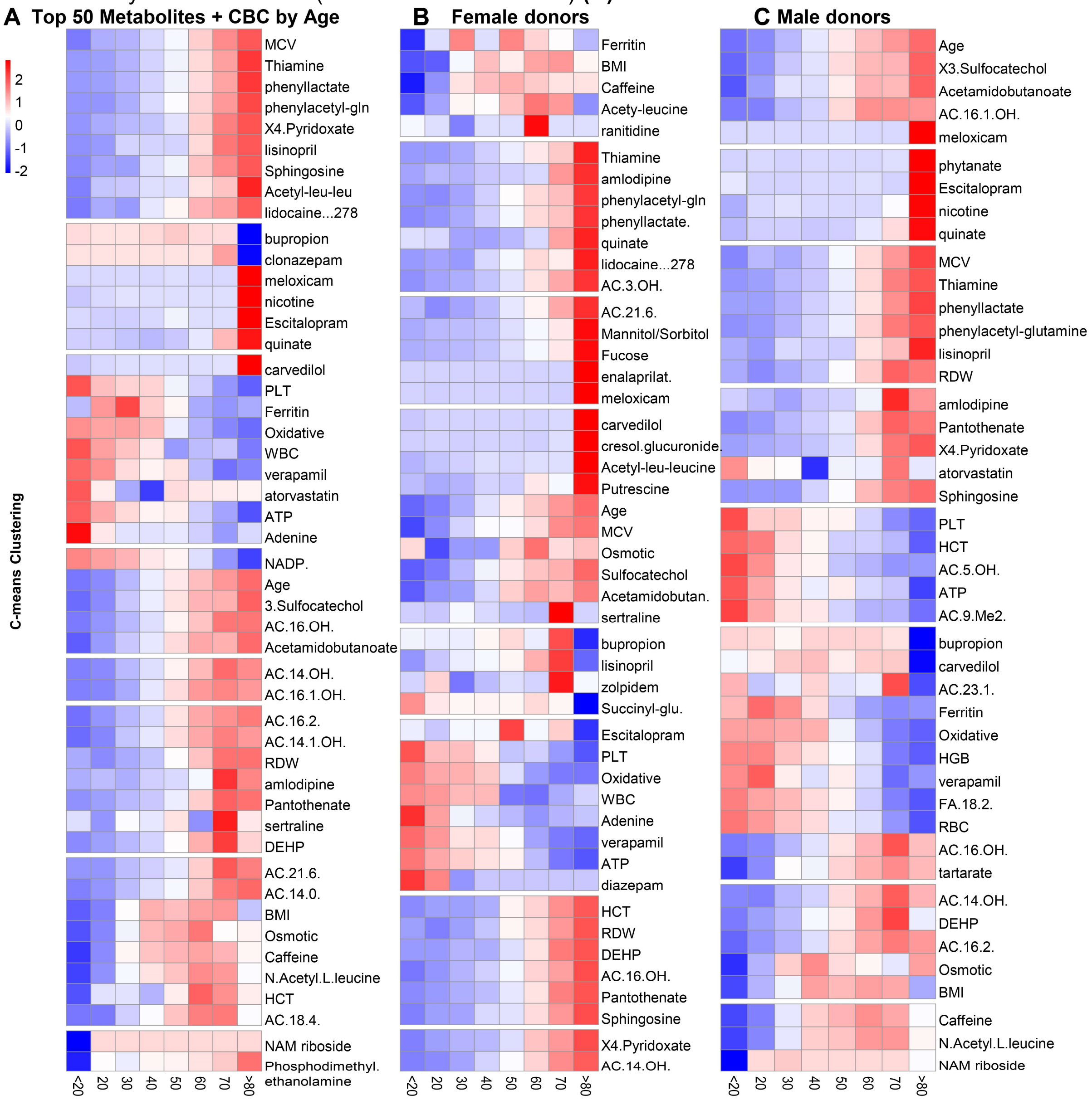

**Supplementary Figure 4. Age-associated metabolomic signatures and CBC features stratified by sex reveal sex-specific metabolic aging trajectories.** Heatmap of the top 50 age-associated metabolites and CBC parameters, identified by fuzzy c-means clustering by age decade buckets for the whole population **(A),** or separately for female or male donors **(B-C**, respectively). This meta-analysis of metabolomics data on the same 13,091 samples from the REDS RBC Omics Index cohort is based upon data reported in Reisz et al. Aging Cell 2025. PMCID: PMC11822668; doi: 10.1111/acel.14388.

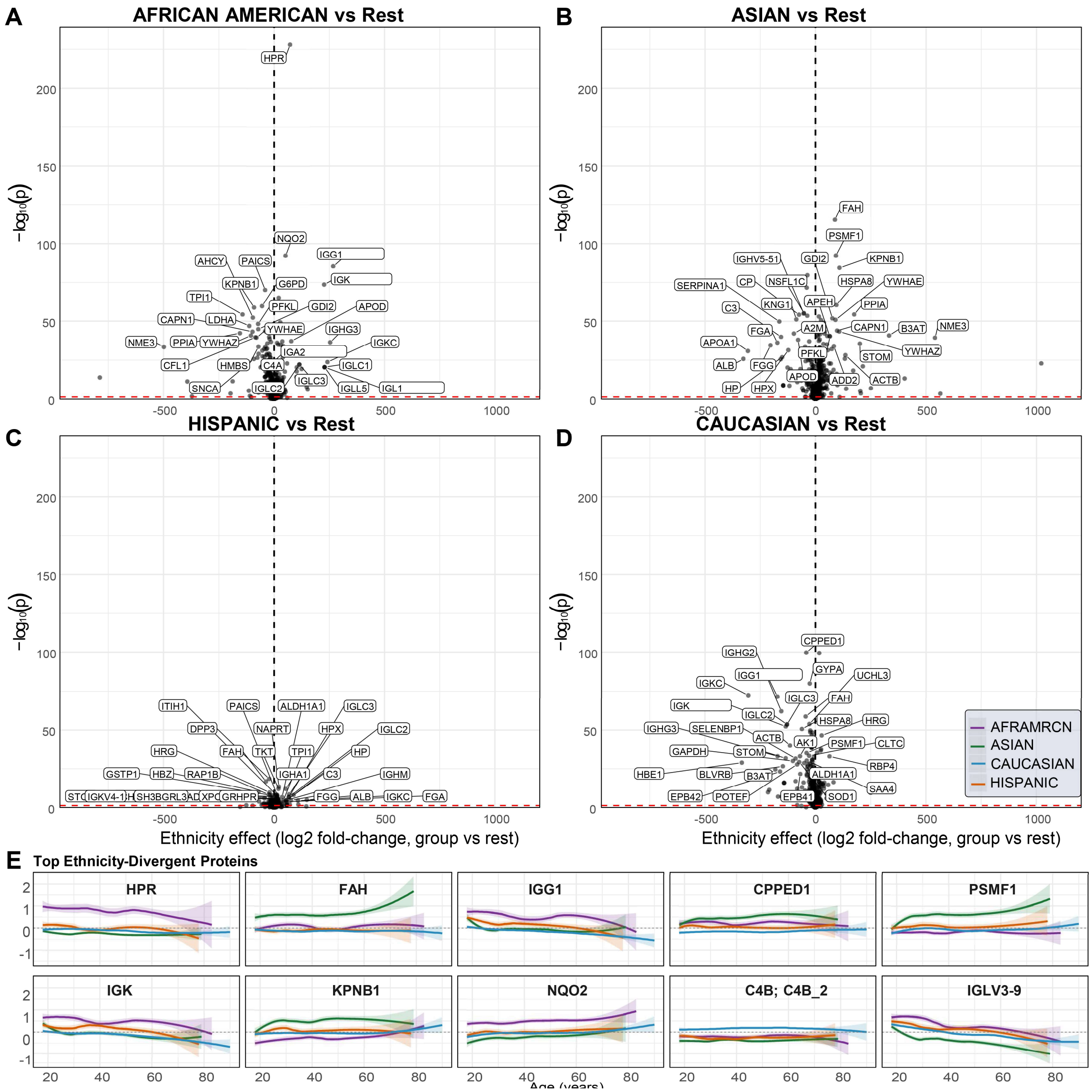

**Supplementary Figure 5. Ethnicity-associated proteomic differences and age-dependent expression trajectories of top ancestry-divergent proteins.** Volcano plots showing ethnicity-associated proteomic differences based on genetic ancestry (Asian, African American, Hispanic or Caucasian) **(A–D)**. Labels highlight proteins surpassing the statistical threshold (horizontal dashed line), with x-axis values indicating ethnicity effect size (log2 fold-change). Age trajectories of the top ethnicity-divergent proteins across all groups (African American, Asian, Caucasian, Hispanic), illustrating how ancestry-associated protein differences evolve over the lifespan **(E)**. These analyses reveal that while ethnicity-associated proteomic signatures are strong at baseline, many proteins show age-dependent amplification or compression of inter-group differences, demonstrating that ancestry and aging jointly shape RBC proteomic architecture.

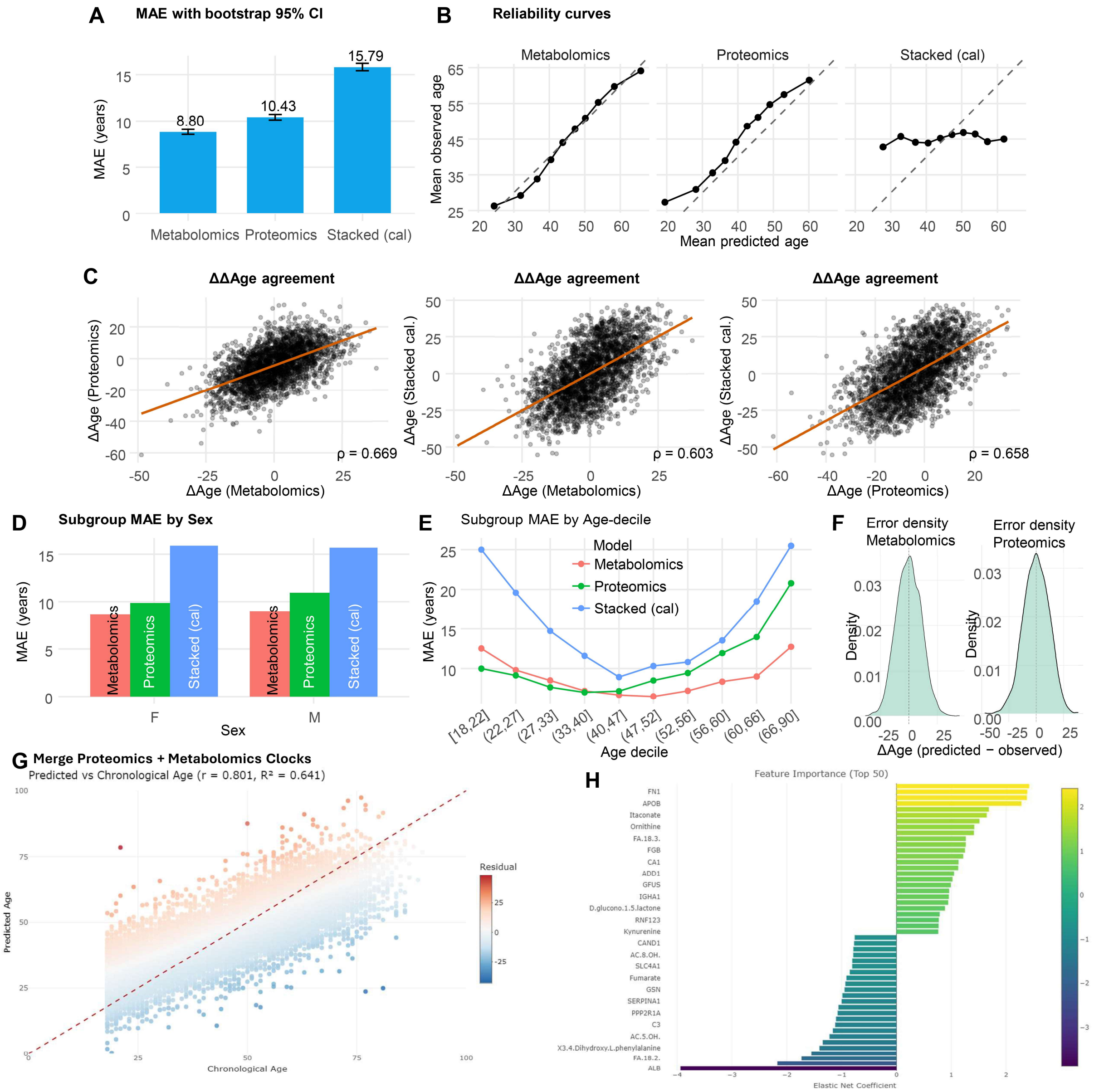

**Supplementary Figure 6. Performance, calibration, and feature contributions of proteomics and metabolomics aging clocks and their stacked ensemble model.** Mean absolute error (MAE) with bootstrap-corrected 95% confidence intervals for the metabolomics, proteomics, and stacked (calibrated) aging clocks **(A)**. Metabolomics achieves the lowest MAE (8.80 years), followed by proteomics (10.43 years), while the stacked model shows modestly higher error (15.79 years) prior to calibration adjustment. Reliability (calibration) curves showing agreement between predicted and observed chronological age, though the stacked model shows negligible residual reliability advantage over independent models **(B)**. Pairwise agreement between ΔAge residuals (Predicted – Chronological Age) across models **(C)**. Strong correlations are observed between metabolomics and proteomics clocks (ρ = 0.669), metabolomics and stacked (ρ = 0.603), and proteomics and stacked (ρ = 0.658), highlighting consistent biological aging signatures across omics layers. Subgroup MAE by sex **(D)**. All models perform similarly in females and males, with metabolomics consistently outperforming proteomics and stacked approaches across both sexes. Subgroup MAE by age-decade **(E)**. Metabolomics and proteomics both show lowest error in mid-life (40–60 years) and higher error at the older extreme of age, consistent with the nonlinear nature of biological aging, healthy donor bias and the relatively smaller n past the age 70 interval. Error density (ΔAge residuals) for metabolomics and proteomics clocks **(F)**. Both clocks show approximately Gaussian residual distributions, with metabolomics displaying a narrower spread and less skew, consistent with lower overall prediction error. Predicted versus chronological age for the merged proteomics + metabolomics model (pre-calibration) **(G)**. The composite model shows strong correlation with observed age (R = 0.801; R² = 0.641), with color-coded residuals illustrating systematic underprediction in younger individuals and overprediction in older individuals. Feature importance (top 25 coefficients) from the metabolomics + proteomics elastic net model **(H)**. Key age-predictive features include complement/coagulation factors, lipid species, redox metabolites (glutathione derivatives), amino acid catabolites (phenylacetyl-glutamine, kynurenine), and xenobiotics. Albumin decreases had the strongest age-related beta-effect in the elastic net model.

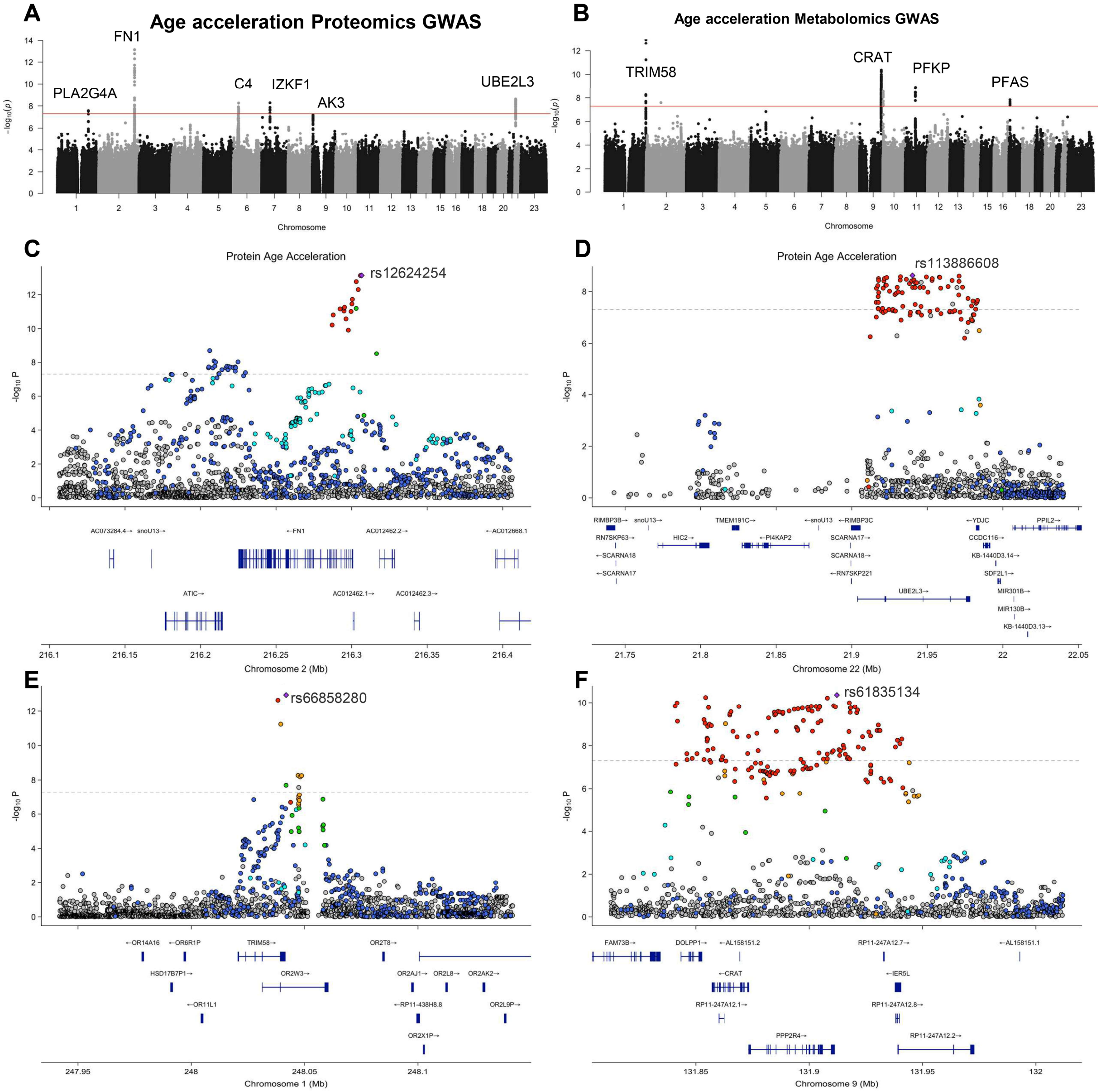

**Supplementary Figure 7. Fine-mapping of genetic loci associated with proteomic and metabolomic age acceleration.** Manhattan plot of the proteomics age-acceleration GWAS, identifying genome-wide significant loci at FN1 (chromosome 2), C4 / IKZF1 (chromosome 7), AK3 (chromosome 9), UBE2L3 (chromosome 22), and PLA2G4A (chromosome 1) **(A)**. These loci suggest genetic determinants of RBC proteomic aging through pathways in extracellular matrix remodeling, immune regulation, mitochondrial metabolism, and ubiquitination. Manhattan plot of the metabolomics age-acceleration GWAS, revealing significant associations at CRAT (chromosome 9), TRIM58 (chromosome 1), PFKP (chromosome 10), and PFAS (chromosome 3) **(B)**. These loci highlight genetic influences on metabolic aging via fatty-acid oxidation, glycolysis, and one-carbon metabolism. Locus zoom highlighting regional association plots for the FN1 locus (rs12624254), strongly associated with proteomic ΔAge. FN1 encodes fibronectin, a component of extracellular matrix and vascular biology, emphasizing RBC–vascular interactions in aging. Regional association plot for the C4 / IKZF1 locus, with lead SNP rs113886608 marking a dense cluster of immune-regulatory variants associated with proteomic age acceleration. The locus encompasses C4A/B, IKZF1, and complement-related genes **(D)**. Fine-mapping of the TRIM58 locus associated with metabolomic ΔAge. The top SNP rs6685280 lies within a regulatory region affecting erythropoiesis-linked genes (TRIM58, GPRC6A, FBXO7), suggesting transcriptional control of metabolic aging **(E)**. Regional association plot for the CRAT locus (chromosome 9), identifying rs61835134 as the lead SNP for metabolomic age acceleration **(F)**. CRAT encodes carnitine O-acetyltransferase, supporting a mechanistic role for fatty acid oxidation and acyl-carnitine metabolism in regulating biological age.

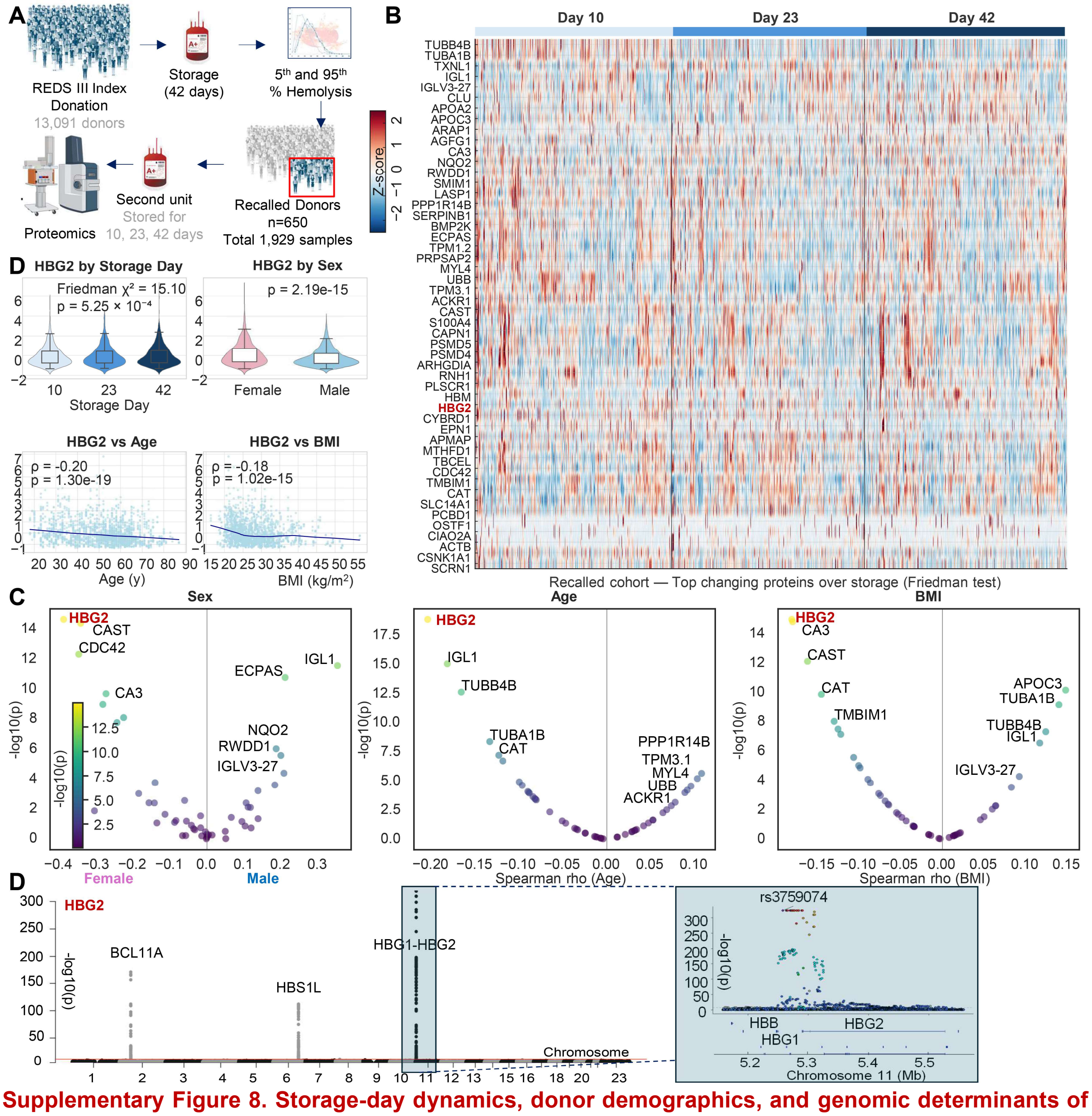

**Supplementary Figure 8. Storage-day dynamics, donor demographics, and genomic determinants of HBG2 abundance in the recalled donor cohort.** Experimental design for the recalled-donor proteomics study **(A)**. From a subset of the REDS III cohort (13,091 donors) – those who ranked <5th or >95th percentile for hemolysis traits or otherwise identified as high frequency donors - a second RBC unit was stored for up to 42 days and sampled on Day 10, Day 23, and Day 42 for proteomics analysis. Proteomic profiling (n = 1,929 samples from 650 donors) was used to quantify storage-dependent changes and donor-specific variation in globin composition—including HBG2, the fetal γ-globin subunit. Heatmap of the top proteins showing significant changes across storage (Friedman test) **(B)**. Proteins involved in cytoskeletal remodeling, proteasomal degradation, redox metabolism, and translation exhibit strong temporal shifts from Day 10 to Day 42. HBG2 – the most significant variable as a function of storage - displays a characteristic pattern of decreasing abundance over storage **(C)**. Volcano plots of proteomics associations to demographic identifies HBG2 as the most strongly influenced variable by Sex, Age and BMI in the recalled donor cohort (higher in younger females with low BMI) **(D)**. GWAS of HBG2 abundance identifies a single highly significant locus on chromosome 11, corresponding to the β-globin gene cluster **(E)**. Locus zoom of the lead SNP (rs3759074) maps in the intergenic region between HBG1 and HBG2, consistent with known regulatory haplotypes modulating fetal hemoglobin persistence. Regional association plots highlight the tight linkage disequilibrium around the globin cluster, with a secondary genome-wide-adjusted significant locus detected on chromosome 2, on the region coding for BCL11, key transcription factor involved in fetal to mature hemoglobin switch.

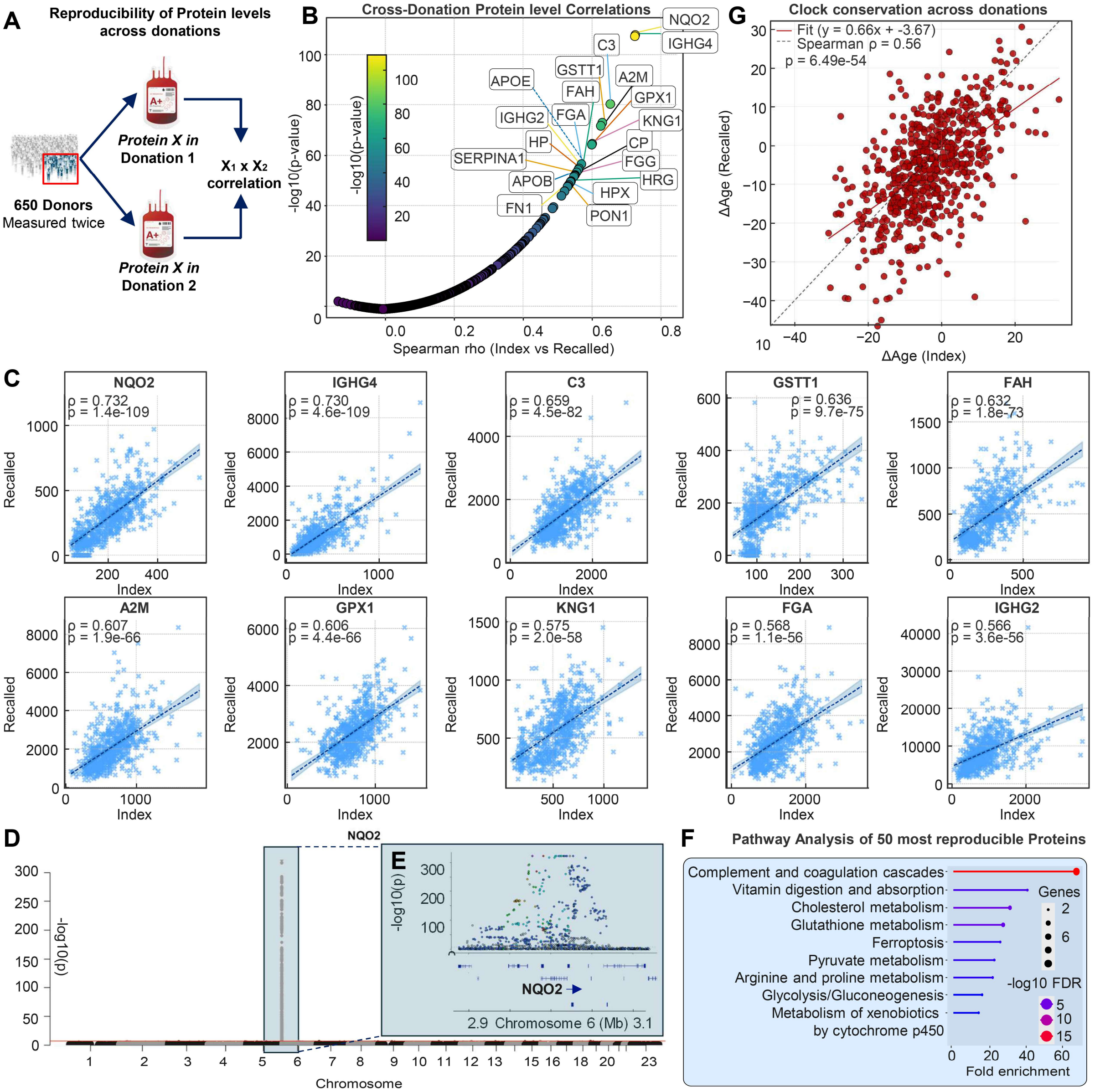

**Supplementary Figure 9. High reproducibility of RBC proteomic measurements across two independent donations from donors enrolled in both Index and Recalled phases of the REDS study is explained by genetic traits.** Overview of the reproducibility analysis **(A)**. For each of the 650 recalled donors, the abundance of each protein was measured twice: once at the end of storage (day 42) for the original index donation and again at day 42 for RBCs derived from units from a second donation collected 6-18 months after enrollment of the Index cohort. Reproducibility was quantified by correlating the paired abundance of each protein across the two independent draws from the same donor. Cross-donation correlations for all quantified proteins. Each point represents one protein, plotted by its Spearman correlation between index and recalled donations (x-axis) and statistical significance (–log10 p, y-axis) **(B)**. A right-skewed with almost all proteins exhibiting positive correlations cross-donations, and some showing exceptionally high reproducibility - including C3, APOE, GSTT1, FAH, FGA, HPX, KNG1, GPX1, and immunoglobulins (IGHG2, IGHG4) - highlighting the stability of RBC proteomic signatures over time within donors (scatter plots in **(C)** display strong linear relationships across donations (ρ ≈ 0.60–0.75; $p < 10^{-50}$). Proteomics quantitative trait loci (pQTL) analysis for the top reproducible protein across donations, NQO2, identifies a single significant cis-QTL locus on chromosome 6 **(D)**, driven entirely by the NQO2 coding region **(E)**. No other loci surpass genome-wide significance, indicating that reproducibility is broadly genetically determined, reflecting intra-donor biological stability of RBC proteome signatures in a short time span. Pathway enrichment of the 50 most reproducible proteins, revealing overrepresentation of innate immune and metabolic pathways, including complement/coagulation, xenobiotic metabolism, vitamin digestion/absorption, cholesterol metabolism, glutathione and redox pathways, ferroptosis, and glycolysis/gluconeogenesis. These pathways represent core, donor-stable components of RBC biology **(F)**. Like proteins, ΔAge calculated off the multi-omic aging clock trained on the REDS Recalled Protoemics data strongly correlated with ΔAge from the Index donation (ρ = 0.66, p = $6.5\times10^{-54}$) **(G)**. The positive slope demonstrates that molecular aging signatures embedded in the RBC proteome are highly stable within individuals over repeated donations.

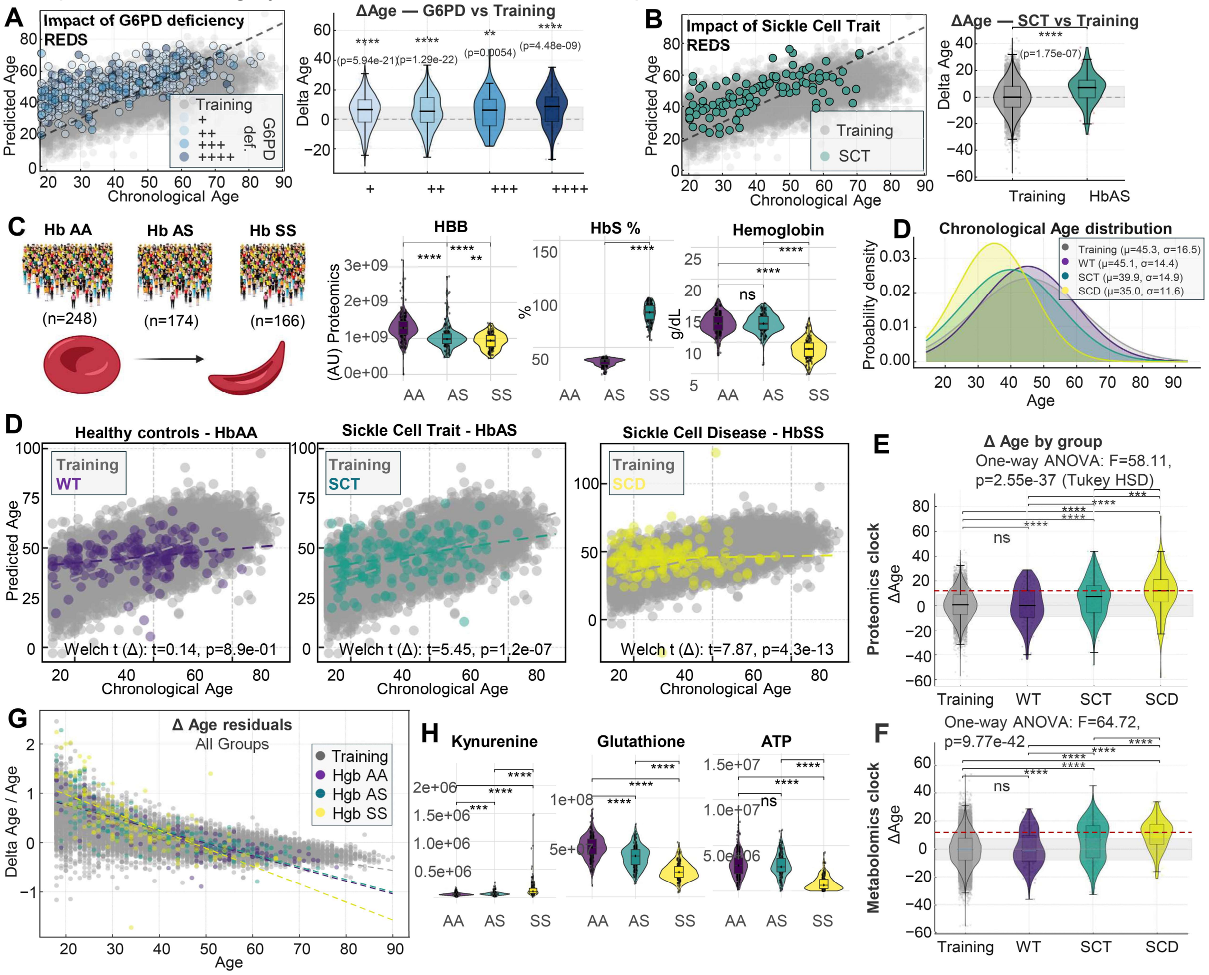

**Supplementary Figure 10. Genetic traits linked to hemolytic disorders accelerate the aging clocks. Effects of G6PD deficiency, sickle cell trait, and sickle cell disease on biological age, proteomic remodeling, and metabolomic aging signatures.** Impact of G6PD deficiency on proteomics-based predicted age in the REDS cohort **(A)**. Left: Predicted vs chronological age for donors with increasing genetic severity of G6PD deficiency ("+" to "++++" – Class I to IV). Right: ΔAge (Predicted – Chronological Age) shows stepwise acceleration of biological age with increasing severity (Kruskal–Wallis $p < 6\times10^{-21}$), indicating that impaired redox homeostasis accelerates multi-omic aging signatures. Impact of sickle cell trait (HbAS) in the REDS Index cohort. Left: Predicted vs chronological age for HbAS donors relative to G6PD-matched training controls. Right: HbAS donors show significantly accelerated ΔAge ($p = 1.75\times10^{-7}$), consistent with chronic low-grade hemolytic stress and altered redox/erythropoietic tone **(B)**.

In an independent cohort, RBC proteomic and hematologic differences were observed in subjects carrying HbAA, HbAS, and HbSS (n = 248, 174, and 166 respectively) **(C)**. Left: schematic of genotypes. Middle: HBB protein abundance is significantly elevated in HbSS and modestly increased in HbAS. Right: HbS% differs across groups as expected, and hemoglobin concentration is markedly reduced only in HbSS. Predicted age for healthy HbAA, sickle cell trait (HbAS), and sickle cell disease (HbSS) donors relative to the training set **(D)**. HbAA matches training predictions, whereas HbAS and HbSS show progressively greater positive residuals. Welch's tests quantify ΔAge shifts, with HbSS showing the strongest acceleration ($t = 7.87$, $p = 4.3\times10^{-13}$). ΔAge (proteomics or metabolomics clock) by genotype **(E-F)** show stepwise acceleration by genotype-associated hemolytic susceptibility. ΔAge/Chronological Age vs chronological age across all groups **(G)**. Both sickle phenotypes show age-independent elevation in biological aging, whereas controls exhibit only mild age-related drifts. Regression slopes highlight genotype-dependent aging acceleration. Redox and energy Metabolite abundance differences among HbAA, HbAS, and HbSS donors for key redox- and stress-linked metabolites **(H)**.

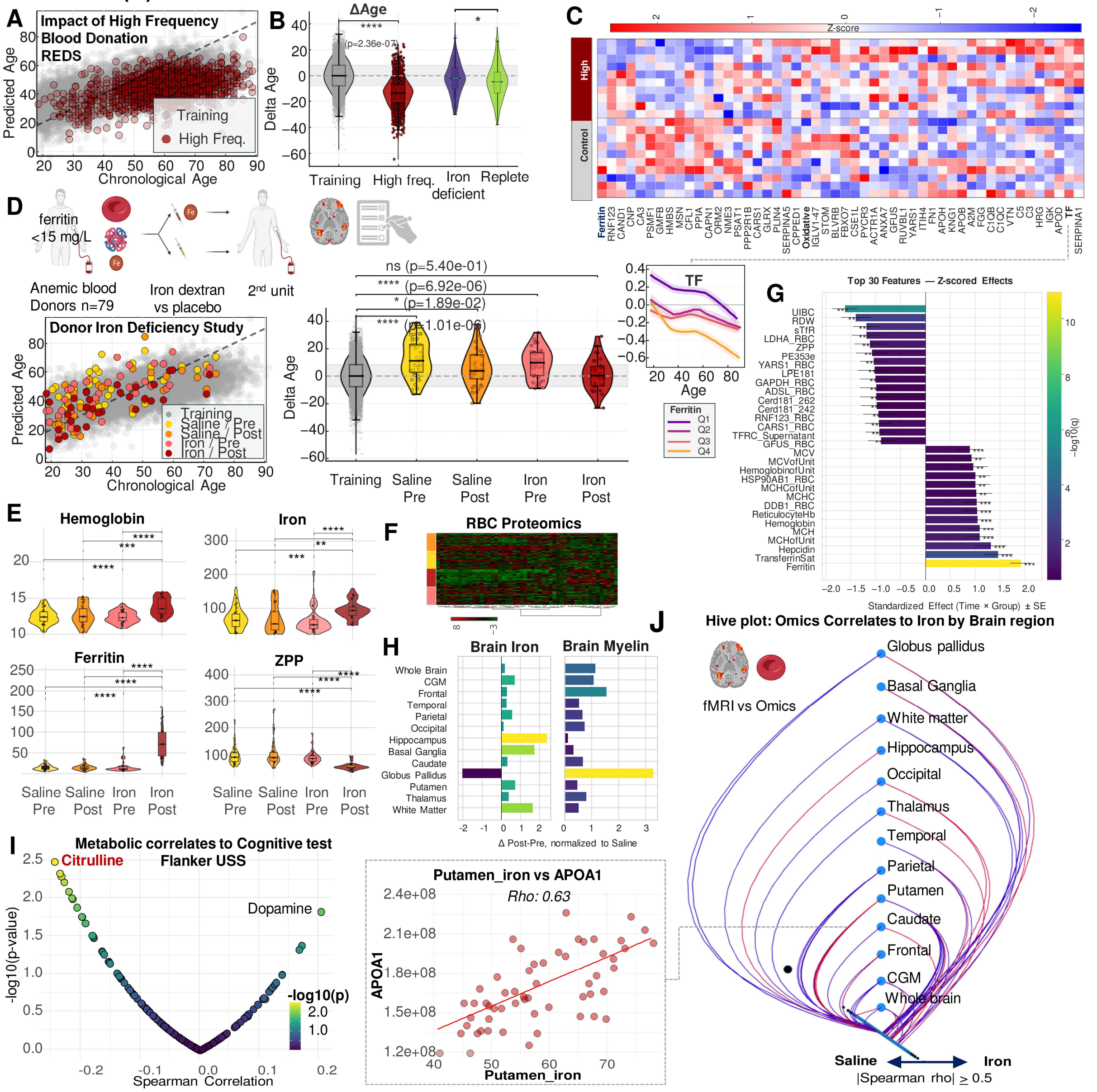

**Supplementary Figure 11. High-frequency blood donation slows down aging clocks, as long as donor do not develop iron deficiency. Iron repletion in iron deficient donors in the DIDS cohort normalizes systemic iron homeostasis, RBC proteomics changes, brain iron/myelin signatures, cognitive assays and resets the aging clock.** High-frequency blood donation slows down biological aging **(A)**. Left: Predicted vs chronological age for REDS donors with high donation frequency compared to the training cohort. Right: ΔAge values show significantly decreased biological age in high-frequency donors in the absence of anemia (ferritin >15 mg/L) ($p = 2.4\times10^{-07}$) in REDS RBC Omics **(A)**. Age trajectories of transferrin (TF) stratified by ferritin quartiles **(B)**. Donors with lower ferritin (Q1–Q2) show higher TF across the lifespan, reflecting compensatory upregulation of iron-transport pathways. Heatmap of proteomic features linked to high frequency

donor status (>3 donations in the past 12 months), showing strong associations with globins, iron-transport proteins, redox enzymes, and stress response pathways. Lower ferritin is associated with upregulation of iron-mobilizing proteins and shifts in RBC metabolic state, consistent with high frequency donation showing trajectories towards depletion of iron pools **(C)**.
Prospective donor iron deficiency study (DIDS) **(D).** Left: Schematic of the trial design—iron-deficient donors (ferritin <15 ng/mL, n = 79) were randomized to receive I.V. supplementation of iron dextran or placebo (saline), followed by a second unit collection. Middle: Predicted age vs chronological age for Training, Saline (Pre/Post), and Iron (Pre/Post) groups. Right: ΔAge values showed significantly faster age acceleration in iron-deficient donors ($p = 6.9 \times 10^{-6}$), and substantial reduction – back to control levels - after iron repletion but not saline (not significant). **(E).** Hematologic and iron biomarkers are significantly impacted by iron repletion, but not saline, with significant ($p<0.0001$) increases in hemoglobin, iron and ferritin, and decreases in zinc protoporphyrin (ZPP) after iron supplementation, while saline has no effect. Heatmap of RBC proteomic changes across saline vs iron groups (Pre/Post) **(F)**. Iron supplementation induces broad remodeling of metabolic, redox, membrane, and proteostasis pathways. Barplot of the top 30 features (proteomics, CBC, iron parameters) differentiating iron vs saline groups (standardized effect sizes) **(G)**. Iron supplementation reverses depletion of structural, glycolytic, and mitochondrial proteins and reduces stress markers, including LDHA, RBC membrane proteins, redox enzymes, regulators of heme synthesis and iron-dependent aminoacyl-tRNA synthesizing enzymes (CARS1, etc). Functional Magnetic Resonance Imaging (fMRI)-**derived brain iron and brain myelin** changes after iron supplementation (post–pre normalized to saline) **(H)**. Regions such **as globus pallidus, putamen, and caudate** - sites of high iron utilization- show measurable increases in iron and myelin metrics, consistent with systemic-to-brain iron redistribution (consistent with the analyses in Hod et al. JCI Insight 2025; https://doi.org/10.1172/jci.insight.194442. PMID: 41118254). Metabolite correlates to cognitve test (cognitive performance by Flanker USS) **(I)** reveal significant associations for aging clock top predictors citrulline and neurotransmitter metabolites linked to neurodegeneration (dopamine), suggesting metabolic signatures of iron status influence cognitive domains. Hive plot showing omics-brain regional iron correlations (Spearman $\rho \geq 0.5$) **(J)**. Proteomic and metabolomic features strongly correlate with MRI-derived iron in deep gray structures (globus pallidus, basal ganglia, putamen) and white matter. Right: Scatterplot showing a representative strong correlation between Putamen iron and APOA1 ($\rho = 0.63$) - the top variable in the elastic net proteomics aging clock in REDS, highlighting RBC molecular signatures linked to molecular aging and neuroradiologic coupling of iron biology and cognitive function.

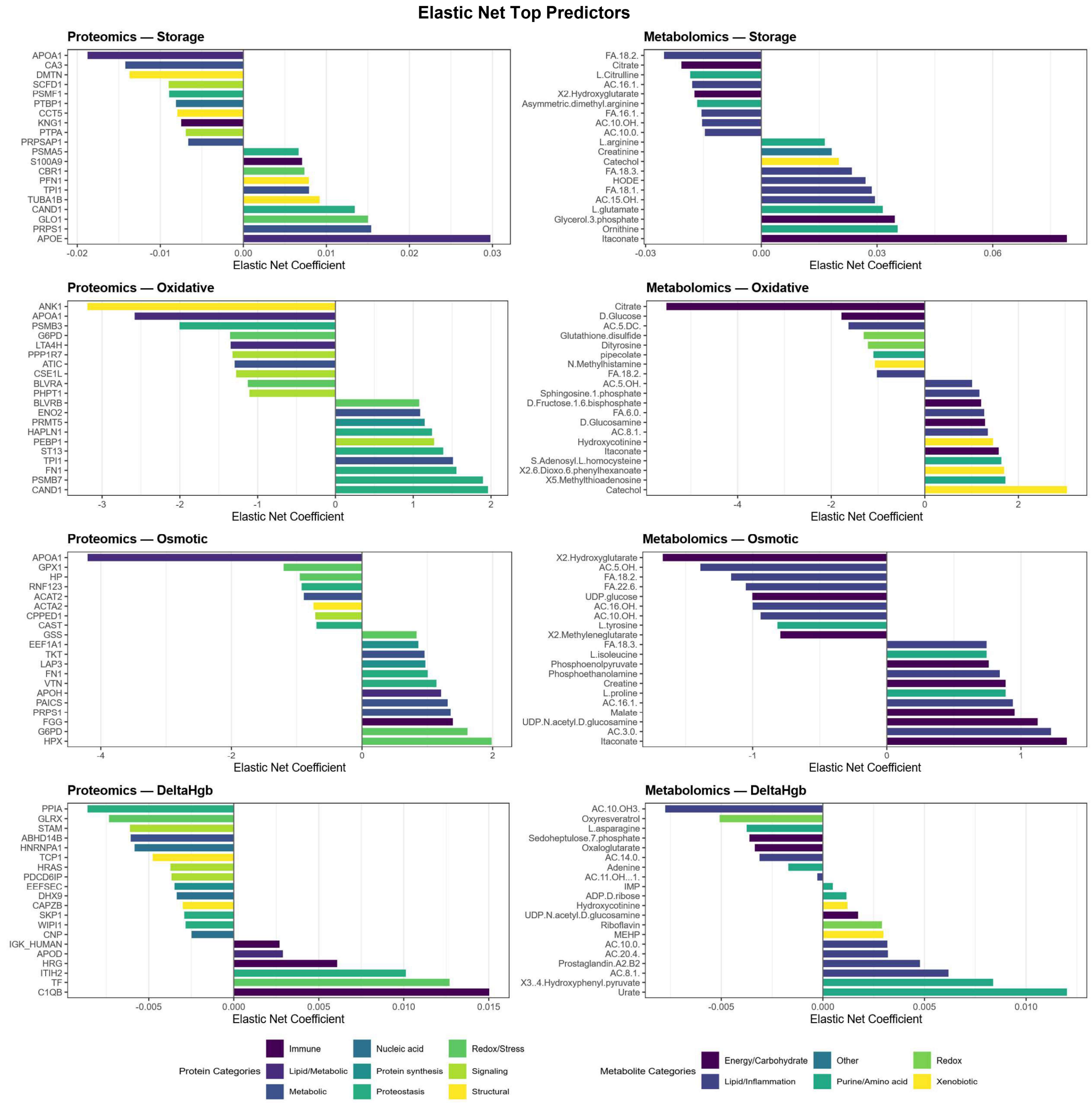

**Supplementary Figure 12. Elastic net models identify top proteomic and metabolomic predictors of storage, oxidative, and osmotic hemolysis, as well as ΔHgb.** Elastic net regression was applied independently to proteomics and metabolomics datasets to identify molecular predictors of four hemolysis-related phenotypes: storage hemolysis, oxidative hemolysis, osmotic hemolysis, and ΔHgb (hemoglobin increment). Features shown represent the strongest positive and negative coefficients, grouped by biological category.

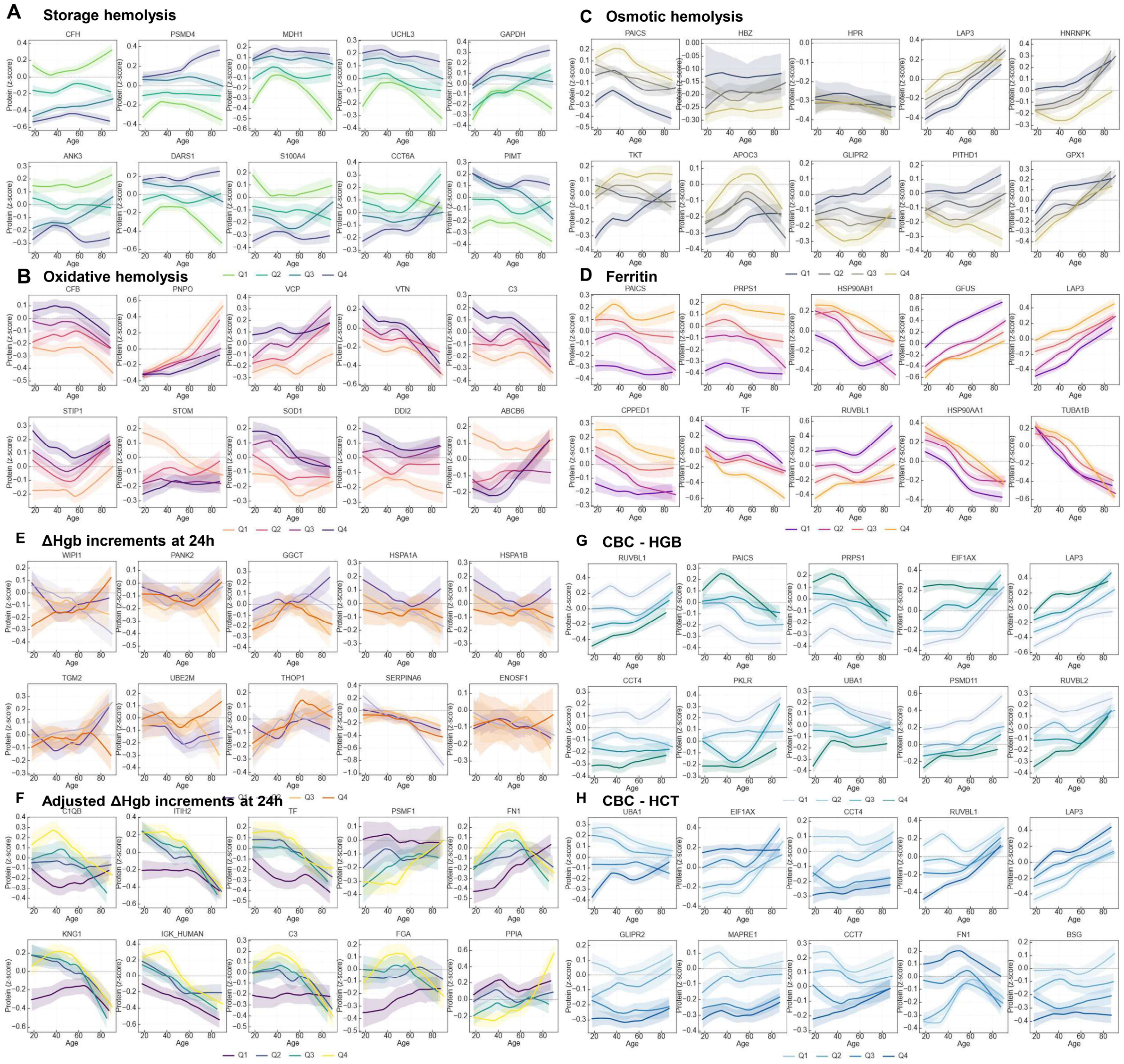

**Supplementary Figure 13. Age-trajectories of top proteomic predictors of hemolysis phenotypes, iron status, hemoglobin increments, and CBC indices.** Age-dependent trajectories (LOESS fits ± 95% CI) are shown for proteins stratified by quartiles of the corresponding phenotype (Q1 = lowest, Q4 = highest). Each panel displays the proteins with the strongest elastic-net effect sizes for the indicated trait, illustrating how molecular aging patterns interact with donor physiology. Results are shown for storage, oxidative, osmotic hemolysis, adjusted hemoglobin increments at 24h from single unit transfusion, complete blood count (CBC) parameters including hemoglobin (HGB) and hematocrit (HCT) **(A-H)**.

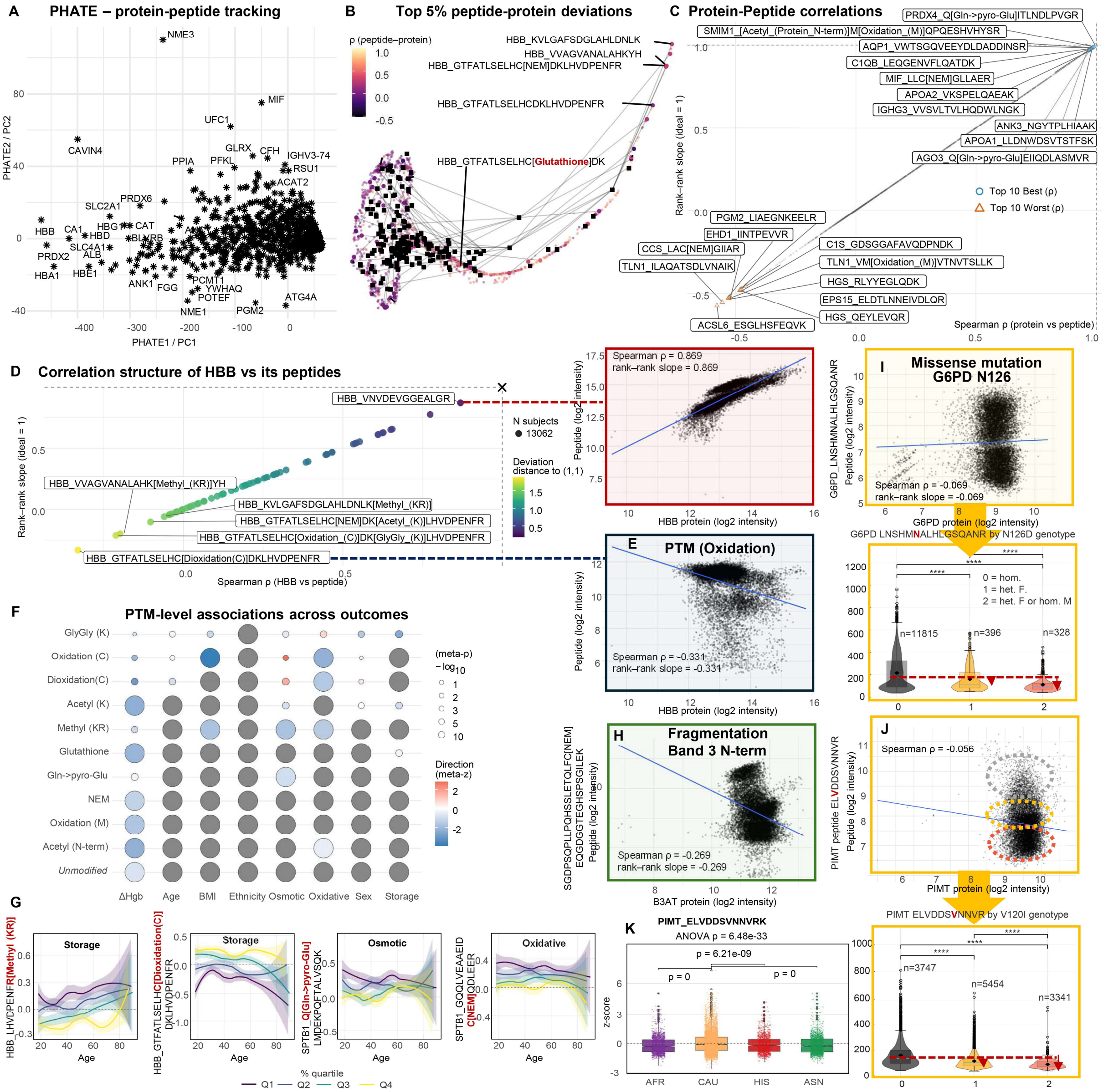

**Supplementary Figure 14. Peptide-level resolution of RBC proteome variation reveals PTM biology, peptide–protein discordance, and genetic determinants of RBC aging and hemolysis.** PHATE embedding of peptide abundances for all quantified proteins highlights peptide-level heterogeneity within individual proteins, including proteins with substantial internal structural diversity (e.g., NME3, MIF, PRDX5, ACAT2, HBB) **(A)**. Protein centroids (labeled) are surrounded by clouds of constituent peptides whose positions reflect PTMs, cleavage patterns, and quantitative variability**.** Top 5% most deviant peptides from parent proteins **(B)**. Each peptide is plotted by its deviation from the expected peptide–protein relationship (residual of a robust regression). Correlation structure of peptide–protein relationships **(C)**. Scatterplots identify peptides with the best (top 10) and worst (bottom 10) peptide–protein concordance across all donors. Poorly correlated peptides cluster in proteins known for extensive PTM regulation (e.g., PRDX1, PGM2, TLN1, IGLV3), whereas highly correlated peptides belong to more structurally stable proteins. Peptides from hemoglobin β (HBB) are strongly enriched among high-deviation events **(D)**, including peptides modified by glutathionylation, oxidation, acetylation, or NEM (thioalkylation), highlighting extensive PTM heterogeneity **(D)**. A network representation orders HBB peptides by pairwise rank-based correlations, revealing distinct clusters corresponding to specific modifications (oxidation, methylation, glutathionylation, N-terminal truncation). Right: example peptide shows high correlation with total HBB (Spearman ρ = 0.869 – right panel **(D)**). Loss of correlation between parent protein and peptide abundance is either explained by protein post-translational modification **(E)** (a full list and association to demographics and hemolysis in **(F)**, with selected highlights in

**(G)**, fragmentation (e.g., SLC4A1 – band 3 N-terminus **(H)**) or missense mutations **(I-J)**. Impact of G6PD N126D missense mutation on PTM abundance **(I)**. Carriers of the G6PD N126D variant show increased abundance of oxidative HBB peptides (G6PD deficiency–linked oxidative load). Violin plots show significant genotype-specific enrichment, consistent with metabolic vulnerability. Carriers of the PIMT V120I missense mutation show significant loss of correlations between peptide-parent protein abundances, consistent with allele distributions for the related SNP rs4816 (chr9), highlighting a genetically influenced RBC peptide level abudnances. Since this allele is prevalent in donors of Asian descent (MAF >60%), ethnic differences in PIMT V120-containing peptpide abundance was lower in donors carrying these alleles **(K).** Collectively, peptide-level profiling reveals previously unresolvable biochemical heterogeneity across the RBC proteome and exposes PTM-, protein fragmentation/cleavage-, and fine mapping of genotype-dependent variation that cannot be inferred from protein-level analyses alone.

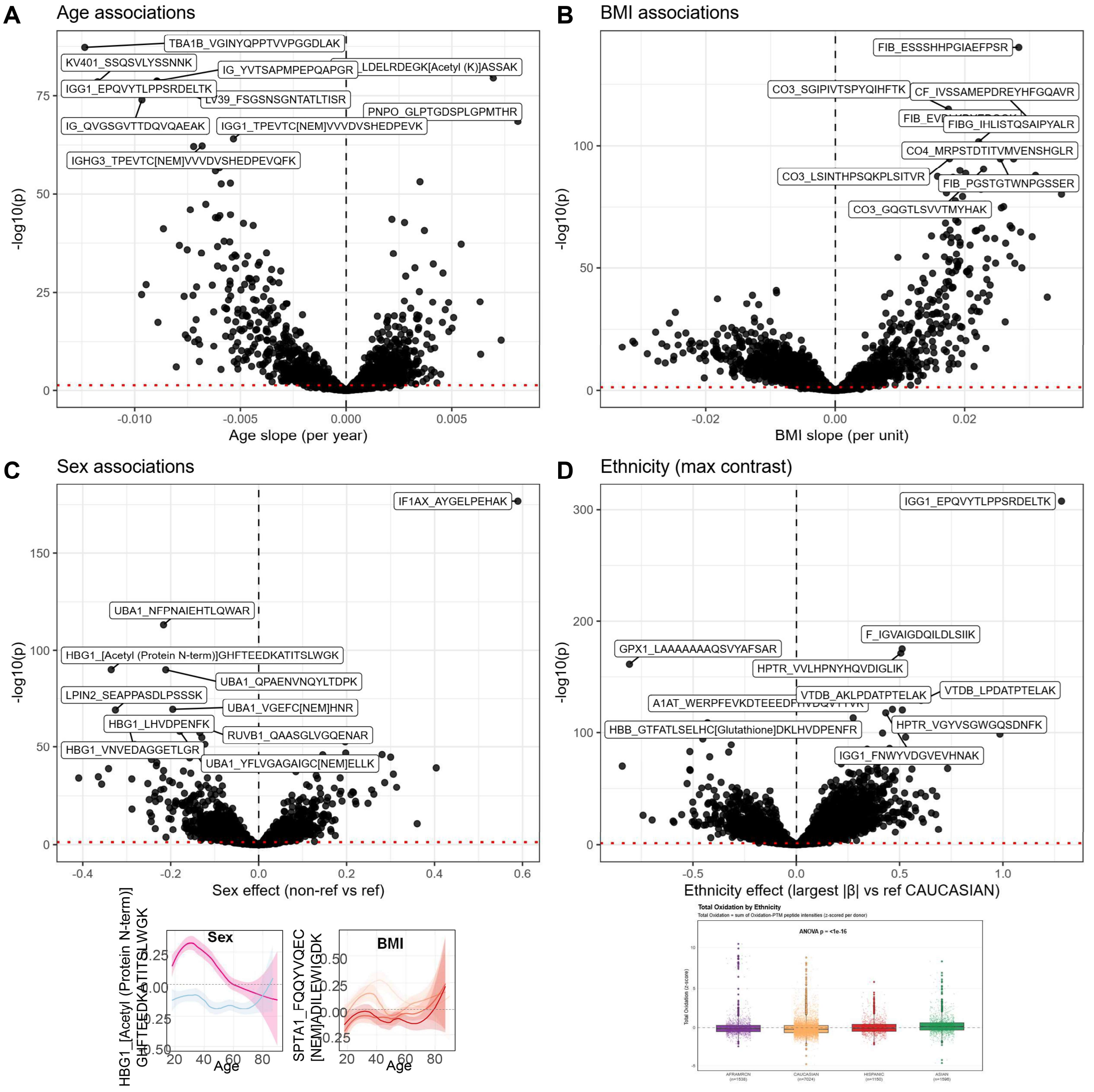

**Supplementary Figure 15. Peptide-level associations with age (A), BMI (B), sex (C), and ethnicity (D) reveal demographic influences on PTMs, fragmentation, and RBC proteome architecture.**

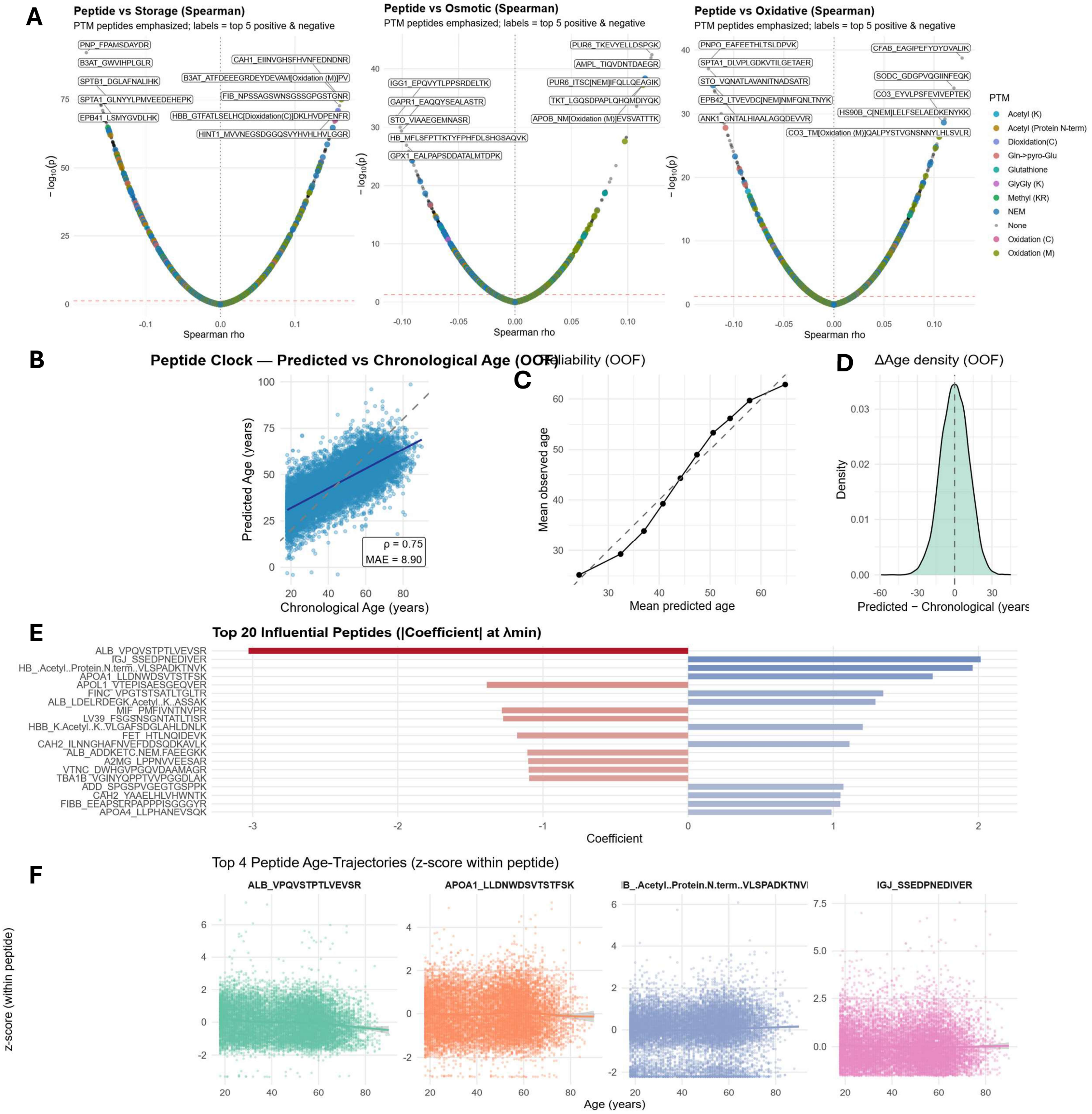

**Supplementary Figure 16. A peptide-based aging clock reveals PTM-level aging biology and identifies key peptides driving predicted biological age.** Spearman correlations between peptide levels and hemolysis **(A)**. Peptide-based aging clock performance **(B)**. Predicted age vs chronological age (ρ = 0.75; MAE = 8.90 years) demonstrates robust biological aging signal derived exclusively from peptide-level data **(B)**. Out-of-fold (OOF) predictions confirms that performance reflects true generalization **(C)**. The peptide clock shows strong calibration across most age ranges, with slight underprediction at the highest ages, consistent with known saturation effects in proteome aging **(C)**. Distribution of ΔAge (Predicted – Chronological Age) in **(D)**.
Residual distribution is centered near zero, indicating no systematic bias, with a symmetric density characteristic of a stable aging signature. Top 20 influential peptides (absolute elastic-net coefficients at λmin) are listed in **(E)**. Age-trajectory plots of top four peptide predictors (z-scored within peptide) in **(F)**. Representative peptides—ALB_VPQSTPTLVEVS, APOA1_LLDNWDSVTSTFSK, HBB_Acetyl(Protein N-term)_VLSPADKTNV, and IGJ_SSEEPDNEDVER—show distinct age-dependent trends, demonstrating the clock's sensitivity to PTMs, cleavage events, and immunoglobulin turnover.
